# Decision-Making Time Cells in Hippocampal Dorsal CA1

**DOI:** 10.1101/2023.10.01.560382

**Authors:** M. Ma, F. Simoes de Souza, G.L. Futia, S.R. Anderson, J. Riguero, D. Tollin, A. Gentile-Polese, J.P. Platt, N. Hiratani, E. A. Gibson, D. Restrepo

**Affiliations:** Department of Cell and Developmental Biology, University of Colorado Anschutz Medical Campus, Aurora, CO, 80045, USA; Center for Mathematics, Computation and Cognition, Federal University of ABC, Sao Bernardo do Campo, SP, Brazil; Department of Bioengineering, University of Colorado Anschutz Medical Campus, Aurora, CO, 80045, USA; Department of Physiology and Biophysics, University of Colorado Anschutz Medical Campus, Aurora, CO, 80045, USA; Neuroscience Graduate Program, University of Colorado Anschutz Medical Campus, Aurora, CO 80045, USA; Department of Neurosurgery, University of Colorado Anschutz Medical Campus, Aurora, CO, 80045, USA; Department of Neuroscience, Washington University, St. Louis, MO 63110, USA

## Abstract

Sequential neural dynamics encoded by “time cells” play a crucial role in hippocampal function. However, the role of hippocampal sequential neural dynamics in associative learning is an open question. In this manuscript, we used two-photon Ca^2+^ imaging of dorsal CA1 pyramidal neurons in head-fixed mice performing a go-no-go associative learning task. We found that pyramidal cells responded differentially to the rewarded or unrewarded stimuli. The stimuli were decoded accurately from the activity of the neuronal ensemble, and accuracy increased substantially as the animal learned to differentiate the stimuli. Decoding the stimulus from individual pyramidal cells that responded differentially revealed that decision-making took place at discrete times after stimulus presentation. Lick prediction decoded from the ensemble activity of cells in dCA1 correlated linearly with lick behavior indicating that sequential activity of pyramidal cells in dCA1 constitutes a temporal memory map used for decision-making in associative learning.

## INTRODUCTION

The hippocampus provides real-time encoding and retrieval of detailed context memories enabling reactions to a constantly changing environment (Wiltgen et al., 2010). Diverse and distributed neuronal activity encodes external features such as contextually referenced space and time and sensory stimuli, as well as features influenced by the animal’s behavior such as speed and direction of motion (Buzsáki and Llinás, 2017; Buzsaki and Moser, 2013; Mau et al., 2018; Stefanini et al., 2020). Plastic circuits involved in associative learning mediate this demanding task (Gruart et al., 2015; Neves et al., 2008). Here we address the neural representation of associative learning in pyramidal neurons of dorsal CA1 (dCA1), a hippocampal area involved in spatial learning and working memory tasks.

The involvement of dCA1 in associative learning of stimulus discrimination in head-fixed animals is still unclear (Biane et al., 2023; Li et al., 2017; Moser et al., 1993; Rangel et al., 2016; Strange et al., 2014; Taxidis et al., 2020). Learning in animals undergoing an odorant discrimination associative learning go-no go task changes the responsiveness of dCA1 pyramidal cells to sensory stimuli (Biane *et al*., 2023; Li *et al*., 2017). Li and co-workers showed that for mice engaged in an odor discrimination go-no go task dCA1 pyramidal cells receiving connections conveying olfactory information from the lateral entorhinal cortex (LEC) develop more selective spiking responses to odor cues as they learn to discriminate odorants and they showed that optogenetic inactivation of the LEC to dCA1 connections slows learning in this task (Li *et al*., 2017). In addition, Biane and colleagues find a learning-related increase in the proportion of cells responsive to the rewarded (S+) odorant, but not to the unrewarded (S-) odorant, suggesting that stimulus representations in dCA1 are sensitive to perceived value. However, learning did not alter decoding accuracy that was already high before training in dCA1 (Biane *et al*., 2023). Notably, temporal patterning of dCA1 neural activity plays a crucial role in odorant working memory tasks (MacDonald et al., 2013; Taxidis *et al*., 2020) and has been postulated to play a role in organizing memories (Eichenbaum, 2017) raising the question whether it contributes to the neural mechanism for go-no go associative learning in this brain region. Hippocampal “time cells” store memory of the temporal order of events and working memory and signal changes in the temporal context (Eichenbaum, 2017; Shimbo et al.; Taxidis *et al*., 2020). It has been hypothesized that the precise temporally structured activity of neurons make perceptually related responses coherent in time (Singer and Lazar, 2016), but it is unknown whether there is a time-tiled representation of *stimulus divergence* in dCA1 that could contribute through sequential neural dynamics to behavioral responses reflecting decision-making in associative learning.

We use two-photon calcium imaging to evaluate ensemble decoding of stimuli from temporally patterned neural activity in dCA1 in mice engaged in an olfactory go-no go task where they receive a water reward after licking on a spout when presented with the S+ odorant and do not receive a water reward for the S-odorant (Ma et al., 2020) (Figure 1A). We characterized stimulus divergence of calcium responses of different cells (regions of interest, ROIs) from the calcium video by multivariate robust estimation (Hakan et al., 2021) and we assessed changes in neuronal activity after switching rewarded with unrewarded stimuli to determine whether the cells respond to the identity of the stimulus as opposed to its valence (is the stimulus rewarded?). Finally, we assessed the accuracy of decoding the stimuli with different subsets of cells and determined whether the onset of stimulus decoding reflecting divergence of stimulus responses was time-tiled.

**Figure 1.**
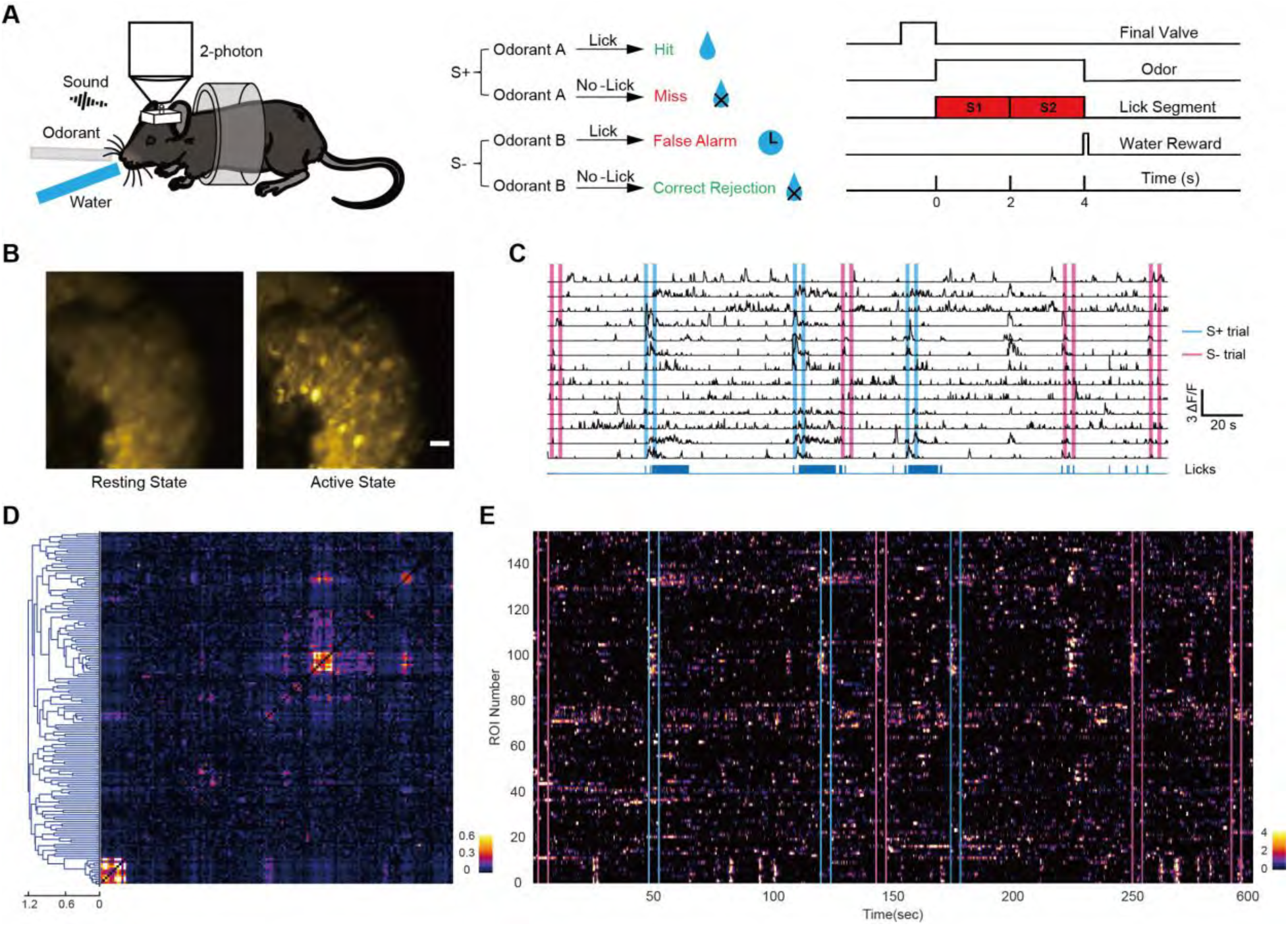
Two-photon Ca^2+^ imaging of pyramidal cells in CA1 in head-fixed mice undergoing the go-no go associative learning task. (A) Go-no go task. Left: Two-photon imaging of a head-fixed mouse responding to odorants by licking on a water spout in response to the rewarded odorant in the go-no go task. Center: Scoring of decision making. Right: time course for the trial. For water reward in Hit trials, the animal must lick at least once in each of the two 2 second lick segments. (B) Two-photon microscopy images of GCaMP6f fluorescence recorded from CA1 pyramidal cells through a GRIN lens in a proficient Thy1-GCaMP6f mouse engaged in the go-no go task. Left: Activity at rest. Right: Activity during the rewarded odorant trial. Video 1 shows fluorescence changes in this group of cells. (C) ΔF/F traces are shown for a subset of the regions of interest (ROIs) for the figure in B. The magenta vertical lines are the on and off times for the odorant in the unrewarded (S-) odorant trials and the cyan vertical lines are on and off times for the rewarded (S+) odorant. The blue lines at the bottom are lick recordings. (D) Cross-correlation and hierarchical clustering of the ΔF/F traces for 153 ROIs for the entire 20 min session corresponding to B and C shows substantial heterogeneity in the calcium responses. (E) Pseudocolor plot of the time course for ΔF/F for the 153 hierarchically clustered ROIs. Figure S1 shows the time courses for S+ and S-trials for the mean ΔF/F traces, lick fraction and a p-value quantifying the difference between S+ and S-licks.

## RESULTS

### Two-Photon Calcium Imaging in Dorsal CA1 in Head-Fixed Mice Undergoing a Go-No Go Olfactory Associative Learning Task

Calcium imaging was performed through a GRIN lens in adult Thy1-GCaMP6f mice (n=4) expressing the calcium indicator GCaMP6f in pyramidal neurons in dCA1 (Figure 1A, Figure S1). Water-deprived mice started the trial by licking on the water spout. One of two odorants was delivered in random order 1-1.5 sec after the start of the trial and the odorant was presented for 4 sec. Mice were trained to respond to the rewarded odorant (S+) by licking at least once during two 2 sec response areas to obtain a water reward. The water reward was not delivered regardless of licking when the unrewarded odorant (S-) was presented. Figure S1A shows this session’s differential lick response to S+ vs. S-. Two-photon calcium imaging movies were denoised (Li et al., 2023), motion-corrected (Pnevmatikakis and Giovannucci, 2017) and time-binned traces of nonnegative changes in fluorescence intensity (ΔF/F) were obtained for multiple ROIs using EXTRACT, an algorithm based on robust estimation theory (Hakan *et al*., 2021). Figures 1B,C,E and Video 1 show calcium imaging data from a 20 min session where EXTRACT found 153 ROIs in dCA1 for a mouse proficient in the go-no go task (proficient stage is defined as percent correct >= 80%). As expected the fluorescence traces displayed calcium spikes whose timing was heterogeneous when evaluated by the cross-correlation between the traces of all ROIs (Figure 1D). In this session a subset of the ROIs responded differentially during S+ vs. S-odorant trials (Figures 1C,E). Consistent with the heterogeneous response there was little difference between S+ and S-trials when the ΔF/F was averaged over all ROIs (Figure S1C). However, there was an increase in mean ΔF/F following trial start. These results suggest the existence of a complex time-structure underlying the calcium responses from a population of dCA1 neurons in the go-no go task.

### Mice Learn to Respond Differentially to the Rewarded and Unrewarded Odorants in the Go-No Go Task

In the first training session the percent correct response to the odorants was between 45% and 65% correct for three of four mice (Figure S2 A-C)(S+: 1% heptanal, HEP, S-: mineral oil, MO) Percent correct was calculated as 100*(hits + CR)/number of trials, we name the session as “learning stage” when percent correct behavior is between 45% and 65% correct. We then trained the mice in 3-6 training sessions per day. As shown in Figure S2 A-D the mice gradually reached the proficient stage (percent correct behavior >80%). The fourth mouse became proficient in the first session (Figure S2D).

We recorded from a total of 18 learning stage sessions in three mice and 66 proficient sessions in four mice. Once the mice had stable proficient performance the odorants were reversed (S+: mineral oil, MO, S-: 1% HEP) to investigate whether dCA1 calcium responses encode for the odorant’s identity or valence. The percent correct behavior dropped immediately below 50% and gradually recovered to proficient (>80%) (Figure S2A-D). We recorded from a total of 25 forward proficient stage sessions and 41 reversed proficient stage sessions in four mice).

The go-no go task used in this study is an olfactory cued task where the animal starts a trial by licking on the water spout. At the start of the trial a diverter valve turns on directing the 2 lt/min air flow away from the nose cone towards an exhaust line (Figure 1A) and one of two odorant valves (S+ or S-) opens allowing the odorant, delivered at 50 ml/min, to equilibrate with the 2 lt/min background air flow. 1-1.5 sec after trial start the diverter valve brings the air back to the nose cone resulting in a sharp increase in odorant concentration providing the mouse with an odorant cue with an onset ∼100 msec after reset of the diverter valve (Figure S3A). In the rest of the manuscript the time when the diverter valve turns back to the odorant cone is time zero. Finally, turning on the odorant valves at the start of the trial delivers a clicking sound that can be used to decode the stimuli (see Methods and Figure S3 B-D). However, mice do not cue on the click as shown by the fact that when the odorant is removed (but the odor valve still clicks) the performance drops to 50% (Figure S2 E-G, consistent with previous studies with the same olfactometer (Slotnick and Restrepo, 2005)).

Figure S4 illustrates lick behavior in learning and proficient stages for the four mice in this study. Figures in S4A show the per trial time course for mean lick fraction for S+ and S-trials calculated for all sessions (18 learning stage and 66 proficient stage, 4 mice). The lick fraction is defined per time point as the fraction of trials within a session when the lick is detected and ranges from zero (no licks detected) to one (a lick was detected in every trial). Mice start the trial by licking on the water spout 1 to 1.5 sec before odorant delivery evidenced by an increase in mean lick fraction before odorant delivery (arrow in Figure S4A right panel). Mice in the learning stage start licking again shortly before the odorant is delivered. The mean lick fraction starts diverging between S+ and S-trials after about one second (Figure S4A, left panel). In order to quantify when the lick behavior differs between S+ and S-we computed a p-value of the difference in S+ vs S-lick fraction using a ranksum test. The left panel in Figure S4B shows that for mice in the learning stage, the p-value for lick divergence decreases monotonically below 0.05 (black dashed line) when the water reward is delivered (first red vertical line). In contrast, for the proficient mice (Figure S4B, right panel) the mean lick fraction diverges between S+ and S-a fraction of a second after odorant delivery. Figure S4C summarizes the mean lick fraction for different time windows within the trial. We find a sharp increase for proficient mice in the difference between S+ and S-trials in lick fraction during the odor period. We did not find a difference in lick fraction between S+ and S-trials during the pre time period (−1.5 to 0 sec) when the animal could use auditory cues. A generalized linear model (GLM) analysis yielded a statistically significant difference for all time windows (pre, odor and reinforcement) vs. baseline time window, for the interaction between learning vs. proficient and the odor or reinforcement windows vs. baseline time window and for the interaction between learning vs. proficient, time window and S+ vs S-for the odor and reinforcement windows (vs. baseline time window) (p<0.001). Still, there was no difference for this interaction for the pre time window (p>0.05), 744 observations, 725 d.f., 4 mice, GLM F-statistic=153, p<0.001 (Table S1). This lick behavior that becomes divergent between S+ and S-trials a fraction of a second after odorant delivery is consistent with a previous study performed with the same olfactometer (Ma *et al*., 2020). Taken together the data on 50% mouse performance in the absence of odorants (Figure S2E-G) and lick behavior (Figure S4) indicate that mice cue on odorants in this multisensory go-no go task.

### n-Dimensional ΔF/F Space Expands before the Animal Starts the Trial and the Dimensionality of Neural Space Increases as the Animal Learns

The highly complex dynamics of the dCA1 network provide a high-dimensional space that could accommodate an immense repertoire of potential states (Figure 2A, left). Presumably, the n-dimensional network dynamics collapses into stimulus specific metastable subregions detectable as manifolds in n-dimensional space (Nieh et al., 2021; Singer and Lazar, 2016)(Figure 2A, right). The manifolds would constrain the trajectory in n-dimensional space in a contextually dependent manner that may be reflected in changes in dimensionality. We proceeded to evaluate changes in n-dimensional neural space where each dimension is the neural activity for each ROI quantified by z-scored ΔF/F calcium changes as a function of time within each trial (zΔF/F).

**Figure 2.**
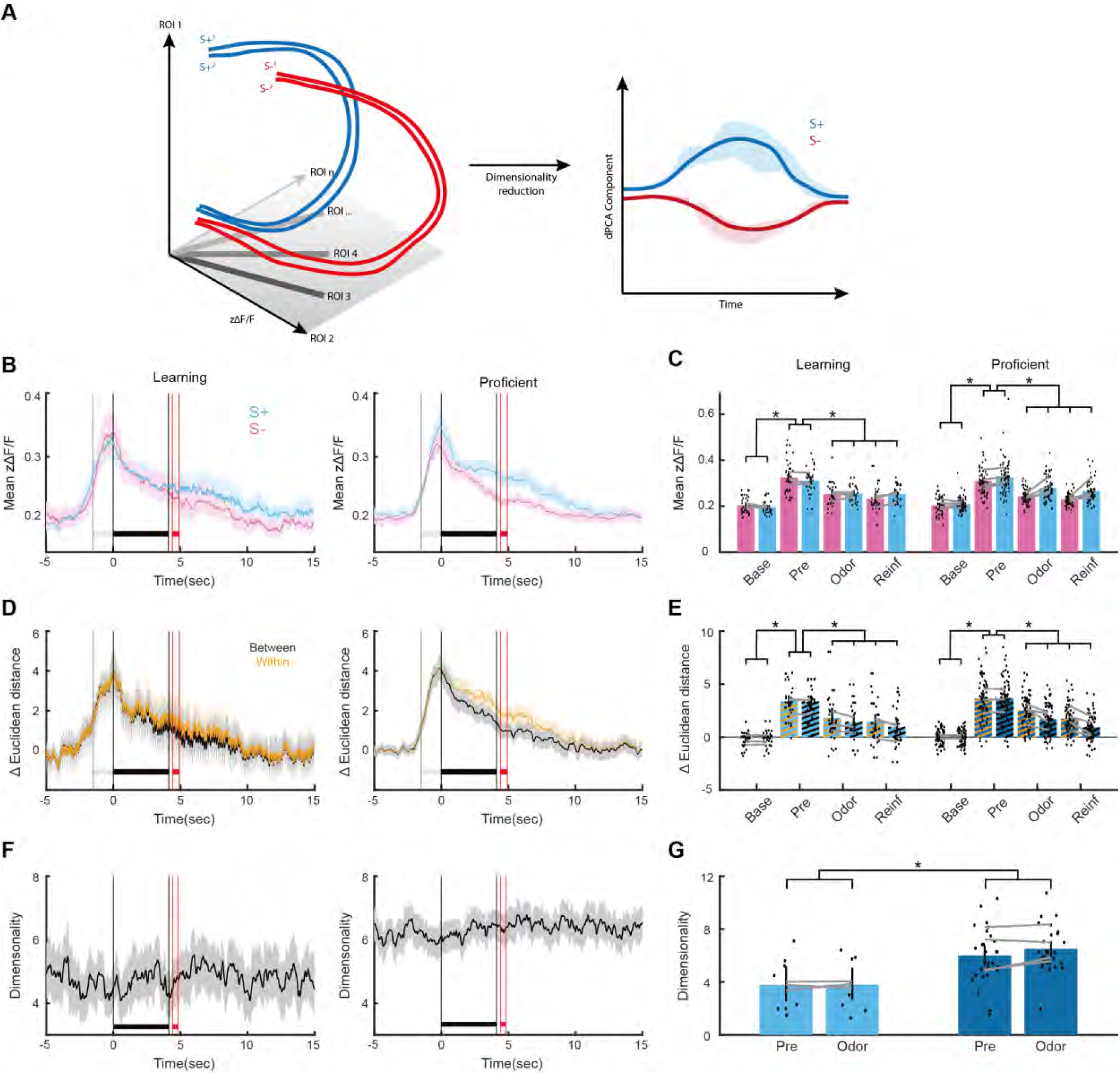
Changes in n-dimensional zΔF/F space start before the animal starts the trial and dimensionality increases with learning. (A) Diagram illustrating the trajectory for the time course of S+ and S- trials in n- dimensional zΔF/F space and how dimensionality reduction by demixed PCA (dPCA, Figure 3) reduces the dimensions to stimulus-relevant components. zΔF/F is z-scored ΔF/F (see methods). (B) Mean of per trial zΔF/F time course for S+ and S- trials for all learning and proficient sessions. Left: learning stage, right: proficient stage. (C) Bar graph showing the mean value of zΔF/F calculated in the following time windows: Base: -4.5 to -3.5 sec, Pre: -1 to 0 sec, Odor: 3.1 to 4.1 sec, Reinf: 4.5 to 5.5 sec. Dots are zΔF/F values per session and grey lines are zΔF/F values per mouse. (D) Mean per trial ΔEuclidean distance calculated in n dimensional zΔF/F ROI space. The ΔEuclidean distance was calculated: 1) At each time point for each pair of S+ vs. S- trials, and we plot the mean of all of the pair ΔEuclidean distances (we call this “Between” ΔEuclidean distance). 2) At each time point for each pair of S+ vs. S+ or S- vs. S- trials, and we plot the mean of all of the pair ΔEuclidean distances (we call this “Within” ΔEuclidean distance). (E) Bar graph showing the mean value of between and within ΔEuclidean distances calculated in the following time windows: Base: -4.5 to -3.5 sec, Pre: -1 to 0 sec, Odor: 3.1 to 4.1 sec, Reinf: 4.5 to 5.5 sec. Dots are ΔEuclidean distance values per session and grey lines are ΔEuclidean distance values per mouse. Hatch bar colors are the same as those for between and within in panel D. (F) Mean per trial dimensionality calculated in n dimensional zΔF/F ROI space. (G) Bar graph showing the mean per trial dimensionality calculated in the following time windows: Base: -4.5 to -3.5 sec, Pre: -1 to 0 sec, Odor: 3.1 to 4.1 sec, Reinf: 4.5 to 5.5 sec. Dots are dimensionality values per session and grey lines are dimensionality values per mouse. GLM statistical analysis and post hoc statistics for the significance of differences in the bar graphs are shown in Tables S2 and S3. For all figures in the manuscript: The bounded lines such as those in B, D, and F, represent the mean and 95% CI. The light grey horizontal bar is the period between the earliest trial start and odorant presentation (at 0 sec), the black horizontal bar is the time period for odorant presentation and the red horizontal bar is the period for water reward delivery.

As a first step we evaluated overall changes in mean zΔF/F averaged among all ROIs at each time step. Interestingly in this task where the mouse starts the trial by licking on the lick spout we found a substantial increase in the overall mean ΔF/F for both S+ and S-*before* the animal started the trial (pre-window compared to baseline increased by ∼50%, Figures 2B and C). There appeared to be a difference in the mean zΔF/F between S+ and S-for proficient mice (Figure 2B). However, there was no statistical difference between S+ and S-trials in the mean zΔF/F calculated in different time windows (Figure 2C). A GLM analysis yielded a statistically significant difference between the time windows (compared to baseline, p<0.05) but there was no difference between S+ and S-or between learning and proficient stages (p>0.05), 744 observations, 725 d.f., 4 mice, F-statistic=34.6, p<0.001 (Table S2).

We then assessed whether there were changes in the overall volume of n-dimensional space where each dimension is defined as the axis depicting zΔF/F for each ROI at each time point evaluated for each trial (Figure 2A). To assess changes in n-dimensional volume we calculated at each time point the Euclidean distance (Edelman, 1998) between zΔF/F for each S+ and all other S-trials (we name this distance “between S+ and S-“) as opposed to the Euclidean distance between zΔF/F for each pair of S+ trials or between zΔF/F for each pair of S-trials (“within S+ or S-“). We estimated the mean change in Euclidean distance (ΔEuclidean distance) by subtracting the mean Euclidean distance between -7 and -2 sec. The increase in mean zΔF/F shown in Figure 2B and C was mirrored by the rise in the mean ΔEuclidean distance calculated either “between S+ and S-“ or “within S+ or S-“ (Figure 2D). This increase in ΔEuclidean distance had an onset slightly before trial start indicating an expansion of n-dimensional neural space as the animal prepared to start the trial. This increase started decreasing when the odorant was presented and subsided by the end of the trial ∼15 seconds after trial start. A GLM analysis of the ΔEuclidean distance calculated for the different time windows (Figure 2E) yielded a statistically significant difference for ΔEuclidean distance calculated in the other time windows (compared to baseline, p<0.001). Still, there was no difference comparing “between S+ and S-“ vs “within S+ or S-“ or between learning and proficient stages (p>0.05), 744 observations, 725 d.f., 4 mice, GLM F-statistic=36.6, p<0.001 (Table S2).

Finally, to understand changes in overall n-dimensional space information content we estimated the dimensionality of the n-dimensional neural space (Litwin-Kumar et al., 2017). Dimensionality is a convenient measure to quantify the representation of information in a neural layer (in this case CA1) that received mixed sensory and motor inputs. Its value ranges from the number of ROIs displaying independent uncorrelated activity to one if all ROIs exhibit correlated activity. Dimensionality is closely related to the classification performance of a decoder. Figures 2F-G show that the dimensionality is constant within trials and that learning elicits increases in dimensionality. A GLM analysis of the per time window dimensionality data shown in Figure 2G yielded a statistically significant difference in dimensionality for the learning stage vs. proficient stage (p<0.05), but there was no difference comparing the pre (−1 to 0 sec) and odor (3.1 to 4.1 sec) time windows (p>0.05), 74 observations, 67 d.f., 4 mice, GLM F-statistic=11.5, p<0.001 (Table S3). These results indicate that the dimensionality of the initially unconstrained n-dimensional neural space increases as the mouse’s dCA1 network changes with learning.

### Demixed Principal Component Analysis Reveals Increased Differential Response to Stimuli upon Learning

Mean zΔF/F did not differ between S+ and S-trials, and did not change with learning (Figures 2B-C). This is not surprising given the heterogeneity of CA1 neural activity (e.g. Figure 1D). To assess changes in neural activity with learning and differences in activity between rewarded (S+) and unrewarded (S-) stimulus trials we assessed changes in neural activity after dimensionality reduction. We performed a demixed principal component analysis (dPCA) (Kobak et al., 2016) of the zΔF/F population data per mouse to evaluate changes in principal components quantifying neural activity in the axis maximizing stimulus variance. An example of dPCA for a proficient mouse is shown in Figure S5. dPCA estimates stimulus components (Figure S5E) and stimulus-independent components (Figure S5F). Figure S5A and B show the explained variance of the first 10 components. We used the sum of stimulus components to estimate the divergence of S+ vs. S-trial component and their changes with learning. Figure S5Di shows that the sum of stimulus components diverged shortly after trial start. As expected, the sum of independent components did not differ between S+ and S-trials (Figure S5Dii).

Figure 3 shows the results of dPCA analysis for all mice. The sum of the stimulus principal components diverged between S+ and S-trials shortly after trial start and the magnitude of the divergence increased as the animal learned (Figure 3A and B). The sum of stimulus-independent components did not differ between stimuli and did not display changes as a function of time (Figure 3C and D). A GLM analysis yielded for the sum of the stimulus principal components a statistically significant difference for the interaction between the learning vs. proficient stages and the odor time window (or the reinforcement time window) vs. the baseline time window and the interaction between S+ and S-, the odor time window (or reinforcement window) vs. the baseline time window and the learning vs. proficient stages (p<0.01), 48 observations, 32 d.f., 4 mice, GLM F-statistic=10.6, p<0.001 (Table S4). The GLM analysis did not yield statistically significant changes between S+ and S-or for the odor time window for stimulus-independent components (Table S4).

**Figure 3.**
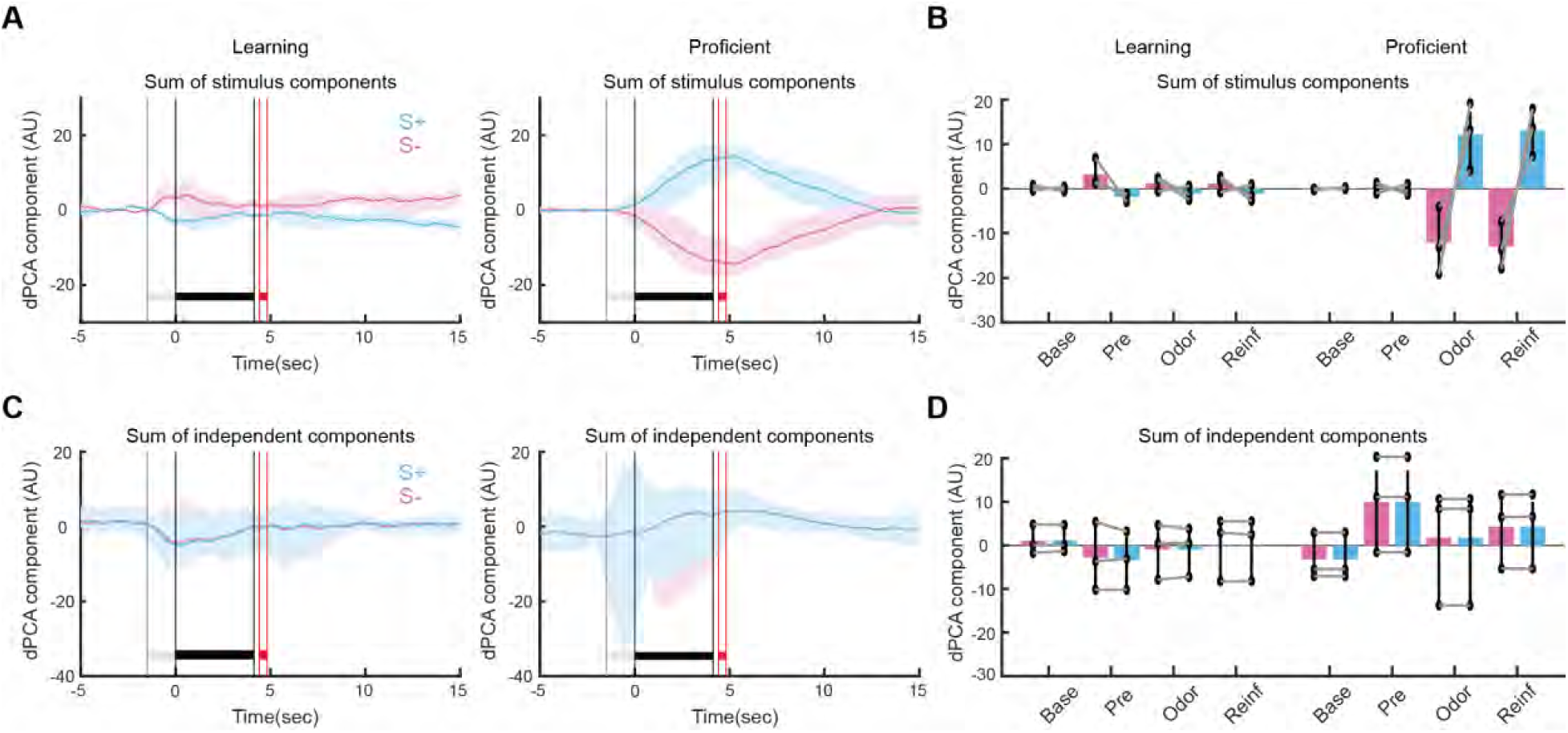
Demixed PCA reveals a substantial increase elicited by learning in the divergence between S+ and S-stimulus components. (A) Time courses for the sum of demixed PCA (dPCA) stimulus components for the learning (i, 20 sessions, 4 mice) and proficient (ii, 55 sessions, 4 mice) stages. (B) Bar graph showing the mean value of the sum of stimulus dPCA components calculated in the following time windows: Base: -4.5 to -3.5 sec, Pre: -1 to 0 sec, Odor: 3.1 to 4.1 sec, Reinf: 4.5 to 5.5 sec. (C) Time courses for the sum of independent dPCA components for the learning (i, 20 sessions, 4 mice) and proficient (ii, 55 sessions, 4 mice) stages. (D) Bar graph showing the mean value of the sum of independent dPCA components calculated in the same time windows as in (B). dPCA was performed following the procedure in (Kobak *et al*., 2016). An example for all components and dPCA explained variance for one mouse is shown in Figure S5. GLM statistical analysis and post hoc statistics for the significance of differences for the bar graphs are shown in Table S4.

### Stimulus-divergent zΔF/F Responses are Heterogeneous, Divergence Increases with Learning and the Onset of Divergence Takes Place at Discrete Times

The results of dPCA shown in Figure 3 indicate a substantial increase in the divergence between S+ and S-as the animal learns. We asked whether this divergence of the responses is heterogeneous when assessed on a per ROI basis. We classified zΔF/F responses in S+ or S-trials within a session as divergent when the p-value for a GLM of the difference between S+ and S- in the time span from -1 to 5.5 sec was smaller than the p-value for significance corrected for multiple comparisons for all sessions using a false discovery rate (Curran-Everett, 2000). Figure 4A shows the time courses for zΔF/F for S+ and S-trials for all divergent ROIs for the proficient stage and Figure 4C shows the cross correlogram of the zΔF/F time courses. We sorted the time courses into three clusters using a hierarchical binary cluster tree. The average zΔF/F time courses for the three clusters are shown in Figure 4E. Cluster 1 displayed a bimodal time course where zΔF/F for S+ increased above S-shortly after the trial started, and this reversed after the reward when S-increased above S+. Cluster 2 displayed an increase in zΔF/F for S+ and a decrease in zΔF/F for S-shortly after the trial started. Cluster 3, that was the cluster with the largest number of ROIs displayed a robust increase in zΔF/F for S- and a smaller increase for S+. Therefore, time courses for zΔF/F for divergent ROIs for the proficient stage display substantial heterogeneity.

**Figure 4.**
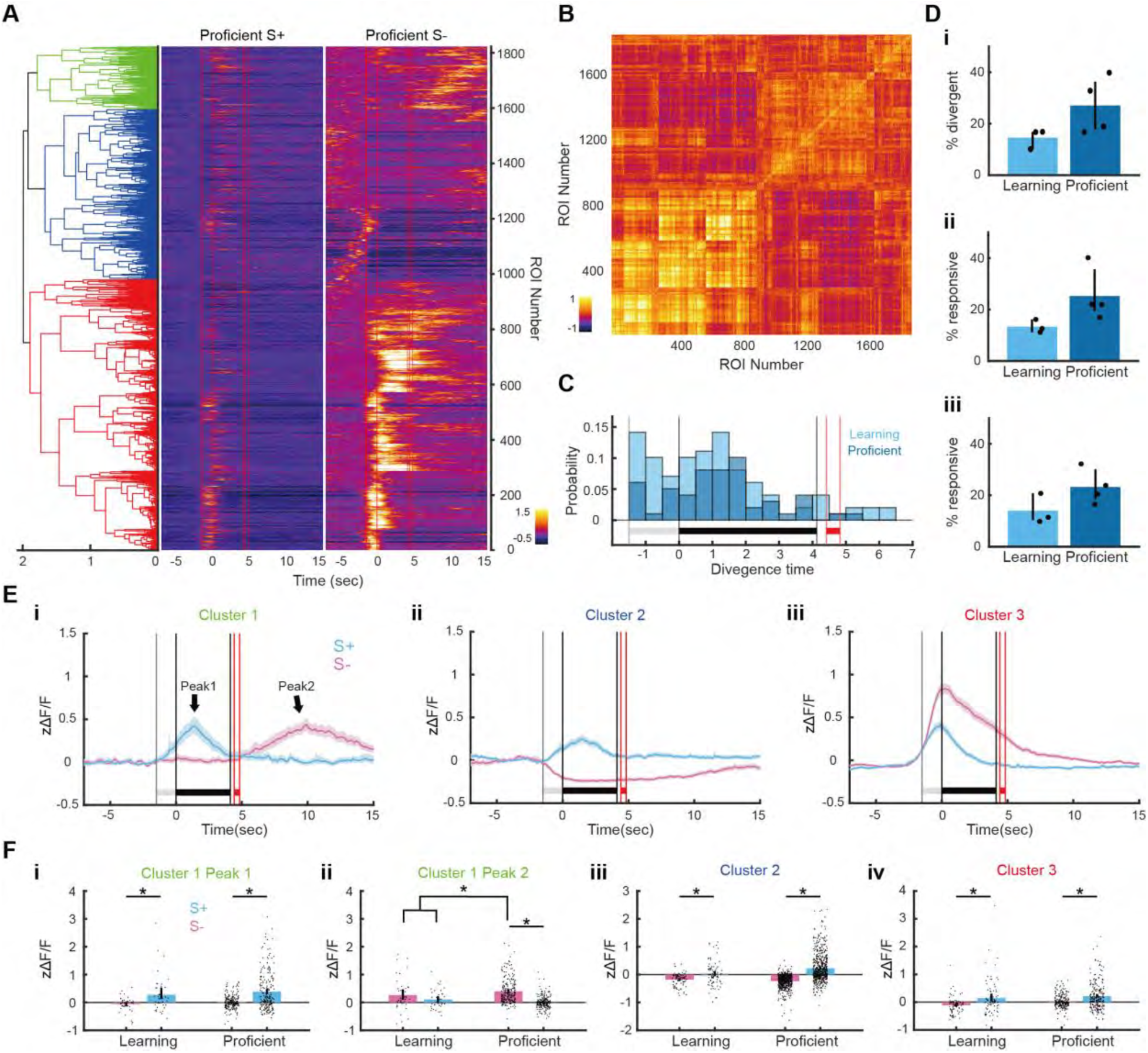
The magnitude of divergence between S+ and S-zΔF/F responses increases with learning and the onset of divergence takes place at discrete times that differ between the learning and proficient stages. (A) zΔF/F time courses for single ROIs that were divergent between S+ and S-trials in the proficient sessions (66 sessions, 4 mice). Time courses were sorted by estimating an agglomerative hierarchical cluster tree shown on the left that was calculated using the cross-correlation coefficients between all divergent zΔF/F time courses shown in B. The red vertical lines show (in order): trial start, odorant on, odorant off, reinforcement on and reinforcement off. (B) Cross-correlation coefficients computed between all per trial zΔF/F time courses shown in A. The coefficients were sorted by the agglomerative hierarchical cluster tree shown in A. (C) Histograms for divergence times for all the ROIs for the proficient stage shown in A (dark blue) and for all divergent ROIs for learning stage sessions (light blue). Divergence time histograms are significantly different between learning and proficient (ranksum p<0.001, n=99 divergence time bins). (D) i. Percent divergent ROIs per mouse. ii and iii. Percent responsive ROIs per mouse for S+ and S-trials. The percent divergent (i) and percent responsive (ii and ii) differences between learning and proficient stages were not statistically significant (two tailed t test p>0.05, n=3 for learning and 4 for proficient, 5 d.f.). (E) Mean zΔF/F time courses for the three clusters in the hierarchical tree shown in A. (F) Peak zΔF/F calculated per session for each cluster. Note that cluster 1 has two peaks (see arrows in Ei). Table S5 shows the GLM statistical analysis for Figure 4F.

Analysis of zΔF/F divergent ROIs for the learning stage also displayed a similar heterogeneity in time courses (Figure S6B). The percent of divergent ROIs tended to be smaller for the learning stage, but was not significantly different from the proficient stage (Figure 4D, t-test, p>0.05, 3 mice for learning, 4 mice for proficient). However, when the per cluster zΔF/F was tested with GLM analysis for statistically significant differences using learning stage and S+ vs. S- as independent variables (Figure 4F) there were statistically significant differences for S+ vs. S- for peak 1 for cluster 1 and for learning vs. proficient and for the interaction of learning vs. proficient and S+ vs. S- for peak 2 for cluster 1. Moreover, there are statistically significant differences for S+ vs. S- and for the interaction of learning stage and S+ vs. S- for cluster 2 and for learning vs. proficient and for learning vs. proficient and S+ vs. S- for cluster 3 (p<0.05, number of observations, d.f. and F statistics are in Table S5). These differences indicate that there are changes in zΔF/F time course elicited by learning.

Finally, responses of a subset of CA1 pyramidal cells that have been named odor-specific time cells take place at discrete time points in the delay period in a delayed non-match to sample task where the animal receives a reward when they respond in the trial where the second odor differs from the first odor (Taxidis *et al*., 2020)(also see (MacDonald *et al*., 2013). Interestingly, perusal of the pseudocolor time courses for zΔF/F in Figure 4A appeared to show differences in the time onset of the divergence.

We assessed the divergence time for each divergent ROI. Figure S6A shows examples of the time course for zΔF/F for a subset of ROIs. The divergence takes place at discrete time points after the start of the trial, ranging from the onset of divergence close to the start of the trial (Figure S6A ii) to the onset close to the time when the reward was delivered (Figure S6A vi). Figure 4C shows histograms of the divergence times for the learning and proficient stages. The divergence times spanned the time period from trial onset to times after the reward was delivered. The divergence times for the learning stage occurred earlier than those for the proficient stage (ranksum p<0.001, 99 divergence time bins in the histograms).

### Switching the Rewarded Stimulus to Unrewarded Elicits Changes in zΔF/F Stimulus Responses Consistent with Response to the Valence of the Stimulus

In order to determine whether stimulus-divergent zΔF/F responses are divergent responses to the stimulus identity (odorant identity or odorant valve click) vs. stimulus valence (are the stimuli rewarded?) we switched the rewarded S+ stimulus (1% heptanal, HEP) with the unrewarded S-stimulus (mineral oil, MO) after the animal became proficient. We call this a switch from a forward go-no go task when HEP was S+ to a reversed go-no go task where MO became S+. As shown in Figures S2A-D, percent correct behavior decreased below 50% after stimulus reversal. Subsequently, it recovered, eventually reaching >80% indicating that the animal learned the new reversed valence of the stimulus. Figure 5A shows the zΔF/F time course for ROIs that responded with divergence to the stimuli when the animal was proficient in the forward task. As performed for zΔF/F time courses in Figure 4 we performed a cross-correlation analysis for this forward task (shown in Figure S7C) and we sorted the time courses into three clusters using a hierarchical binary cluster tree. The average zΔF/F time courses for cluster 2 that responded with a large increase to S- is shown in Figure 5C and the average time courses for the other clusters are shown in Figure S7A and B. As in Figure 4 the responses to S- (MO) tended to be larger than responses to S+ (HEP) in the most abundant cluster of responses (cluster 3) in this forward task (Figure 5A).

**Figure 5.**
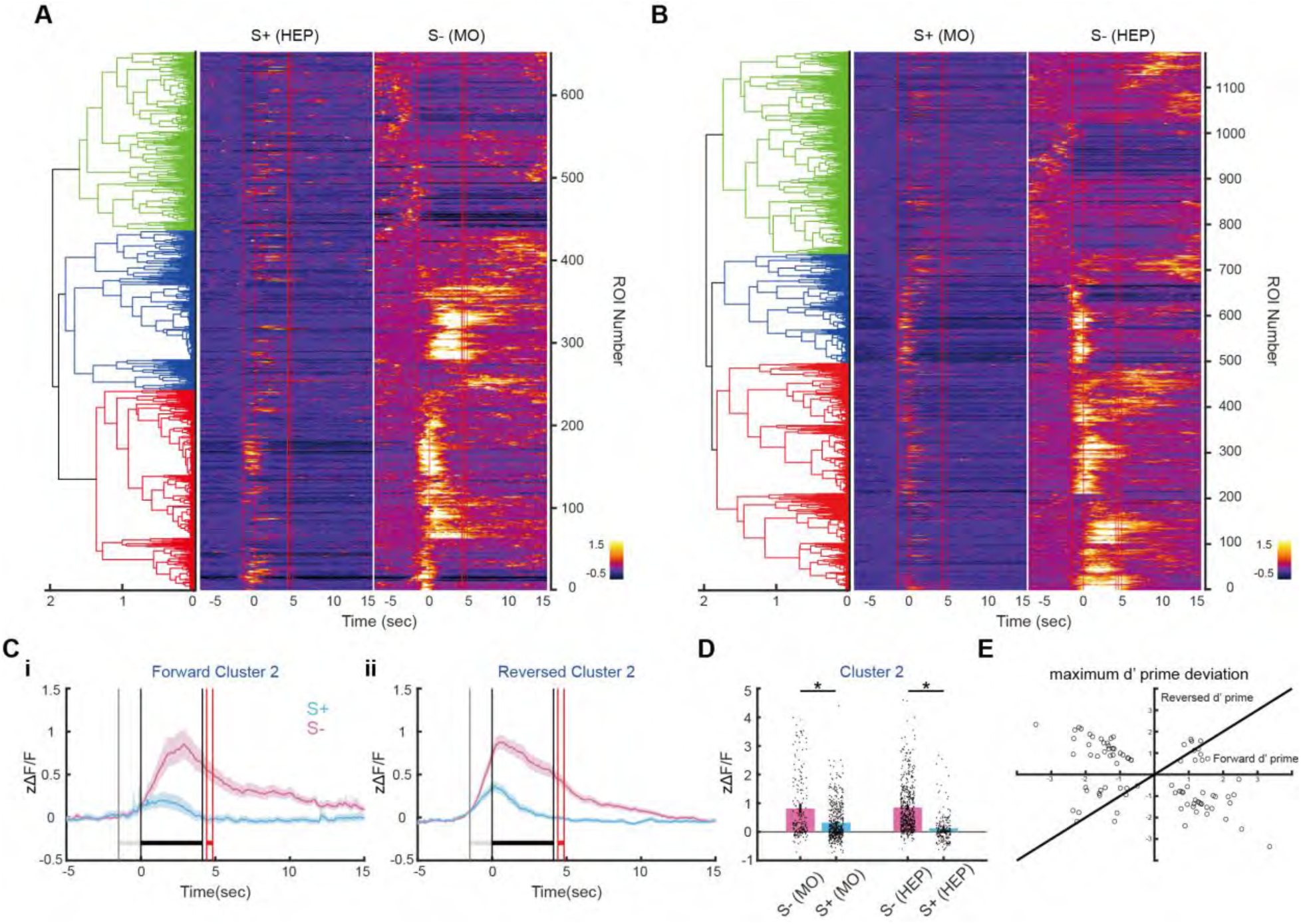
Reversal of odorant valence reveals that a substantial number of dCA1 cells respond to stimulus valence. (A and B) zΔF/F time courses for single ROIs that were divergent between S+ and S-trials for proficient mice. (A) shows zΔF/F time courses for the forward proficient sessions where the rewarded odorant (S+) was HEP and the unrewarded odorant (S-) was MO (25 sessions, 4 mice) and (B) shows time courses for the reversed proficient sessions where the rewarded odorant (S+) was MO and the unrewarded odorant (S-) was HEP (41 sessions, 4 mice). For both A and B time courses were sorted by estimating an agglomerative hierarchical cluster tree shown on the left that was calculated using the cross-correlation coefficients between all divergent zΔF/F time courses shown in Figure S7 C. The red vertical lines in A and B denote (in order): trial start, odorant on, odorant off, reinforcement on and reinforcement off. (C) Mean zΔF/F time courses for cluster 2 (blue cluster in A and B). Mean zΔF/F time courses for clusters 1 and 3 are shown in Figure S7. (D) Peak zΔF/F calculated per session for cluster 2. Table S6 shows the GLM statistical analysis for these data. (E) Relationship for recordings in one mouse between forward and reversed peak d’ for HEP vs. MO for all forward stimulus divergent zΔF/F time courses for a set of forward/reversed sessions where ROIs were matched from session to session. The per ROI zΔF/F and d’ time courses for these forward and reversed sessions are shown in Figure S8. The line shown is d’ reversed = d’ forward, which would be followed if the ROIs represent stimulus identity.

Interestingly, Figures 5A-D show that when we reversed the stimuli the majority of the divergent zΔF/F time courses still responded with larger increases for the S-stimulus (cluster 2, now switched to HEP, also see mean zΔF/F time courses for each cluster in Figures 5D and S7A,B). Therefore in both the forward and reversed tasks the unrewarded odorant stimuli elicited a larger zΔF/F response in a large portion of cluster 2 ROIs when the animal was proficient. We quantified this shift in zΔF/F time courses with reversal by determining the peak value for stimulus-induced changes in zΔF/F. Figure 5D shows a bar graph of the peak changes in zΔF/F for forward and reversed runs for cluster 2 and Figure S7Aiii and Biii show the results for clusters 1 and 3. When the zΔF/F peak values were tested with GLM analysis for statistically significant differences using odorant and forward vs. reversed as independent variables there were statistically significant differences for both odorant and forward vs. reversed (p<0.001, number of observations, d.f. and F statistics are in Table S6). These data suggest that the most of the zΔF/F stimulus responses are responses to the odorant value (is the odorant rewarded?) instead of odorant identity.

For the mouse whose percent correct behavior is shown in Figure S2D we performed the last forward session and the first proficient reversed session on the same day allowing us to perform an analysis where the ROIs in the reversed session were matched to the ROIs determined in the forward session. Figure S8A shows divergent zΔF/F responses for matched ROIs in forward and reversed reward proficient sessions and Figure S8C shows the d’ time course calculated for HEP vs. MO for these divergent zΔF/F responses. d’ was calculated per time point for zΔF/F time courses for HEP vs.

MO. As defined (see methods), a change in d’ polarity indicates that the response is a response to the valence. Figure 5E plots the peak value of d’ in the odorant period for the forward and reversed runs for all the matched ROI divergent d’ responses shown in Figure S8C. Most of the d’ values reverse polarity with the forward to reversed switch indicating that these ROIs represent the odorant value. However, a smaller number kept the polarity of the response as the odorant reward was reversed, indicating that a smaller number of ROIs does respond to the stimulus identity. In conclusion, for proficient sessions (>80% correct behavior), most of the zΔF/F divergent ROIs represent stimulus valence, and fewer ROIs respond to the odorant identity.

### The Accuracy for Decoding the Stimulus Increases with Learning and is Dependent on the Fit Window

We asked whether the information embedded within zΔF/F activity of all ROIs can be used to decode the stimulus (S+ vs. S-). Figure 6 shows the time courses for the accuracy of GLM decoding of stimulus identity (other decoding algorithms yielded similar results; see Figure S9). GLM was trained with different time windows using zΔF/F activity of all ROIs in all S+ and S-trials within each session to evaluate the dependence of decoding accuracy per trial on the information embedded within particular time windows. As a control time courses for accuracy are shown where the trial identity (S+ or S-) was shuffled (red time courses in i and ii panels and light grey bars in iii panels in Figure 6). When we trained the GLM using a broad fit time window spanning the odorant period and the beginning of the reward period (0.5 to 5.5 sec) decoding accuracy started increasing above 0.5 slightly after trial start (∼-1 sec) and reached ∼0.8 through a window spanning the later 3 sec of the odorant window for the proficient animal (Figure 6Aii). In contrast, the decoding accuracy for the learning stage only reached ∼0.65 (Figure 6Ai). The bar graph in Figure 6Aiii shows the data for pre (−1 to 0), odor (3.1 to 4.1) and reinforcement (4.5 to 5.5) time windows. A GLM analysis yielded a statistically significant difference for odor and reinforcement time windows vs. pre-window and for the interaction between these time window comparisons and proficient vs. learning stages (p<0.05-0.001), 279 observations, 270 d.f., 4 mice, GLM F-statistic=37.7, p<0.001 (Table S7).

**Figure 6.**
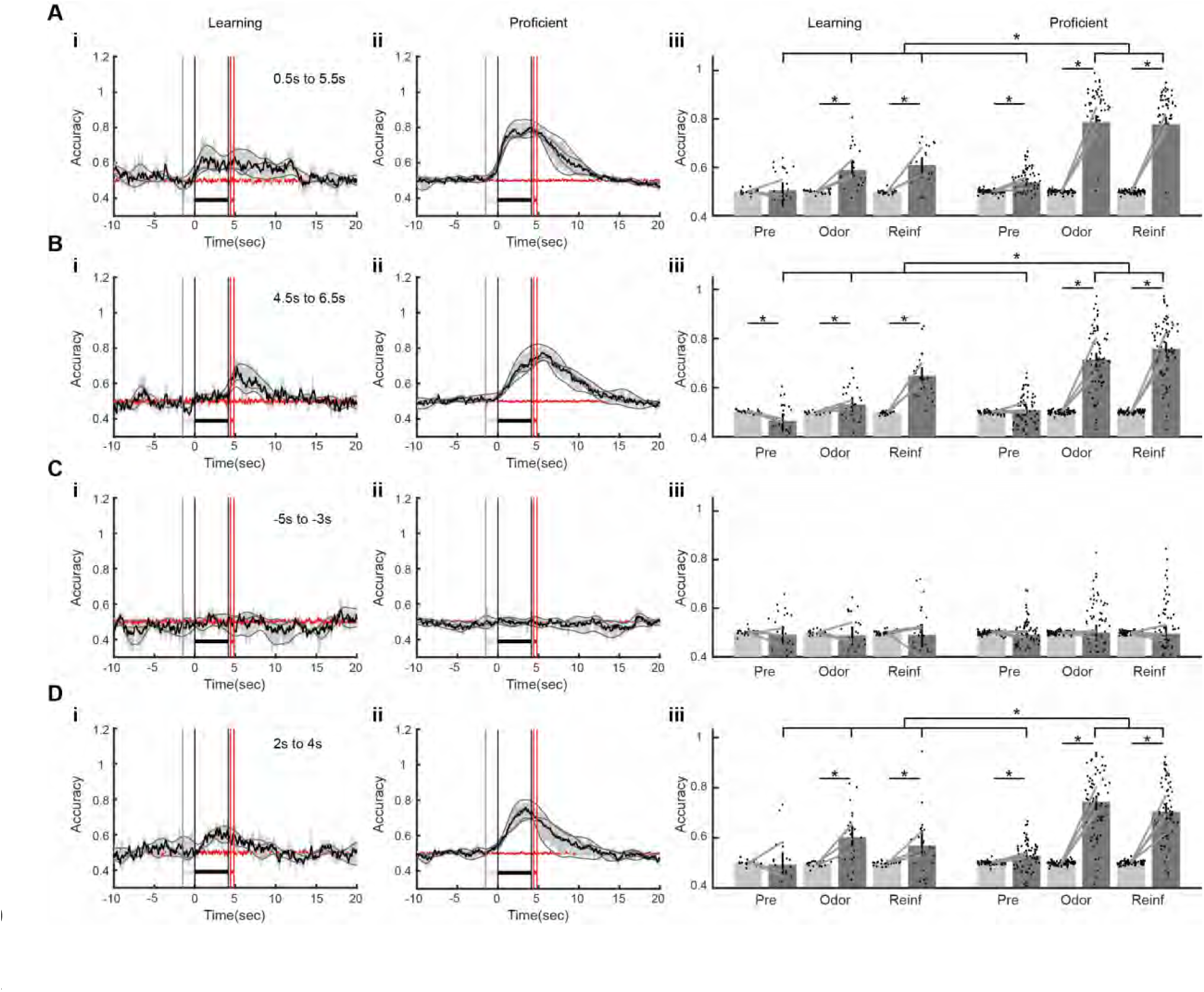
Learning elicits an increase in stimulus decoding accuracy. (A to D). Each panel shows the accuracy of GLM decoding of the stimulus (S+ vs. S-) from per trial zΔF/F responses for all trials and all ROIs per session for all learning stage sessions (i) and proficient sessions (ii) (4 mice, 18 learning sessions, 66 proficient sessions). The bounded black line represents the mean accuracy bounded by the 95% CI. The grey lines are per mouse accuracy. The red line is the stimulus decoding accuracy calculated after shuffling the stimulus labels (S+ vs S-). A to D differ by the time period used to train the GLM decoder. The A training period (0.5 to 5.5 sec) covers odorant presentation and reward delivery, the B training period included reward delivery (4.5 to 6.5 sec), the training period for C takes place before trial onset (−5 to -3 sec) and the training period for D was the latter half of the odorant period (2-4 sec). The bar graphs in iii show the mean accuracy for different accuracy periods (Pre -1 to 0, Odor 3.1 to 4.1 and Reinf 4.5 to 5.5). Light gray bars are the shuffled stimulus accuracies. Points are per session accuracies and bars are 95% CIs. *p<0.05 for a pFDR-corrected t-test or ranksum tests, GLM statistics are in Table S7.

We then asked how the decoding accuracy time course varies depending on the fit time window. As expected when the fit window spanned a period before stimulus presentation (−5 to -3 sec) accuracy did not increase above shuffled (Figure 6C, the p-value for GLM was >0.5 for all comparisons, see Table S7). In contrast, when the window spanned the reinforcement period (4.5 to 6.5 sec) decoding accuracy increased above 0.5 for the learning stage after the mouse was given the reinforcement (Figure 6Bi) and the increase in decoding accuracy above 0.5 shifted to the time when the odorant was presented when the mouse became proficient (Figure 6Bii). GLM analysis of the bar graph in Figure 6Biii yielded a statistically significant difference for odor and reinforcement time windows vs. pre-window and for the interaction between odor window vs. pre-window and proficient vs. learning stages (p<0.05-0.001), but was not significant for the interaction between reinforcement window vs. pre window and proficient vs. learning stages (p>0.05), 273 observations, 264 d.f., 4 mice, GLM F-statistic=45, p<0.001 (Table S7). Finally, the increase in accuracy for the proficient mice did not reach a plateau during the odorant period when the fit window was a narrow window at the end of the odorant period (2 to 4 sec) (Figure 6D, see GLM results in Table S7). The results indicate that zΔF/F activity of all ROIs contains information sufficient to decode the stimuli. As expected this information differs when different time windows are used to fit the GLM decoding algorithm.

### Between Trials Stimulus Decoding is Biased to Predict the Unrewarded Stimulus and there are Sudden Brief Replays of Rewarded Stimulus Prediction

A hippocampal offline replay of neural patterns plays a role in spatial learning (Michon et al., 2019; Singer et al., 2013). We explored whether a replay of neural activity for prediction of the stimulus happened between trials for proficient mice engaged in the go-no go task. Figure 7A shows a time course for predicting the stimuli by GLM decoding calculated during an imaging session. As in Figure 6A, GLM was fit to neural activity for all ROIs during the 0.5 to 5.5 sec spanning most of the odorant window. A prediction value of 1 corresponds to S+ prediction and a value of 0 represents S-. The blue shade shows the 5^th^ to 95^th^ percentile band for prediction calculated after shuffling the stimulus labels. As expected, prediction increases to 1 for some time longer than the odorant window during S+ trials while it increases briefly at the start of S-trials but returns to zero before the end of the odorant period. Figure 7B shows the mean prediction time course for all trials in proficient sessions for the 4 mice (a total of 1848 trials). For this within-trial decoding the prediction starts increasing before trial start for both the S+ and S-trials and it diverges between S+ and S- at the start of the trial. The bar graph in Figure 7C shows the S+ and S-prediction for a baseline period (−2.5 to -1.5) and an odor period (2 to 4.1 sec). A GLM analysis yields significant differences for baseline vs. odor time windows and for the interaction between S+ vs. S- and time windows, p<0.001, 4 mice, 220 observations, 213 d.f., F-statistic 194, p <0.001 (Table S8).

**Figure 7.**
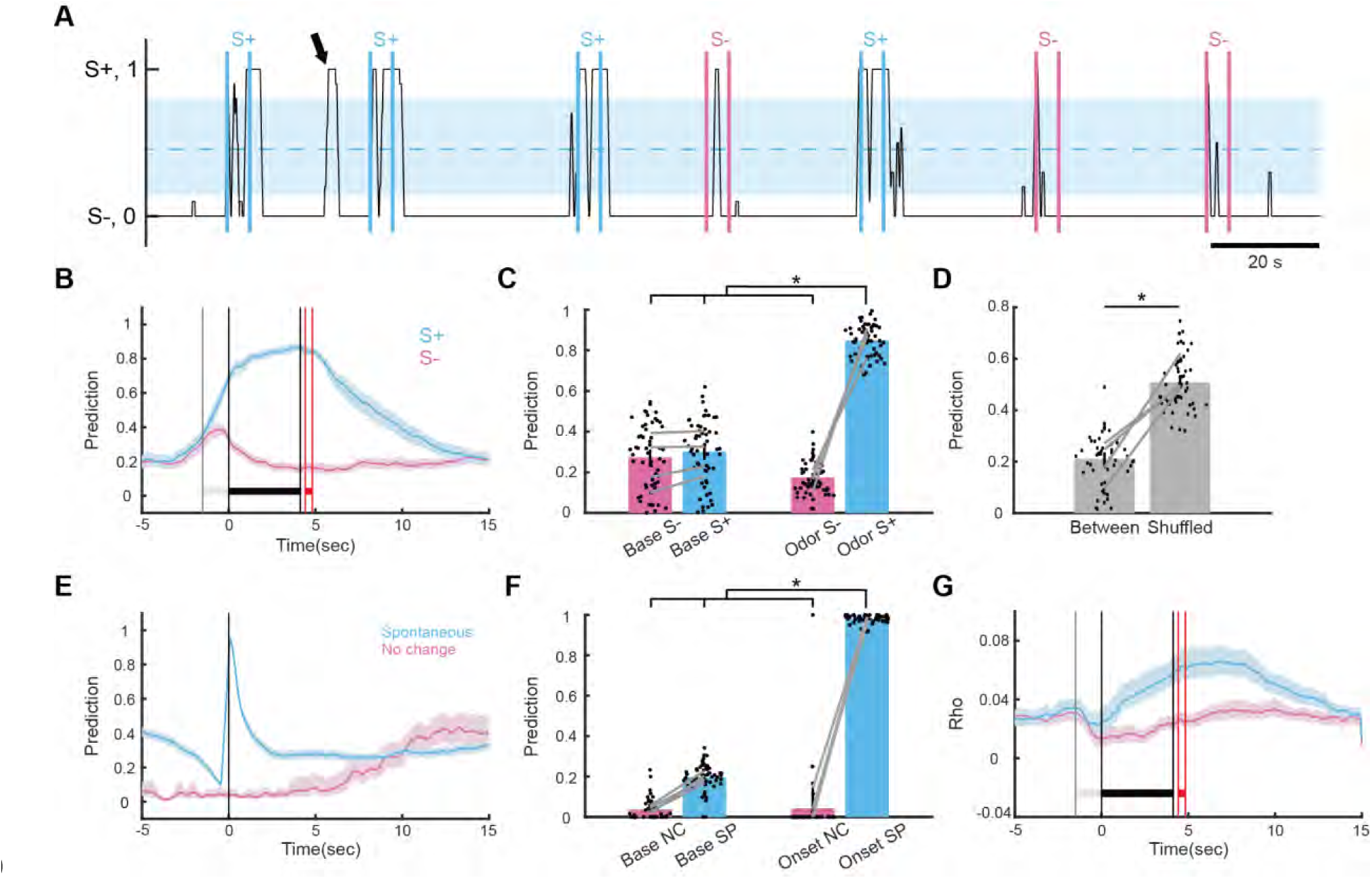
Replay of prediction of S+ for stimulus decoding between trials. (A) Example of the time course for stimulus decoding prediction during a session when the animal was proficient in the go-no go task. Prediction ranges from 0 (S-) to 1 (S+). Cyan vertical bars are the odor on and odor off times for the S+ trials and magenta vertical bars are the odor on and odor off times for the S-trials. The shaded blue area is the 5 to 95 percentile decoding prediction calculated for decoding with shuffled trial labels. The arrow points to a sudden shift in prediction above the shuffled 95 percentile that took place between trials (we call this a prediction replay). (B) Mean label prediction time course within trials calculated for S+ and S-trials for all sessions when the mice were proficient (66 sessions, 4 mice). (C) Bar graph quantifying mean label prediction for S+ and S-trials in two time windows: Base (−2.5 to -1.5 sec) and odor (2 to 4.1 sec). Dots are per session mean label prediction and grey lines are per mouse mean label prediction. (D) Bar graph quantifying the mean label prediction between trials compared to the mean label prediction between trials for shuffled label decoding. (E) Mean label prediction time course for between-trial prediction replays (“replay”) and for time courses in between trials periods where we did not find spontaneous shifts to S+ (“no change”) (66 sessions, 4 mice). (F) Bar graph quantifying mean label prediction for between-trial spontaneous prediction shifts and for no change between-trial prediction time courses calculated in two time windows: Base (−2.5 to -1.5 sec) and odor (2 to 4.1 sec). Dots are per session mean label prediction and grey lines are per mouse mean label prediction. (G) Correlation between label prediction calculated at each time point in the prediction time course for all S+ or S-trials within a session and the label prediction value found at the point of spontaneous shift for spontaneous shifts in prediction found in between trials in the same session (66 sessions, 4 mice). *p<0.05 for a pFDR-corrected t-test or ranksum tests, GLM statistics are in Table S8.

Intriguingly, the average prediction is lower than shuffled in between trials (Figure 7A and 7D) indicating that the default decoding is S- for most of the time between trials (GLM p-value for between vs. shuffled <0.001, 110 observations, 105 d.f., F-statistic 70.7, p<0.001, Table S8). However, there are precise brief shifts of prediction to 1 (predicting S+, e.g. arrow in Figure 7A). We identified these spontaneous shifts of prediction from 0 to 1 between trials by finding increases beyond the 95 percentile of the shuffled prediction values (blue shade in Figure 7A). We found a total number of 1635 spontaneous shifts to S+ prediction. The duration of these spontaneous shifts in prediction varied between a fraction of a second to two seconds (example in Figure 7A). The blue bounded line in Figure 7E shows the mean time course for between trial spontaneous shifts compared to an adjacent time period with no spontaneous changes in prediction (“no change”, magenta). The bar graph in Figure 7F shows a significant difference between the prediction before the onset of the spontaneous increase in prediction and the peak of prediction for the spontaneous shift. A GLM analysis yields significant differences for spontaneous vs. no change and for the interaction between spontaneous vs. no change and baseline vs. peak time windows, p<0.001, 4 mice, 192 observations, 185 d.f., F-statistic 703, p <0.001 (Table S8).

Finally, we calculated the correlation between zΔF/F time courses for spontaneous shift and zΔF/F time courses for either S+ or S-trials for all ROIs within each session to determine whether the neural activity underlying the spontaneous prediction shifts from S- to S+ are more correlated with S+ vs S-. Figure 7G shows the time courses for the average correlations. The correlation of spontaneous zΔF/F with S+ diverges from the correlation with S-shortly after the start of the trial and remains high for a few seconds after the reward. A GLM analysis yields significant differences in average correlations for comparison of the time windows (baseline vs. odor) and for the interaction between S+ vs. S- and time window, p<0.001, 4 mice, 216 observations, 209 d.f., F-statistic 15.7, p-value <0.001 (Table S8). The sudden shift in prediction from S- to S+ between trials and the fact that zΔF/F activity during these spontaneous shifts correlates with zΔF/F neural activity for S+ suggest that these events are replay of neural activity for the rewarded stimulus.

### Decoding of Decision Making in the Go-No Go Task from CA1 Neural Activity is Time-tiled

Divergence of zΔF/F responses between S+ and S- stimuli occurs at different times after the trial start (Figure 4C) suggesting that the onset of the increase in accuracy for stimulus decoding above the accuracy for the shuffled control, the time for decision-making, is time-tiled. In order to assess time tiling of the time for decision making (is the stimulus S+ or S-), we performed an analysis of the time course of stimulus decoding from zΔF/F responses of *subsets of ROIs* per session for mice proficient in the go-no go task. The number of ROIs used to calculate GLM decoding per session was varied from 1, 2, 5 to 15 ROIs per session and the results were compared to decoding with all ROIs in each session. We first performed a visual survey of the within-trial dynamics of stimulus decoding accuracy when the decoding was performed with zΔF/F responses of single ROIs (Figure S10). We found that decoding accuracy for different ROIs started increasing at different times after the trial started and reached accuracy values above 0.65. Three examples of decoding accuracy time courses and their corresponding zΔF/F time courses for S+ and S- trials are shown in Figure S10 A-C. In Figure S10 A, accuracy increases to ∼0.6 at the start of the trial (immediately following valve click and before odorant addition) and increases to a higher value (∼0.8) after odorant presentation. In Figures S10 B and C, there is no increase in accuracy at the start of the trial and accuracy increases either shortly after odorant addition (Figure S10 B) or ∼3 sec after odorant addition (Figure S10 C). A histogram of average accuracy values computed during the odor window (3.1 to 4.1 sec) showed a clear difference between S+ and S- trial zΔF/F for these three examples (Figures S10 A-Ciii). In addition, we found a subset of ROIs whose decoding yielded decreases in accuracy below 0.5. An example is shown in Figure S10 D, where the accuracy decreased to ∼0.1. These decreases in accuracy took place for ROIs with largely overlapping S+ and S- odorant period zΔF/F values except for one or two trials where the zΔF/F deviated from the other trials as evidenced in the histogram in Figure S10 Diii.

We then performed stimulus decoding analysis with subsets of ROIs for the entire dataset for mice proficient in the go-no go task (four mice, 66 sessions). Figure 8A shows histograms of decoding accuracy calculated in the odor window (3.1 to 4.1 sec) for stimulus decoding performed with zΔF/F from subsets of ROIs. As the number of ROIs per decoding run decreases, stimulus decoding accuracy declines (see Figure 8Av with 8Ai). In addition, accuracies calculated in the pre window (−1 sec to 0 sec) are smaller than accuracies calculated in the odor window (compare Figures 8A and 8B), and the pre accuracies also appear to decrease as the number of ROIs per decoding run is decreased (compare Figure 8Bv with 8Bi). A GLM analysis yields significant differences in decoding accuracy for comparisons between the number of ROIs and the time windows, p<0.001, 4 mice, 27344 observations, 27344 d.f., F-statistic 798, p-value <0.001 (Table S9).

**Figure 8.**
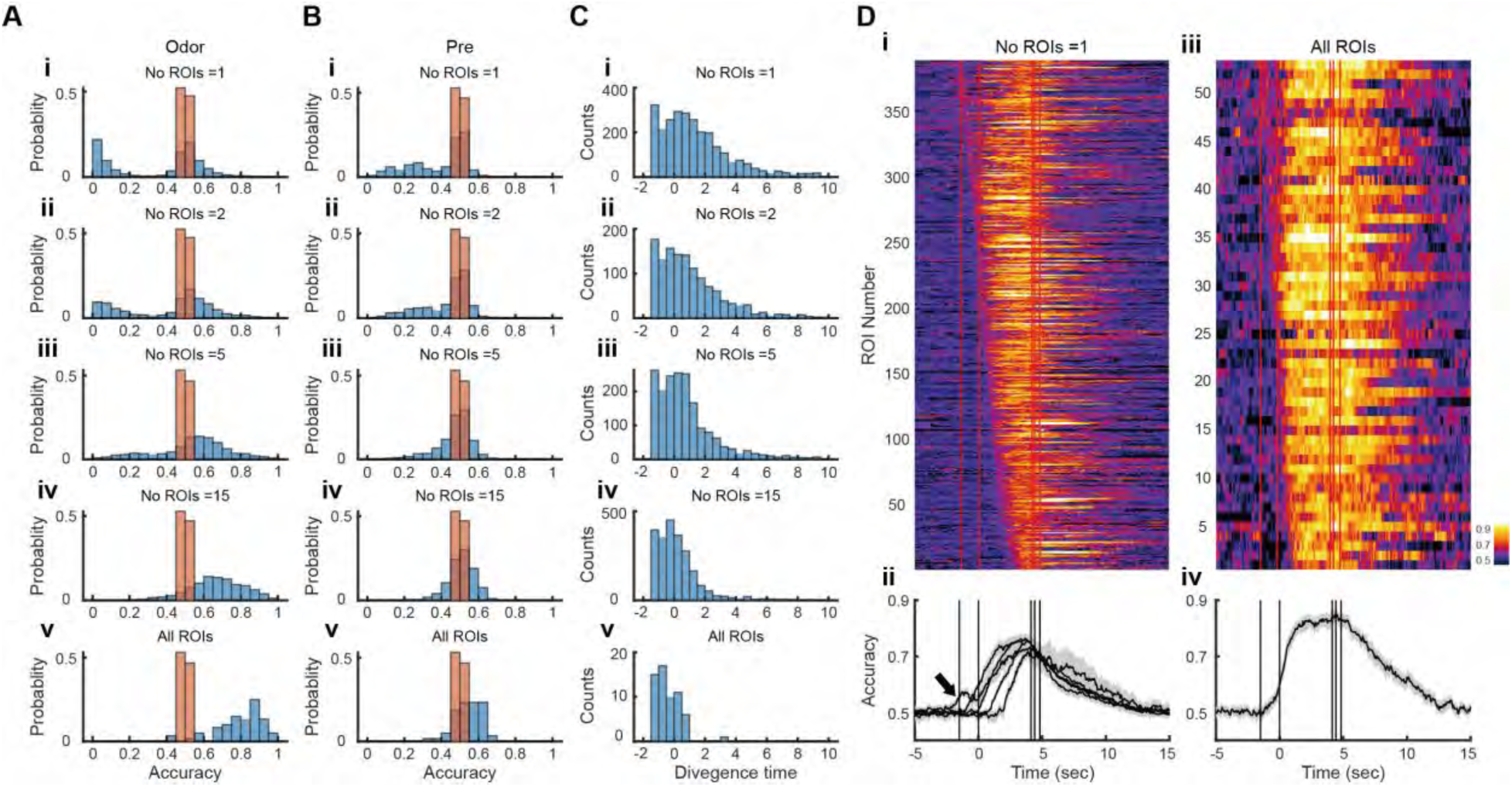
Stimulus decoding accuracy calculated with different subsets of ROIs per session reveals time tiling of the increase in stimulus decoding accuracy. (A) Histogram for stimulus decoding accuracy calculated in the odor period (3.1 to 4.1 sec) for all proficient sessions (66 sessions) for 4 mice. i to v show the histogram for an increasing number of ROIs per decoding session. Blue is stimulus decoding accuracy, brown is stimulus decoding accuracy calculated after shuffling the labels. Histograms were normalized by dividing by the total number of counts. (B) Histograms calculated in the Pre period (−1 to 0 sec). (C) Histograms for the onset of the increase in decoding accuracy for decoding runs that achieved at least 0.65 accuracy after trial start. (D) (i). Time courses for decoding accuracies calculated for a single ROI per session that reach at least 0.65 after trial start. (ii) Mean accuracy time courses calculated for one ROI accuracy time courses shown in Di with accuracy increase onsets in the following time windows: -1.5 to -1, -1 to 0, 0 to 1, 1 to 2, 2 to 3, >3 sec. (iii) Time courses for decoding accuracies calculated all ROIs per session. (iv) Mean accuracy time course for the time courses shown in Diii. GLM analysis indicates that all histograms in A and B differ from each other, and all histograms in C differ from each other (Table S9).

In the histograms shown in Figures 8A and B a subset of the accuracy values fall below 0.5 for decoding runs with smaller numbers of ROIs (see for example Figure 8 Ai). As explained above and shown in Figure S10 Diii for these accuracy decreases the zΔF/F odorant period values overlap for many of the S+ and S- trials. At the same time, they diverge for one or two trials. However, for the odorant window a large number of decoding accuracy values remain above 0.65 for decoding with small ROI numbers. For example, for decoding with a single ROI 5.8% out of 6713 ROIs yields odor window accuracy above 0.65 (Figure 8 Ai). We quantified the divergence time for each decoding accuracy time course with odor window decoding accuracy >0.65. Divergence time was calculated after trial start when accuracy increased above 0.55 for at least 0.2 sec. Figure S11 shows the divergence times estimated using this method for all decoding accuracy time courses calculated with zΔF/F from single ROIs that yield odor accuracy above 0.65 for one session. Histograms of these divergence times are shown in Figure 8C for the different multiple ROI decoding sessions. The divergence times decrease as the number of ROIs used for decoding calculation increases (Figure 8C). A GLM analysis of divergence times yields significant differences for comparisons between accuracies calculated with different numbers of ROIs, p<0.001, 4 mice, 27354 observations, 27349 d.f., F-statistic 1400, p-value <0.001 (Table S9). Finally, Figure 8D shows all the decoding accuracy time courses with odorant decoding accuracy >0.65 calculated with a single ROI (Figure 8 Di) or all ROIs (Figure 8 Diii) sorted by divergence time. For the single ROI decoding there is a large variancein the start of the increase in accuracy ranging from trial start to reward delivery (Figure 8 Di). Computing average accuracy time courses for different divergence time windows shows that the onset of the increase in decoding accuracy is time-tiled and that decoding accuracy calculated for the earliest divergence time window is biphasic with a small increase at trial start followed by a larger increase after odor onset (Figure 8 Dii). These data show that the onset of increases in decoding accuracy, quantifying time of decision making, is time-tiled in hippocampal CA1 suggesting that these are decision-making time cells.

### Lick Decoding and Stimulus Decoding Differ in their Relationship to Lick Behavior

Next we asked how stimulus decoding prediction is related to lick behavior (quantified as lick fraction defined per time point as the fraction of trials when the animal was making contact with the lick tube). We expected to find differences between stimulus decoding and lick fraction because the accuracy of stimulus decoding starts increasing above shuffled at trial start before the odorant is presented (Fig. 6A), and, in contrast, licking starts diverging between S+ and S- after the odorant is delivered (Figs. S4A and 9A). In addition, we asked whether it was possible to decode lick behavior from neural activity and whether the relationship between lick prediction and lick fraction differed from the relationship between stimulus prediction and lick fraction. We performed this analysis separately for different trial types classified on behavioral outcome: correct responses (hits and correct rejections, CRs) and incorrect responses (miss and false alarms, FAs)(Figure 1A).

Figure 9A shows the time course for lick fraction for the four types of trials. There was a transient increase in lick fraction for all trial types when the animal started the trial by licking on the water spout (arrow in Figure 9A). For hits (red) this transient increase was followed by a steady increase in lick fraction during the odor period (0-4 sec), while for CR trials (blue) there was a decrease toward zero shortly after odorant presentation. In contrast, for miss trials, the lick fraction did not increase during the first second of the odorant period (cyan), and for FA trials, the lick fraction was high for the first half of the odorant period (magenta). Figure 9B is a bar graph showing the mean lick fraction during the first and second 2 sec response areas (RAs) during the odorant period (RAs are defined in Figure 1A). A GLM analysis yields significant differences in mean lick fraction for comparisons between CR or miss vs. FA or hit and for the interaction of miss or hit and FA and the RAs, p<0.001, 4 mice, 452 observations, 441 d.f., F-statistic 36.7, p-value <0.001 (Table S10).

**Figure 9.**
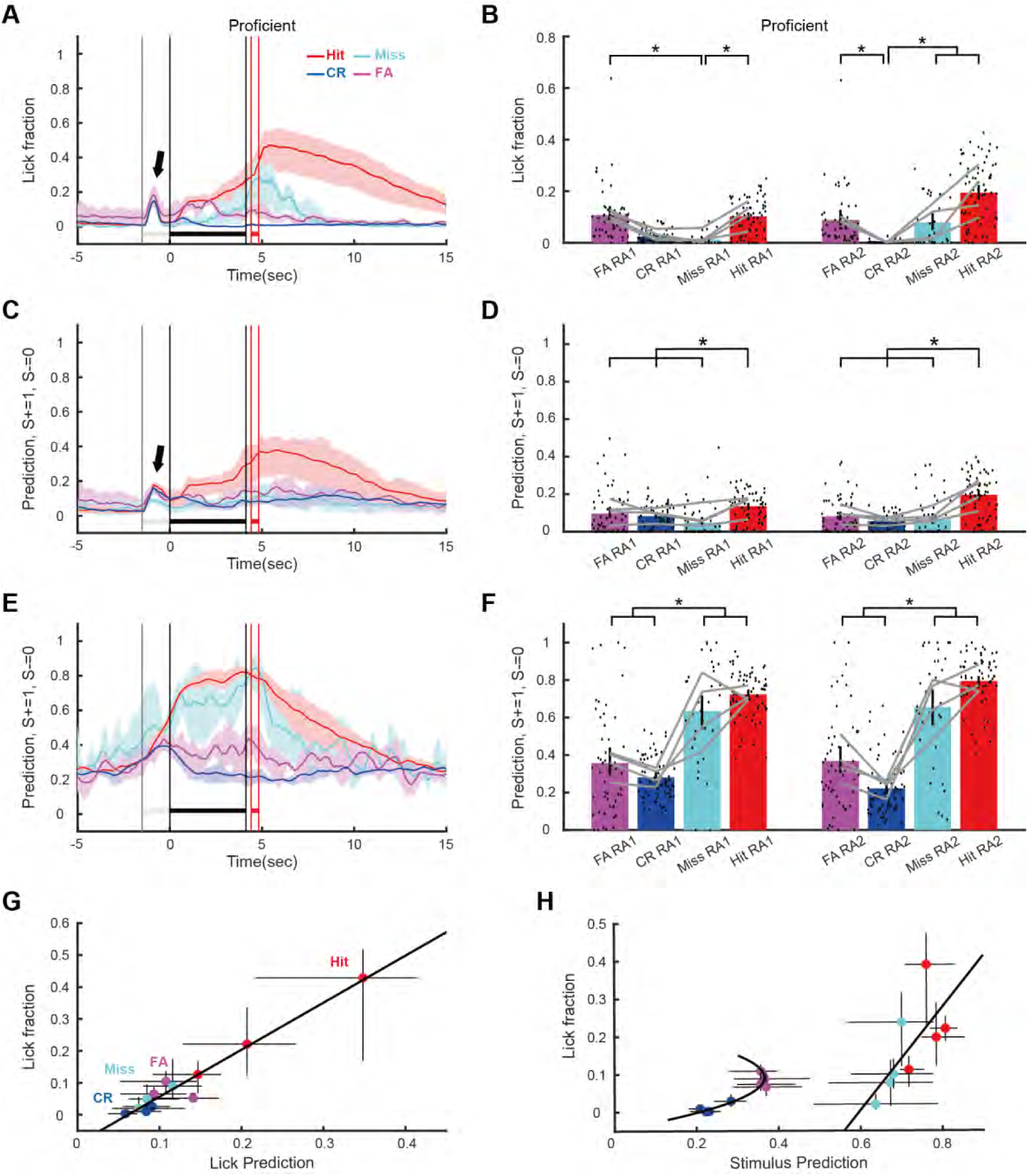
The relationship between lick fraction and prediction differs between stimulus decoding and decoding of lick fraction. (A) Mean lick fraction time course calculated for hits (red), miss (cyan), correct rejections (CR, blue) or false alarm (FA, magenta) trials for all sessions when the mice were proficient (66 sessions, 4 mice). (B) Bar graph quantifying mean lick fraction for proficient mice (66 sessions, 4 mice) in the two 2 second response areas of the odor period where the mouse must lick at least once to get water in a Hit trial (Figure 1A). Dots are per session lick fraction and grey lines are per mouse averages. (C) Mean prediction time course for lick fraction decoding within trials calculated for hits (red), miss (cyan), correct rejections (CR, blue) or false alarm (FA, magenta) trials for all sessions when the mice were proficient (66 sessions, 4 mice). (D) Bar graph quantifying mean lick fraction prediction for proficient mice (66 sessions, 4 mice) in the two 2 second response areas of the odor period where the mouse must lick at least once to get water in a Hit trial (Figure 1A). Dots are per session prediction and grey lines are per mouse averages. (E) Mean stimulus prediction time course within trials calculated for hits (red), miss (cyan), correct rejections (CR, blue) or false alarm (FA, magenta) trials for all sessions when the mice were proficient (66 sessions, 4 mice). (F) Bar graph quantifying mean stimulus prediction for proficient mice (66 sessions, 4 mice) in the two 2 second response areas of the odor period where the mouse must lick at least once to get water in a Hit trial (Figure 1A). Dots are per session prediction and grey lines are per mouse averages. The training period for the GLM decoding algorithm was 0.5 to 5.5 sec. (G) Relationship between mean lick fraction and mean lick fraction prediction (calculated with per mouse values). The bars are 95% CIs calculated per mouse. Lick fraction and lick fraction prediction were calculated in two second time windows spanning from odor onset to 6 sec after odor onset. The line is a linear fit. (H) Relationship between mean lick fraction and mean stimulus prediction (calculated from per mouse values). The bars are 95% CIs calculated per mouse. Lick fraction and stimulus prediction were calculated in two second time windows spanning from odor onset to 6 sec after odor onset. Lines are a linear fit of the data for Hit and Miss and a second order polynomial fit for CR and FA. For the graphs the data are shown separately for S+ Hits (red) and Miss (cyan) and S- CR (blue) and FA (magenta) trials. *p<0.05 for a pFDR-corrected t-test or ranksum tests, GLM statistics are in Table S10.

The accuracy for neural network decoding of lick fraction from zΔF/F for all ROIs per session when the mice were proficient (four mice, 66 sessions) is shown in Figure S12. In contrast with accuracy for stimulus decoding that starts diverging from shuffled at the trial start before the odorant is presented (Fig. 6A) lick decoding accuracy starts diverging from shuffled after the odorant presentation (Fig. S12A). The bar graph in Figure S12B shows the mean lick decoding accuracy data for pre (−1 to 0), odor (3.1 to 4.1) and reinforcement (4.5 to 5.5) time windows. A GLM analysis yielded a statistically significant difference for odor and reinforcement time windows vs. pre window and for the interaction between these time window comparisons and shuffling (p<0.001), 378 observations, 369 d.f., 4 mice, GLM F-statistic=62.1, p<0.001 (Table S10). Figures 9C and D show the lick prediction time course and mean lick prediction calculated in the RA windows for the different trial types (hit, miss, CR and FA). A GLM analysis yields significant differences in mean lick prediction for comparisons between hit or miss and FA and for the interaction of hit and FA and the RAs, p<0.001, 4 mice, 488 observations, 477 d.f., F-statistic 24.3, p-value <0.001 (Table S10).

Figure 9E shows the time course for stimulus prediction for the four types of trials. For hits (red) there was a steady increase in stimulus prediction that started at trial onset and leveled to a value of ∼0.8 during the odor period for hits (red) while for CR trials (blue) there was an initial increase after trial start that decreased to ∼0.2 during the odor period. For miss trials stimulus prediction appeared to increase to a value below hit prediction (cyan). For FA trials stimulus prediction started increasing at the trial start, but did not decrease to ∼0.3 during the odor period (magenta). Figure 9D is a bar graph showing the mean stimulus prediction during the first and second 2 sec response areas (RAs) during the odorant period. A GLM analysis yields significant differences in mean stimulus prediction for comparisons between CR, miss or hit, and FA, p<0.05, 4 mice, 452 observations, 441 d.f., F-statistic 57.3, p-value <0.001 (Table S10).

The time course for lick fraction (Figure 9A) appears to be similar to the time course for lick prediction (Figure 9C) and appears to differ from stimulus prediction (Figure 9E). In order to get information on whether there is a typical monotonic relationship between lick fraction and stimulus prediction, we plotted the mean lick fraction and mean stimulus prediction calculated during each second of the odor period. The relationship between lick fraction and lick prediction was linear (Figure 9G). In contrast, the relationship between lick fraction and stimulus prediction differed between S+ (hit/miss) and S- (CR/FA) (Figure 9H). For hits and miss the stimulus/lick fraction relationship was linear (Figure 9E, cyan and red circles), while for CR/FA points (Figure 9E, magenta and blue circles), the relationship appeared to be quadratic and the points did not fall along the hit/miss line. Therefore, the relationship between lick fraction and stimulus prediction in CA1 does not follow the simple linear relationship that is expected from neural activity closely related to motor action. Instead, the relationship differs strikingly for S+ vs. S- stimuli. However, decoding lick behavior from the same neural activity did yield a prediction that showed a linear relationship with lick fraction.

Finally, while decoding accuracy for stimulus decoding did not differ substantially between decoding algorithms (Figure S9), the algorithms that use nonlinear decoding (neural networks, NN and binary tree decision, BDT) performed substantially better than those that perform linear decoding (support vector machine, SVM, generalized linear model, GLM and linear discriminant analysis, LDA, Figure S12) suggesting that decoding of lick behavior involves nonlinear neural activity interactions in dCA1.

## DISCUSSION

In this study head-fixed mice engage in an associative learning go-no go task where they decide whether they lick to obtain a reward when presented with rewarded (S+) or unrewarded (S-) odorants at concentrations below irritant (trigeminal) threshold (Schaper, 1993). Unlike other sensory systems, the efferent axons from the olfactory bulb bypass the thalamus and synapse (directly or through relay in the piriform cortex) onto the LEC constituting a primary sensory input to dCA1 (Imai, 2014; Witter et al., 2017). Indeed, before it was realized that the hippocampus plays a crucial role in learning, Ramón y Cajal called the hippocampal formation the “quaternary” olfactory system because of this lack of relay through the thalamus (Cajal, 1901). Olfactory responses mediated by LEC input are particularly prominent in the mouse dCA1 (Igarashi et al., 2014; Manns et al., 2007; Wood et al., 1999), and odorant responses in dCA1 are known to be involved in tasks such as delayed match/non-match to sample (working memory)(MacDonald *et al*., 2013; Taxidis *et al*., 2020), odor place associative learning (Igarashi *et al*., 2014), odorant gradient navigation (Radvansky and Dombeck, 2018) and odorant sequence evaluation (Martinez et al., 2019).

Dorsal CA1 is arguably one of the most studied areas of the brain involved in learning and memory (Basu and Siegelbaum, 2015; Citri and Malenka, 2008; Fanselow and Dong, 2010) and it is well documented to play a role in spatial and episodic learning and memory (Eichenbaum, 2017; Moser *et al*., 1993). Yet, its role in go-no go associative learning has been called into question. Li et al. found that calbindin positive pyramidal neurons in CA1 became more selective for odorants as animals became proficient in the go-no go task and optogenetic inhibition of these cells slowed learning (Li *et al*., 2017). In contrast, a study by Biane and co-workers found that the accuracy of odor decoding in dCA1 was not altered by learning in the go-no go task (Biane *et al*., 2023). Here we find that there is a substantial increase in stimulus decoding accuracy when the animal becomes proficient in the go-no go task (Figure 6); that learning results in increased stimulus divergence of zΔF/F responses for single cells (Figure 4) and that learning elicits an increase in the stimulus divergence of stimulus components in demixed PCA (Figure 3). Our results are consistent with a role for dCA1 pyramidal cells in go-no go associative learning proposed by Li et al. and with local field potential recordings in dCA1 that find increased beta power (Martin et al., 2007) and increased theta-referenced beta and gamma power (Ramirez-Gordillo et al., 2022) upon learning in dCA1 in the go-no go associative learning task.

The go-no go task in this study also presents the animal with an auditory cue delivered by opening one of two odorant valves at the start of the trial, 1 to 1.5 sec before the odorant is delivered (Figures 1A and S3). Using dCA1 neural activity we could decode, albeit with low accuracy, this differential stimulus at the start of the trial (arrow in Figure 8Dii, Figure S10Ai). The hippocampus responds to auditory cues (Aronov et al., 2017). Interestingly, even though we find that the information on the odor valve click is represented by neural activity in dCA1, when we removed the odorants the behavioral performance of the mice dropped to 50% (Figure S2E-G). Therefore, although this auditory information is present in CA1, these animals do not use it as a stimulus cue in the go-no go task. In addition, for proficient mice, lick fraction prediction in error trials in lick decoding differs from prediction for stimulus decoding (compare Figures 9D and F, and Figures 9G and H). Indeed, prediction for stimulus decoding tends to represent the correct stimulus even when the animal makes a mistake (Figure 9F). In contrast, for lick decoding the predictions for errors align closer with the mistakes the animal makes in the lick response (compare Figure 9D with Figure 9B). Likely the animal does perceive the odorant, but is satiated and makes a mistake on purpose. These results indicate that CA1 encodes for different information manifolds that can be used differentially depending on the context and behavioral purpose.

We asked whether dCA1 pyramidal cells represent the stimulus identity (HEP vs. MO for odorant identity) or its valence (is this the rewarded stimulus?, S+ vs. S-). We found that predicting the stimulus decoded from zΔF/F ensemble neural activity in dCA1 was biased towards the unrewarded stimulus (S-) between trials (Figure 7D). This is unlike stimulus identity decoding in areas such as the auditory cortex, where there is no bias in stimulus prediction (Libby and Buschman, 2021) suggesting that dCA1 ensemble activity represents stimulus value (S+ vs. S-) as opposed to odorant identity (HEP vs. MO). Indeed, when the stimuli were reversed the responses were consistent with representation of valence for a large portion of the ROIs (Figure 5). This result is consistent with valence representation found in olfactory bulb single units after reversal in the go-no go learning task (Doucette and Restrepo, 2008; Li et al., 2015), functional plasticity of sensory inputs elicited by learning in the olfactory bulb (Abraham et al., 2014; Chu et al., 2016), and with decoding of stimulus identity after stimulus reversal for measurements of theta-referenced beta and gamma LFP power in dCA1 (Ramirez-Gordillo *et al*., 2022). Thus, pyramidal cells represent both odorant valence and odorant identity.

Time-tiled odorant responses by “time-cells” in dCA1 are thought to be involved in delayed match/non-match to sample where they presumably provide short term working memory of the identity of the first odorant that is presented in the task (MacDonald *et al*., 2013; Taxidis *et al*., 2020). Interestingly, we find that for the go-no go task the divergence between S+ and S- stimuli in zΔF/F odorant responses of individual pyramidal cells takes place at different times between trial start and reinforcement (Figure 4) and, consistent with these results, stimulus decoding accuracy decoded from single neuron activity starts diverging from accuracy computed from shuffled stimulus labels (50%) at discrete times ranging from trial start to several seconds after reinforcement (Figures 8, S10 and S11). Therefore, decision making for the difference between stimuli based on single cell dCA1 neural activity takes place at different discrete times in dCA1. This finding of “decision-making time-cells” is novel and complementary to stimulus time-cells engaged in delayed match/non-match to sample tasks (MacDonald *et al*., 2013; Taxidis *et al*., 2020). We hypothesize that this discrete time tiling of information for decision making in dCA1 by sequential neural dynamics plays a role in go-no go associative learning by representing a memory of whether the rewarded odorant is presented to the animal that the mouse uses to decide whether it should sustain licking to get a reward. Whether this is the case will be determined in future studies.

Finally, our data raise the question of the specific role of sequential activity in associative learning in dCA1. On one hand, sequential activity in dCA1 is often associated with transition in the state space where each state encodes for a predictive representation of future states given the current state (Nieh *et al*., 2021; Stachenfeld et al., 2017). Under this interpretation, time-tiling represents state transition in a cognitive map representing the temporal task structure. On the other hand, sequential activity potentially encodes a single working memory that makes the animal more likely to choose a specific outcome as time passes. Previous modeling work revealed that depending on the model configuration, a recurrent neural network learns to represent working memory either by sequential or persistent activity (Orhan and Ma, 2019). It indicates that time-tiling might encode a binary decision (go or no-go) with spatial-temporal patterns. Sequential working memory representation is also beneficial for binding multiple information, such as stimulus identity, valence, and context, into a single working memory (Feldman, 2013; Hiratani and Sompolinsky, 2023). Future work will clarify whether time-tiling in the decision period represents a transition in the cognitive map or a go/no-go decision variable.

## ACKNOWLEDGEMENTS

We would like to thank Brooke Baxter for technical help and Kira Steinke for discussions and feedback on the manuscript. This research was supported by the US National Institutes of Health (NIH UF1 NS116241 and NIH R01 DC000566), and the National Science Foundation (NSF BCS-1926676).

## CONTRIBUTIONS

M.M., F.S.de S., E.G. and D.R. conceived and designed the experiments, F.S.de S. performed the surgeries, acquired experimental data and did data analysis, M.M. performed experiments and data analysis, E.G. and G.F. designed and implemented optical instrumentation, G.F. provided technical assistance with multiphoton microscopy, S.A. and D. T. performed sound recordings and analysis, J.R. assisted with immunohistochemistry, A.G.-P. provided technical help, generated illustrations and did immunohistochemistry, J.P. generated code for NWB data sharing, N.H. provided advice on data analysis and writing, M.M., F.S.de S. and D.R. wrote the manuscript and all authors edited the manuscript.

## STAR * METHODS

### Key resources table

| REAGENT or RESOURCE | SOURCE | IDENTIFIER |
| --- | --- | --- |
| **Antibodies** |  |  |
| Donkey anti-rabbit 594 secondary antibody | Jackson Immunoresearch | 11-585-152 |
| Rabbit anti GFAP antibody | Abcam | ab7260 |
| **EXPERIMENTAL MODELS: ORGANISMS/STRAINS** |  |  |
| Thy1-GCaMP6f mouse, C57BL/6J-Tg(Thy1-GCaMP6f)GP5.17Dkim/J | The Jackson Laboratory | 025393 |
| **SOFTWARE AND ALGORITHMS** |  |  |
| Demixed PCA | (Kobak *et al.*, 2016) | http://github.com/machenslab/dPCA |
| Matlab 2022b and 2023a | https://www.mathworks.com/products/matlab.html | N/A |
| **OTHER** |  |  |
| GRIN lenses | GRINTECH GmbH | NEM-050-25-10-860-DS |
| Metabond | Thermo Fisher | NC0877382, NC2048900, NC2048904 |
| Paraformaldehyde | Electron Microscopy Sciences | 15714-S |
| DAPI | Thermo Fisher | 62248 |
| Buprenorphine SR/ER | Zoopharm/Wedgewood |  |
| Vetbond | MWI | 006245 |
| Light mineral oil | Fisher | O121-1 |
| Heptanal | Sigma-Aldrich | H-6129 |
| OCT | Sakura Finetech | 4583 |

### Animals

All experimental protocols were approved by the Institutional Animal Care and Use Committee of the University of Colorado Anschutz Medical Campus, in accordance to NIH guidelines. Mice were bred in the animal facility, and we used both males and females, a total of four adult Thy1-GCaMP6f mouse (Jax 025393 C57BL/6J-Tg(Thy1- GCaMP6f)GP5.17Dkim/J) for head-fixed awake behaving two-photon imaging of CA1 pyramidal cells through a GRIN lens. We used the 5.17 strain of Thy1-GCaMP6f mice because of their strong expression of GCaMP6f in CA1 (Dana et al., 2014). The animals were housed in a vivarium with a 14/10 h light/dark cycle. Food was available ad libitum. Access to water was restricted in behavioral training session according to protocols. All mice were weighed daily and received sufficient water during behavioral sessions to maintain >80% of their original body weight.

### Surgical Procedures

Mice were injected with carprofen (5 mg/kg, IP) one hour before surgery and were anesthetized with ketamine/xylazine (IP, 100 mg/kg and 10 mg/kg, respectively). Anesthetic redosing was assessed by pinching the hind feet every 5 min. A craniotomy of diameter ∼1.2 mm was made over the right hippocampus and a GRIN lens of 4 mm length and 1 mm diameter (GRINTECH, NEM-050-25-10-860-DS) was implanted at 1.8 mm lateral and 2.4 mm caudal of Bregma and 1.25 mm below dura (Radvansky and Dombeck, 2018). The GRIN lens edges were sealed with Vetbond glue (3M, USA) and a custom-made steel head bracket was glued to the skull with Metabond (Parker, USA) for head-fixing and imaging. The mice were treated with buprenorphine SR/ER for two days after surgery (0.05 mg/kg, SQ). dCA1 imaging was initiated two weeks after surgery.

### Immunohistochemistry and CT-Scan

Immunohistochemistry was performed after animals finished all the training to visualize GCaMP6f expression in the hippocampus CA1 region and the track of the GRIN lens implant. Mice were sacrificed and transcardially perfused with ice cold 4% paraformaldehyde (PFA), followed by equilibration in 30% sucrose. After the brain sank, the tissue was embedded in optimal cutting temperature (OCT) compound and was frozen at -80°C. Slices of 60 μm thickness were cut with a Leica cryostat. The slices were stained with rabbit anti-GFAP primary antibody (Ab7260, Abcam, USA) and Alexa Fluor 594 donkey anti-rabbit secondary antibody (11-585-152, Jackson Immunoresearch, USA). The slices were counterstained with DAPI (Thermo Fisher Scientific, USA) and imaged using a confocal laser scanning microscope (Nikon A1R, Japan). To further verify the 3D position of the GRIN lens in the skull and brain, we performed microCT-Scan recording. We perfused the mice with PBS and the ex-vivo PFA fixed head was used for acquiring the microCT imaging on a Siemens Inveon microCT system (Siemens, Germany).

### Go-No Go training

Mice were water deprived by restricting daily water consumption to 1-1.5 ml. Mice were monitored for signs of dehydration or decreased body weight below 80% of the initial weight. If either condition occurred, the animals received water ad-lib until they recovered. When the animals were water-deprived, they were trained in a head-fixed olfactory go-no go task with 1% HEP vs MO odorant application (Sigma-Aldrich, USA)(Gire et al., 2013; Li *et al*., 2015; Ma *et al*., 2020). An olfactometer controlled valves to deliver a 1:40 dilution of odorant at a rate of 2 L min^-1^. An Intan RHD2000 acquisition system (Intan, USA) recorded licks measured by monitoring the resistance between the lick spout and the floor of the olfactometer. The water-deprived mice started the trial by licking on the water port. The odorant was delivered after a random time interval ranging from 1 to 1.5 seconds. In S+ trials, the mice needed to lick at least once in two 2 sec lick segments to obtain a reward (0.1 g ml^-1^ sucrose water) (Figure 1A). In S- trials, the mice need to refrain from licking in one of the two 2 sec segments to avoid a longer inter-trial interval (+ 10 sec). The animal’s behavior performance was evaluated in a sliding window of 20 trials and the calculated value was assigned to the last trial in the window. Therefore, it estimated the performance in the last 20 trials. The percent correct value represents the percent of trials in which the animal successfully performed appropriate actions (100*(Hits+CRs)/number of trials), and we considered the animal proficient if the percent correct performance is above 80%. In reverse go-no go training sessions the S+ and S- odorants were switched after reaching a proficient level in forward training.

### Acoustic recordings of auditory cues at the start of the trial in the go-no go task

In addition to the odorant cue the olfactometer emits a click at the start of the trial when one of the two odorant valves opens 1-1.5 sec before odorant onset. Acoustic recordings of the olfactometer were made using a GRAS 46BH-1 omnidirectional microphone and GRAS 26AC-1 preamplifier with Type 12AA power module, M-Audio Pro interface, and Audacity software. The microphone was positioned to approximate the location of the mouse’s head during experiments. Recordings were time-aligned to experimental hardware using a trigger pulse. Recordings were windowed 7 seconds prior to 15 seconds following stimulus offset and separated according to S- or S+ conditions in MATLAB. Spectrograms of each stimulus were then generated to identify auditory events.

Figures S3B-C show a sound spectrogram of the click recorded with the ultrasound microphone at the location of the mouse’s head in the two-photon microscope aligned with the opening of the odorant valve. Figure S3D shows that the identity of the odorant valve (S+ vs S-) can be decoded from the sound click using a generalized linear model (GLM) decoding algorithm, albeit with low accuracy (∼ 65%) (ranksum test p<0.05, 58 trials for decoding stimuli vs. 290 trials for shuffled stimulus decoding).

While this task provides both olfactory and sound sensory inputs, the mouse cues on the olfactory stimulus as shown by the fact that percent correct behavior decreases close to 50% when the odor cues are removed while the valve opening (and consequent sound) are not altered (Figure S2E-G, consistent with previous studies with the same olfactometer (Slotnick and Restrepo, 2005)).

### Two-photon imaging of dorsal hippocampus CA1 activity in animals undergoing the Go-No Go task

All the animals were first habituated to the setup to minimize stress during the imaging experiments. All the imaging sessions started at least 10 minutes after the mice had been head-fixed. The head fixed two-photon imaging system consisted of a movable objective microscope (MOM, Sutter Instrument Company, USA) paired with a 80 MHz, ∼100 femtosecond laser (Mai-Tai HP DeepSee, Spectra Physics, USA) centered at 920 nm. The MOM was fitted with a single photon epifluorescence eGFP filter path (475 nm excitation/500-550 nm emission) used for initial field targeting followed by switching to the two-photon laser scanning path for imaging GCaMP6f at the depth of the CA1 cell body layer. Either the galvometric laser scanning system or a resonant scanner was driven by SlideBook 6.0 (Intelligent Imaging Innovations, USA). The two-photon time lapses were acquired at 395 x 380 pixels using a 0.4 NA/10x air objective (Olympus, Japan) at either 5.3 Hz (galvo) or 20 Hz (resonant). On the day of initial imaging, a FOV was selected to image a large number of active hippocampal CA1 neurons through the GRIN lens, and GCaMP6f movies for several sessions (12-40 trials each) were collected each day. After two-photon imaging a second image of the vasculature was captured with wide field epifluorescence to reconfirm the field.

### Statistical analysis

Statistical analysis was performed in Matlab R2022b or R2023a (Mathworks, USA). Statistical significance for changes in measured parameters for multiple factors such as learning and odorant identity (S+ vs. S-) was estimated using a generalized linear model (GLM)(Yu et al., 2022) with post-hoc comparisons performed either with a two-sided t test, or a ranksum test, depending on the result of an Anderson-Darling test of normality. Post-hoc tests were corrected for multiple comparisons using the false discovery rate p-value (pFDR) (Curran-Everett, 2000). Asterisks in bar graphs specify statistically significant differences (p<pFDR) on post-hoc tests. Not all statistically significant differences are shown with asterisks because showing all significant differences would make the figures too complicated. However, all pairwise post-hoc test p-values and pFDR are shown in the supplementary tables. Finally, we provide 95% confidence intervals (CIs)(Halsey et al., 2015) estimated by bootstrap analysis of the mean by sampling with replacement 1000 times using the bootci function in MATLAB (shown in the figures as vertical black lines for bar graphs or shade bounding of mean value lines for time courses).

### Data analysis

Raw imaging data was first surveyed in ImageJ 1.52 (NIH, USA) to exclude image sequences exhibiting axial movement. We did not find evidence of axial movement while the animal was engaged in the go-no go task. The raw imaging data was processed with DeepCAD-RT to denoise time-lapse images (https://github.com/cabooster/DeepCAD-RT)(Li *et al*., 2023), and then motion corrected with NoRMCorre algorithm (Pnevmatikakis and Giovannucci, 2017). The nonegative fluorescence traces were generated with EXTRACT (https://github.com/schnitzer-lab/EXTRACT-public) (Hakan *et al*., 2021) with appropriate parameters. After Extract analysis, the ΔF/F traces of the spatial components were sorted and we assigned trial traces to different behavioral events (Hit, CR, Miss and CR) and aligned them to trial start, odorant onset or reinforcement delivery for further analysis. Data will be We will deposit the data in NWB format in DANDI. We will add the following statement to the manuscript: Data and associated metadata were uploaded to the DANDI archive [RRID:SCR_017571] using the Python command line tool (https://doi.org/10.5281/zenodo.7041535). The data were first converted into the NWB format (https://doi.org/10.1101/2021.03.13.435173) and organized into a BIDS-like (https://doi.org/10.1038/sdata.2016.44) structure.

Data analysis was performed using custom code in Matlab R2022b or R2023a (https://github.com/restrepd/CaImAnDR). Four mice were trained in 3-6 training sessions per day. Analysis was performed per session and classified into either the learning stage with percent correct behavior within the 45-65% range (blue in Figure S2, 20 sessions) or the proficient stage with percent correct behavior in the 80-100% range (red in Figure S2, 55 sessions). For each ROI we z-scored ΔF/F by dividing by the standard deviation of ΔF/F calculated over the entire session (we call this z scored ΔF/F zΔF/F). Below we describe the different data analysis methods.

#### Dimensionality

Following Litwin-Kumar et al. (2017) we defined the dimension of the system (dim) with *M* inputs as the square of the sum of the eigenvalues of the covariance matrix of the measured ΔF/F for all ROIs in the FOV divided by the sum of each eigenvalue squared:

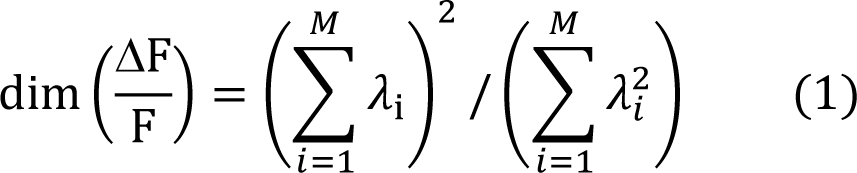

where λ_i_ are the eigenvalues of the covariance matrix of ΔF/F computed over the distribution of ΔF/F signals measured in the FOV. If the components of ΔF/F are independent and have the same variance, all the eigenvalues are equal and dim(ΔF/F) = *M*. Conversely, if the ΔF/F components are correlated so that the data points are distributed equally in each dimension of an *m*-dimensional subspace of the entire *M*- dimensional space, only *m* eigenvalues will be nonzero and dim(ΔF/F) = *m*.

#### Demixed principal component analysis

Demixed principal component analysis (dPCA) is a dimensionality reduction method decomposes population activity into components related to task parameters such as stimuli, decisions, or rewards (Kobak *et al*., 2016). dPCA was calculated for S+ or S- trials using code provided by Kobak et al. modified to yield stimulus and independent components using data from all ROIs from all sessions belonging to either the learning or proficient stages.

#### Divergent cell detection and analysis

Whether a cell’s zΔF/F diverged between S+ and S- (Figures 4 and 5) was determined as follows: Only sessions with a minimum of 12 trials were included. ROIs with calcium spikes in fewer than 25% of the trials were excluded. For the rest of the ROIs the zΔF/F in the session was considered to be divergent if the GLM p-value for S+ vs. S- was below the pFDR. Examples of the time course zΔF/F for divergent ROIs are shown in Figure S6A.

As shown in Figures 4A, 5A, 5B and S8A, the per trial time courses for zΔF/F were heterogeneous. In order to quantify the heterogeneity we calculated within-trial cross-correlation coefficients between all divergent zΔF/F time courses (including both S+ and S- in the calculation) for all divergent ROIs (e.g. Figure 4B). The zΔF/F time courses for the different ROIs were then separated into different clusters by estimating an agglomerative hierarchical cluster tree using the linkage function of MATLAB. The number of clusters was specified arbitrarily.

In order to quantify the divergence in zΔF/F between HEP and MO trials for Figure S8 we calculated d’, which is a measure of the difference in between two distributions.

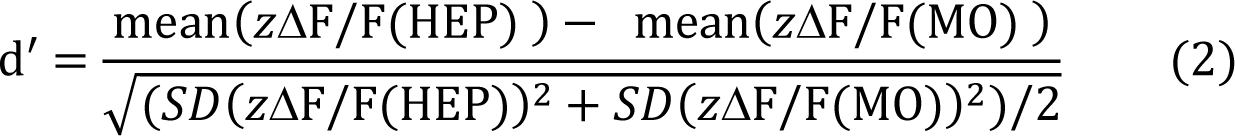

where SD is the standard deviation. As done for zΔF/F time courses we used cross-correlation coefficients and an agglomerative hierarchical cluster tree to sort the time courses shown for d’ in Figure S8C,D.

#### Stimulus decoding

Stimulus decoding was performed using GLM (fitglm in MATLAB). The algorithm was trained per session for all time points and all trials within time periods and number of ROIs per session was indicated in the text. Only sessions with 16 or more trials were processed. The predicted stimulus was assessed using leave one trial out, and winner takes all procedures. We report the results of GLM decoding, but we obtained similar results with linear discriminant analysis, neural network decoding, binary decision tree decoding and support vector machine decoding (Figure S9). For GLM stimulus decoding performed with a subset of 2, 5 or 15 ROIs (Figure 8) decoding was performed in separate runs with 40 unique subsets of ROIs drawn randomly from the total number of ROIs per session.

#### Lick fraction decoding

Lick fraction decoding was performed using a neural network classification model (fitcnet in MATLAB). The algorithm was trained per session for one second windows covering from -5 to + 10 sec in each trial. Only sessions with 16 or more trials were processed. The predicted stimulus was assessed using leave one trial out and winner takes all procedures.

*Analysis of prediction replay of S+ prediction for stimulus decoding between trials* Sudden changes in stimulus prediction from S- to S+ between trials were determined in decoding prediction moving window averages (with windows of 10 time points) such as the prediction time course shown in Figure 7A. We searched for a prediction replay (arrow in Figure 7A) by searching for a sudden shift in prediction from below the five percentile of the shuffled stimulus control for decoding prediction (lower edge of the blue shade in Figure 7A) to above 95 percentile for the shuffled stimulus decoding prediction (upper edge of the blue shade in Figure 7A). We compared the average time courses for these sudden between-trial increases to S+ (blue line in Figure 7E) to the time course of decoding prediction centered in the middle of adjacent between-trial intervals where we did not find a sudden increase to S+ (magenta line in Figure 7E).

---

**Figure S1.**
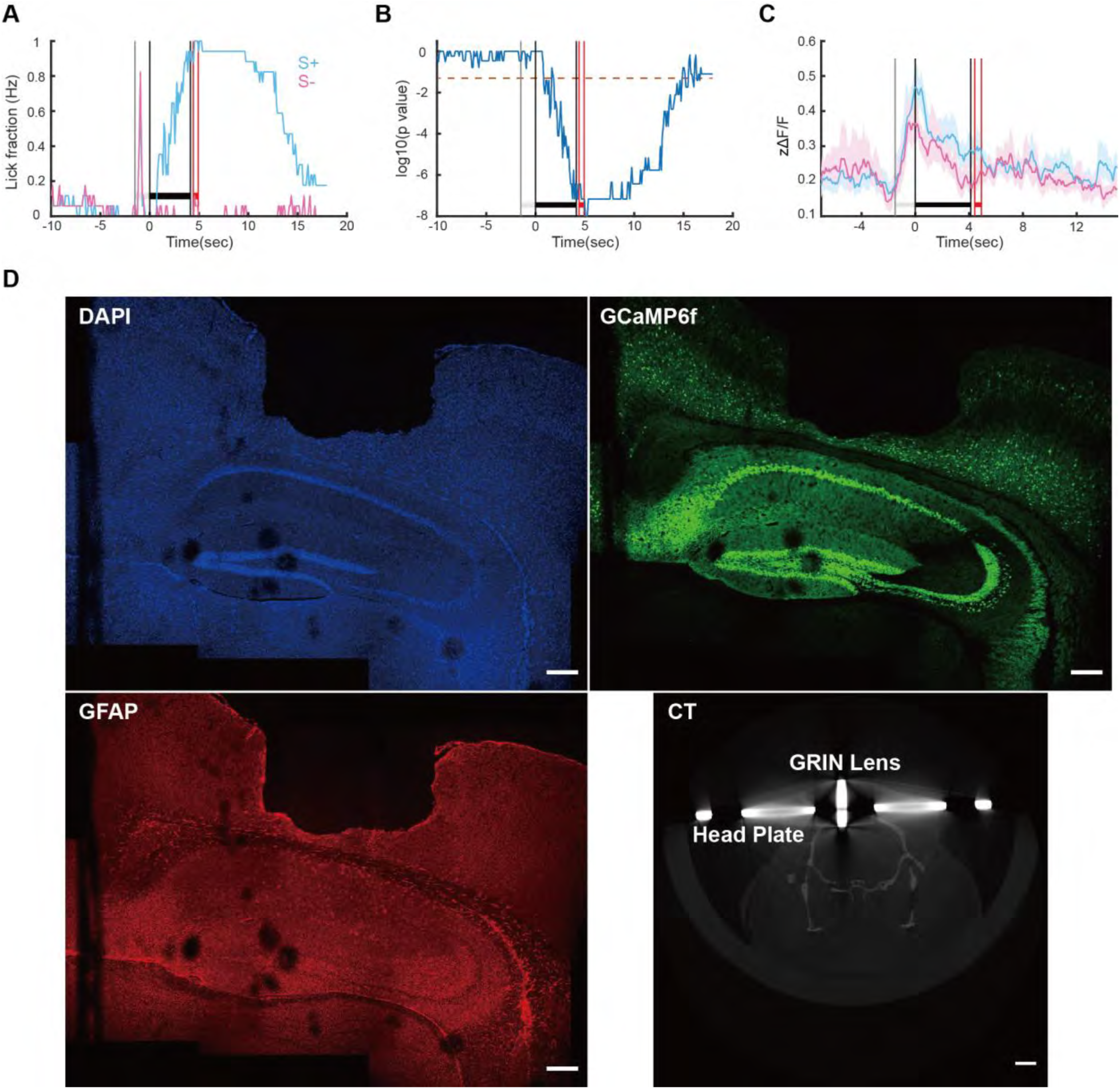
Supplementary data for Figure 1. (A) Mean of lick fraction time course for S+ and S- trials for the session for the data shown in Figure 1. (B) p-value for a ranksum test comparing S+ vs. S- per trial lick time courses for the session for the data shown in Figure 1. The horizontal red line is p=0.05. (C) Mean of per trial zΔF/F time course for S+ and S- trials for the session for the data shown in Figure 1. (D) Immunofluorescence and CT scan showing the location of the GRIN lens in one mouse. Scale bars are 200 μm for the immunohistochemistry and 2 mm for the CT scan.

**Figure S2.**
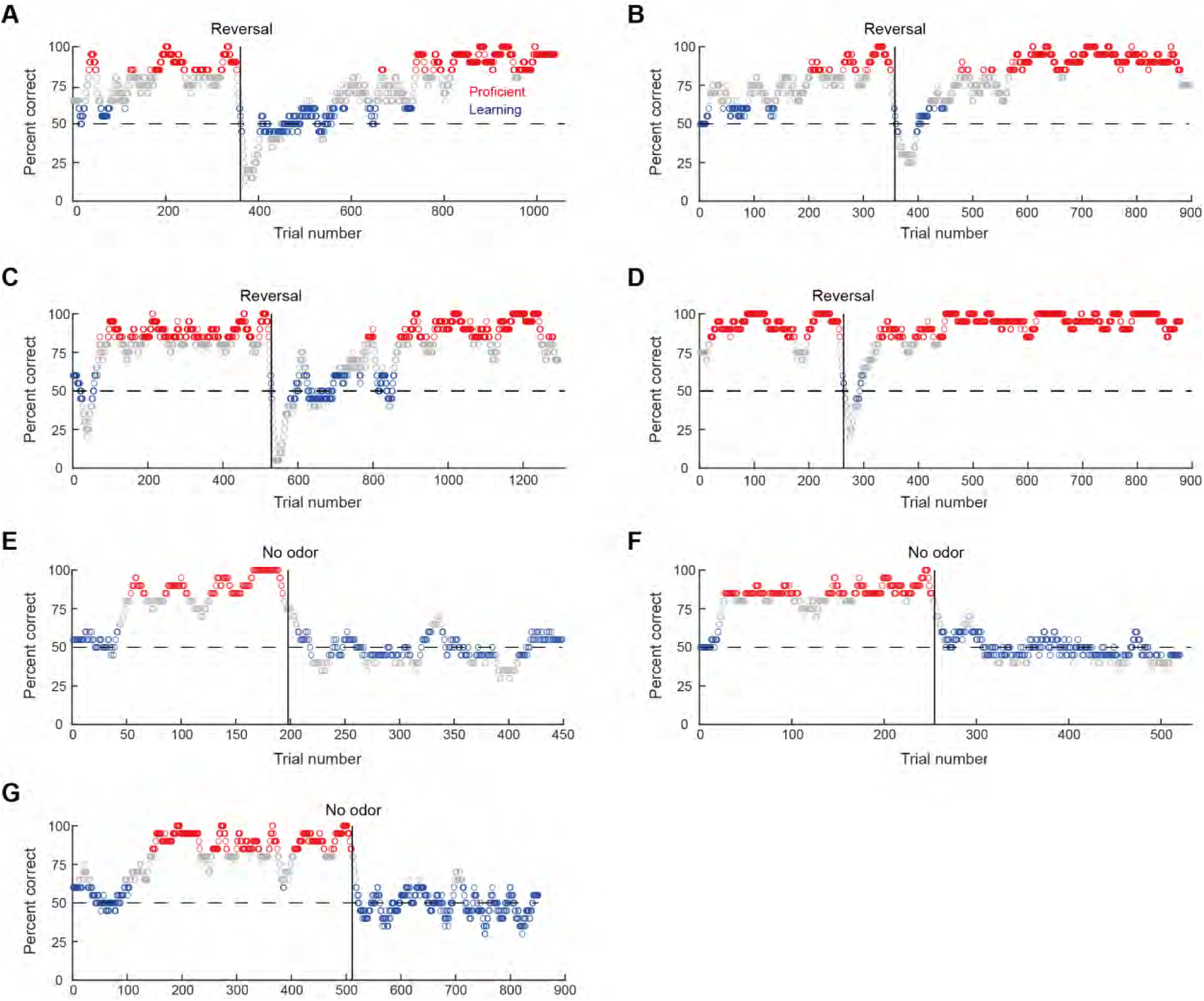
Behavioral performance of mice in the go-no go task. (A to D) show the percent correct behavior calculated in a moving window of 20 trials for the entire training period for the four mice used in this study. Percent correct behavior is calculated as 100*(Hit trials+CR trials)/20. The trials with performance between 45 and 65 % correct (shown in blue) were classified as learning stage trials while the trials with performance at or above 80 % correct were classified as proficient (red). The task was started with heptanal (HEP) as S+ and mineral oil (MO) as S- (forward sessions). The vertical black line shows the point where these stimuli were reversed (MO became S+ and HEP became S-, reversed sessions). (E, F and G) show for three mice that the behavioral performance drops to 50% when the odorants are not delivered, but the odorant delivery valves still produce a click sound at the start of the trial (vertical line, no odor).

**Figure S3.**
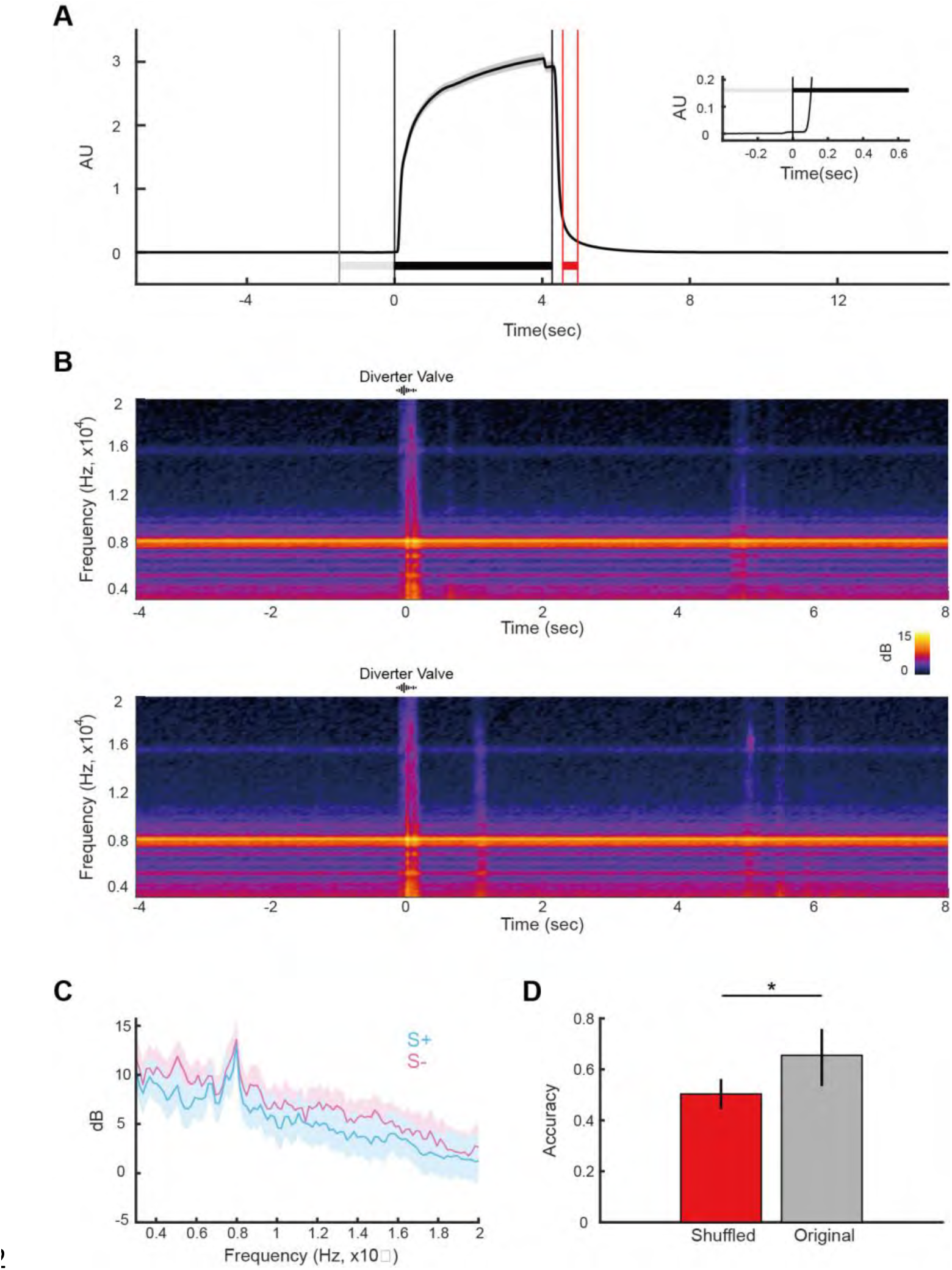
Time course for odorant onset and auditory cues. (A) Increase in odorant concentration measured with a photoionization detector (PID). The inset shows that the onset of the increase in odorant concentration takes place ∼100 msec after the diverter valve is turned off to deliver the odorant to the nose cone (time zero). (B) Spectrogram time course for the mean sound power (in dB) for the clicks produced by the olfactometer for S+ and S- trials measured at the location where the mouse’s head is in the 2P imaging enclosure. Time is referenced to the time point where the diverter and odor valves are turned on simultaneously (the start of the trial). Sound recording was performed in 30 S+ and 28 S- trials. The average power before diverter valve activation was subtracted from the power at each time point. (C) Spectrogram for the mean sound power (in dB) measured at the time bin when the click occurred for the clicks produced by the olfactometer for S+ and S- trials measured at the location where the mouse’s head is in the 2P imaging enclosure. (D) Accuracy for decoding of trial type (S+ or S-) from sound power measured at the click. Decoding of trial type from sound power for clicks recorded in 30 S+ and 28 S- trials was performed using GLM with a leave one out procedure. As a control the labels (S+ or S-) were shuffled. A ranksum test yields a p-value of 0.01 for the difference in per trial label prediction between shuffled and original (no shuffling) (n=58 trials).

**Figure S4.**
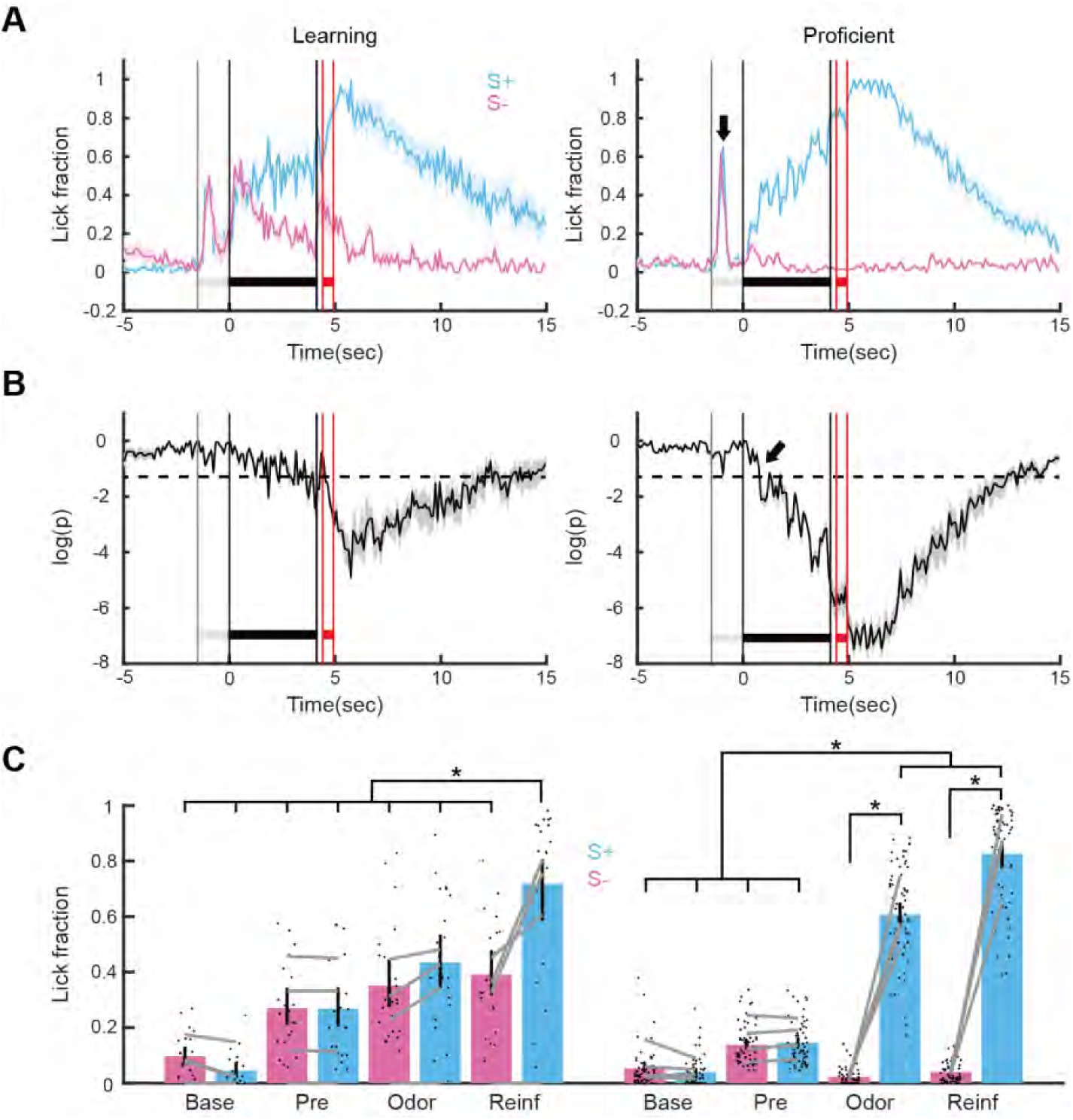
Lick responses for learning and proficient stages in the go-go task. (A) Time course for lick fraction mean ± 95% CI for S+ and S- trials for all learning and proficient sessions (4 mice). Left: learning stage, right: proficient stage. Arrow points to the increase in lick fraction that takes place at the start of the trial. (B) p-value (mean ± 95% CI) calculated by a per time point ranksum test of the difference in lick fraction between S+ and S- trials (evaluated per session). The p-value was calculated for all learning and proficient sessions (4 mice). The horizontal dashed line is p=0.05. The arrow points to the time when the p-value decreases monotonically below 0.05 for proficient mice. (C) Bar graph showing the mean value of lick fraction calculated in the following time windows: Base: -4.5 to -3.5 sec, Pre: -1 to 0 sec, Odor: 3.1 to 4.1 sec, Reinf: 4.5 to 5.5 sec. Dots are lick fraction values per session and grey lines are lick fraction values per mouse. *p<0.05 for a pFDR-corrected t-test or ranksum tests, GLM statistics are in Table S1.

**Figure S5.**
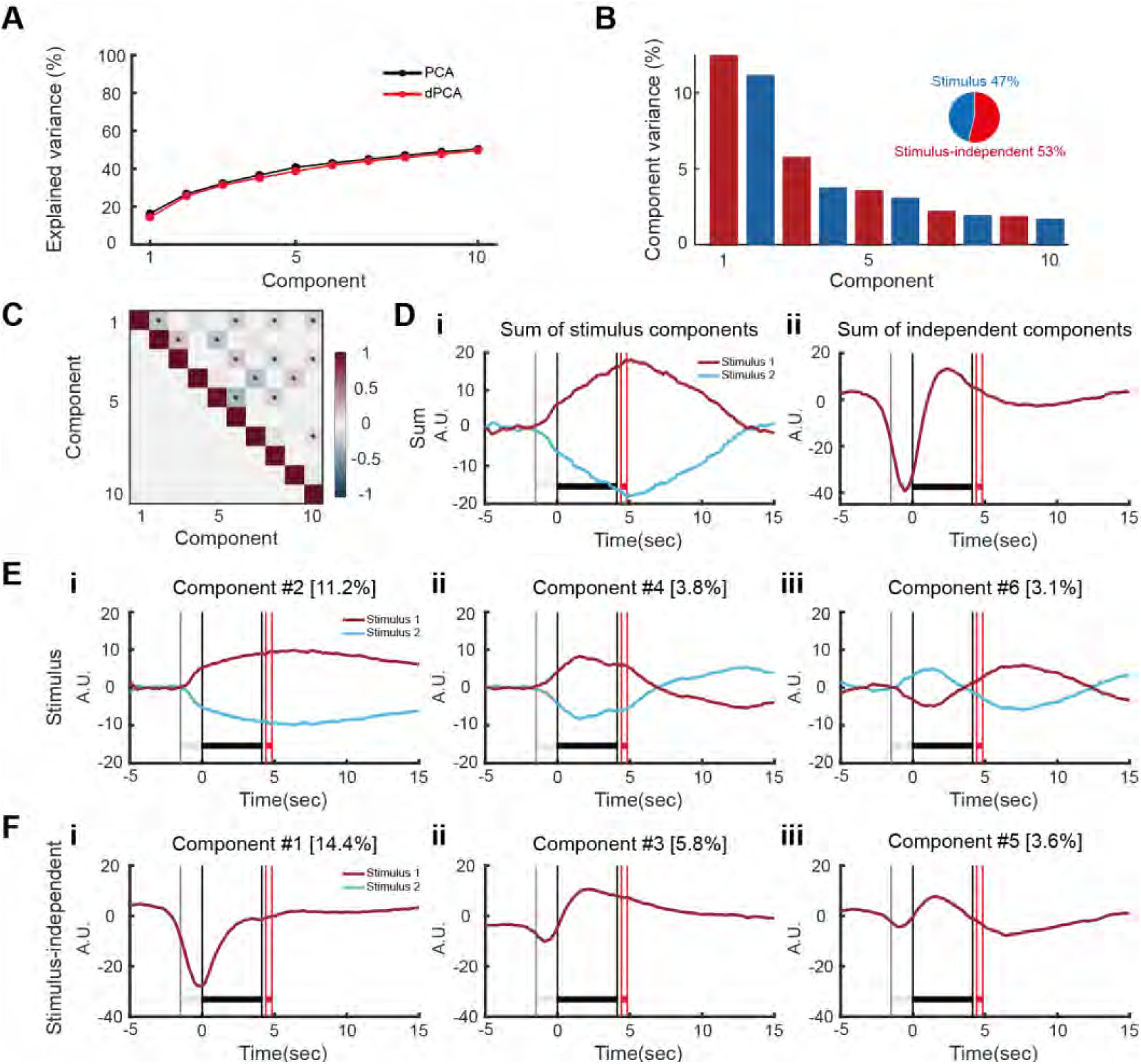
Example of demixed PCA for one mouse. (A) Cumulative variance explained by PCA (black) and dPCA (red). Demixed PCA explains almost the same amount of variance as the standard PCA. (B) Variance of the individual demixed principal components. Each bar shows the proportion of total variance, and is composed out of two stacked bars of different color: blue for stimulus variance and red for stimulus independent variance. Pie chart shows how the total signal variance is split among parameters. (C) Upper-right triangle shows dot products between all pairs of the first 15 demixed principal axes. Stars mark the pairs that are significantly and robustly non-orthogonal (see Materials and methods). Bottom-left triangle shows correlations between all pairs of the first 15 demixed principal components. (D) Time course for the sum of all stimulus (i) or independent (ii) dPCA components (E) Time course for the three top stimulus dPCA components (F) Time course for the three top independent dPCA components

**Figure S6.**
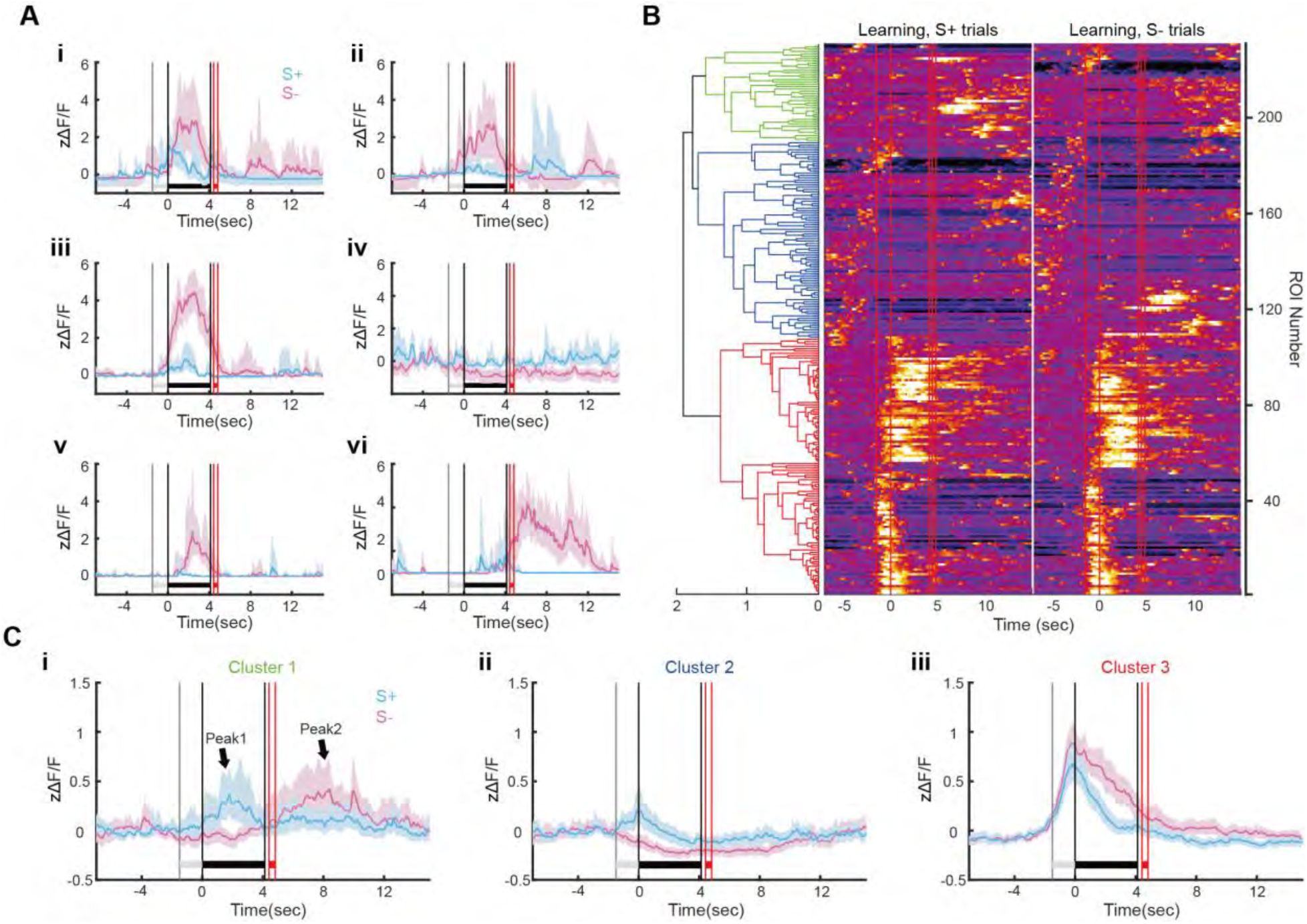
Stimulus-divergent zΔF/F responses for the learning stage and examples of zΔF/F responses that diverge between S+ and S- at different discrete time points after the start of the trial. (A) Examples of zΔF/F time courses for single ROIs that differ in onset time for divergence between S+ and S- trials (B) zΔF/F time courses for single ROIs that were divergent between S+ and S- trials in the learning sessions (18 sessions, 4 mice). Time courses were sorted by estimating an agglomerative hierarchical cluster tree shown on the left that was calculated using the cross-correlation coefficients between all divergent zΔF/F time courses (not shown). The red vertical lines show (in order): trial start, odorant on, odorant off, reinforcement on and reinforcement off. (C) Mean zΔF/F time courses for the three clusters in the hierarchical tree shown in B.

**Figure S7.**
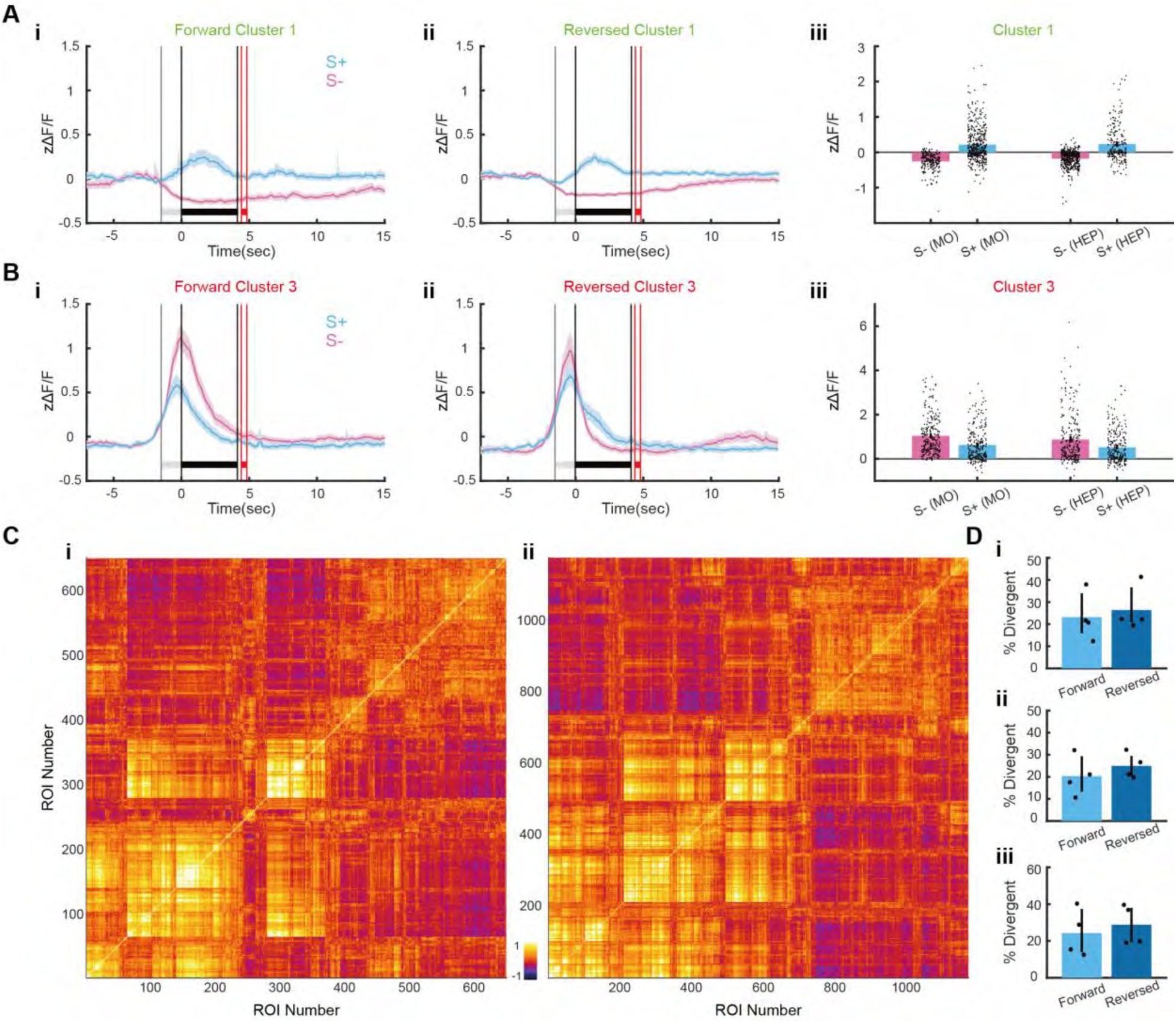
Supplemental data for Figures 5A and B characterizing changes in stimulus divergent zΔF/F responses after reversal of odorant valence. (A and B) Mean zΔF/F time courses (i,ii) and peak zΔF/F (iii) for clusters 1 and 3 of the hierarchical cluster trees shown in Figures 5 A and B. Mean zΔF/F time courses for cluster 2 is Figure 5C. Table S6 shows the GLM statistical analysis for iii. (C) Cross-correlation coefficients computed between all per trial zΔF/F time courses shown in Figures 5A and B. The coefficients were sorted by the agglomerative hierarchical cluster tree shown in those figures. (D) i. Percent divergent ROIs per mouse for the data in Figures 5A and B. ii and iii. Percent responsive ROIs per mouse for S+ and S- trials. The percent divergent (i) and percent responsive (ii and ii) differences between learning and proficient stages were not statistically significant (two tailed t test p>0.05, n=3 for learning and 4 for proficient, 5 d.f.).

**Figure S8.**
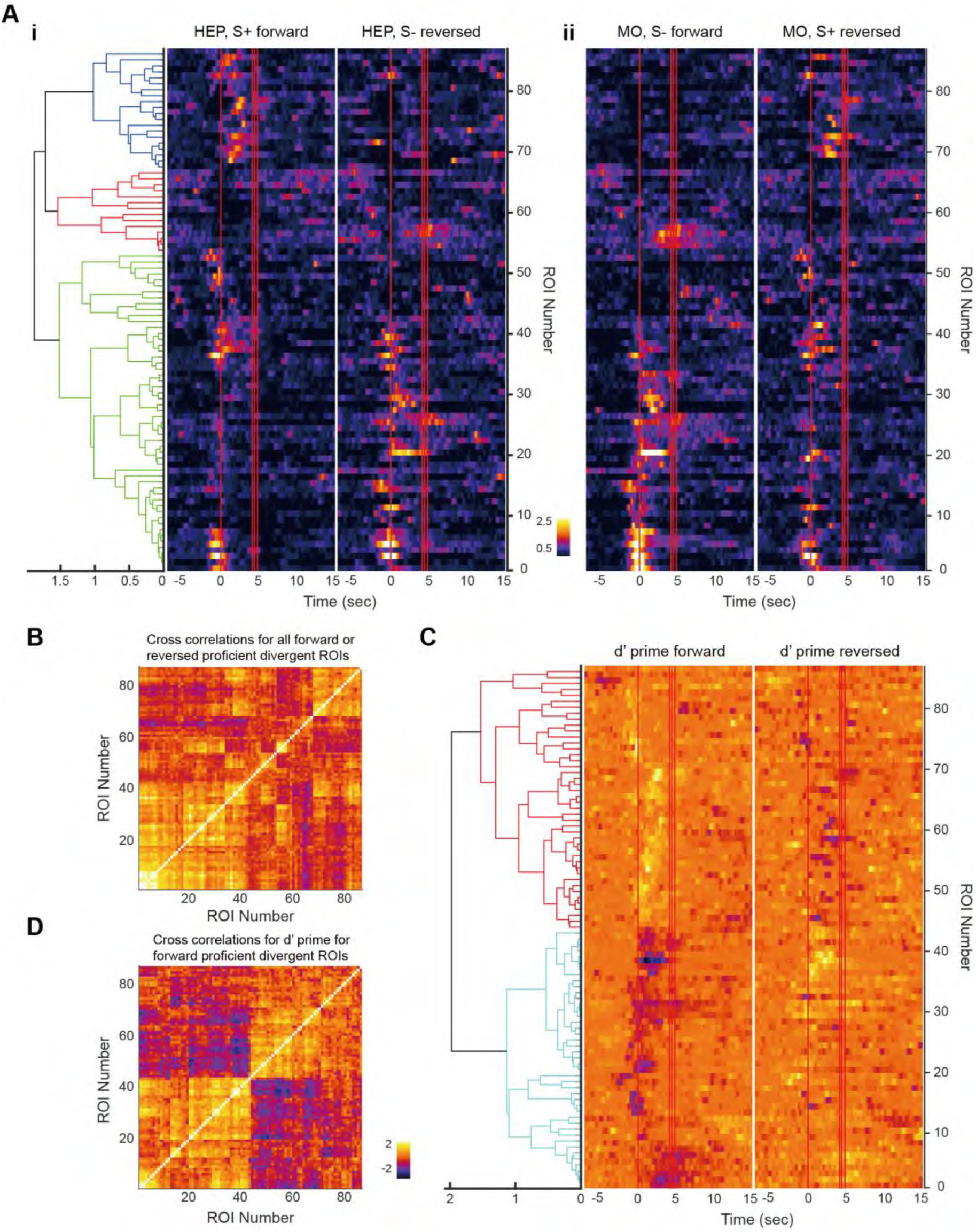
Supplemental figure for Figure 5D characterizing zΔF/F for reversal calculated for one mouse with matched ROIs between forward and reversed sessions. (A) zΔF/F time courses for single ROIs that were divergent between S+ and S- trials. (i) shows zΔF/F time courses for HEP in the forward and reversed proficient sessions (1 session) and (ii) shows zΔF/F time courses for MO in the forward and reversed proficient sessions (2 sessions). Time courses were sorted by estimating an agglomerative hierarchical cluster tree shown on the left that was calculated using the cross-correlation coefficients between all divergent zΔF/F time courses (panel B). The red vertical lines show (in order): trial start, odorant on, odorant off, reinforcement on and reinforcement off. (B) Cross-correlation coefficients computed between all per trial zΔF/F time courses shown in Figure S8A. The coefficients were sorted by the agglomerative hierarchical cluster tree shown in that figure. (C) zΔF/F d’ time courses for single ROIs that were divergent between S+ and S- trials. (i) forward (1 session) and (ii) reversed (2 sessions) (d’ was calculated for HEP vs. MO, see methods). Time courses were sorted by estimating an agglomerative hierarchical cluster tree shown on the left that was calculated using the cross-correlation coefficients between all d’ zΔF/F time courses (panel D). The red vertical lines show (in order): trial start, odorant on, odorant off, reinforcement on and reinforcement off. (D) Cross-correlation coefficients computed between all d’ zΔF/F time courses shown in Figure S8C. The coefficients were sorted by the agglomerative hierarchical cluster tree shown in that figure.

**Figure S9.**
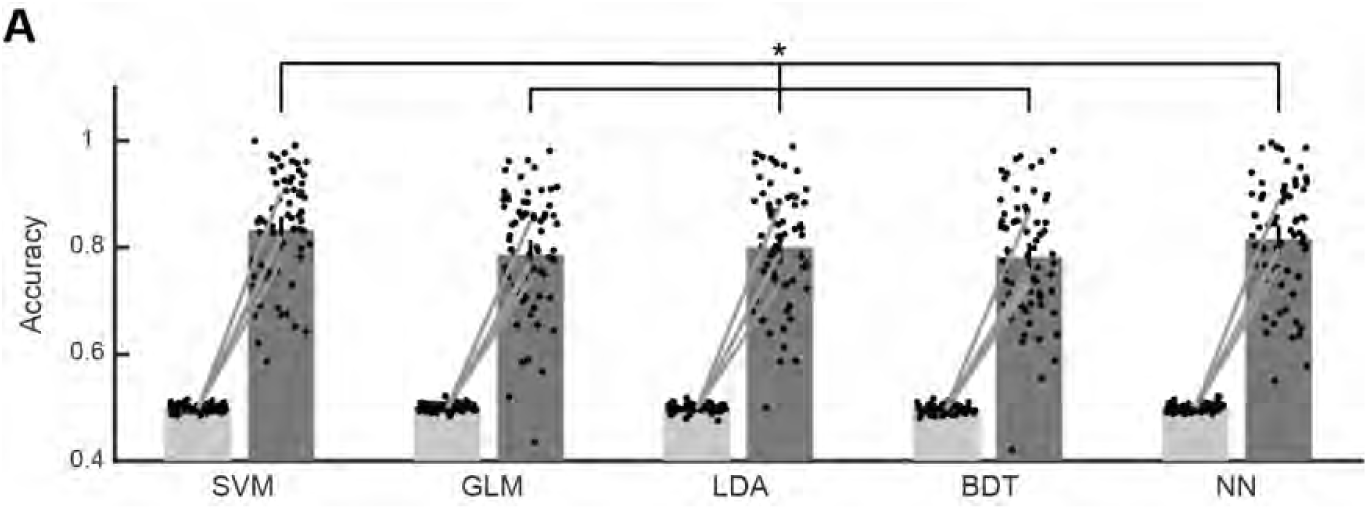
Stimulus decoding accuracy obtained by training in the wide odorant window with different decoding algorithms. The bar graph shows the mean stimulus decoding accuracy obtained in the 3.1 to 4.1 sec window for different decoding algorithms trained with data in the 0.5 to 5.5 sec window for all proficient sessions (66 sessions, 4 mice, related to Figure 6). The different algorithms are: GLM: generalized linear model, SVM: support vector machine, BDT: binary decision tree, NN: neural network and LDA: linear discriminant analysis. A GLM analysis yields a statistically significant difference between GLM vs. SVM and GLM vs. NN (n=684, 671 d.f., F-statistic 203, p-value <0.001, Table S11).

**Figure S10.**
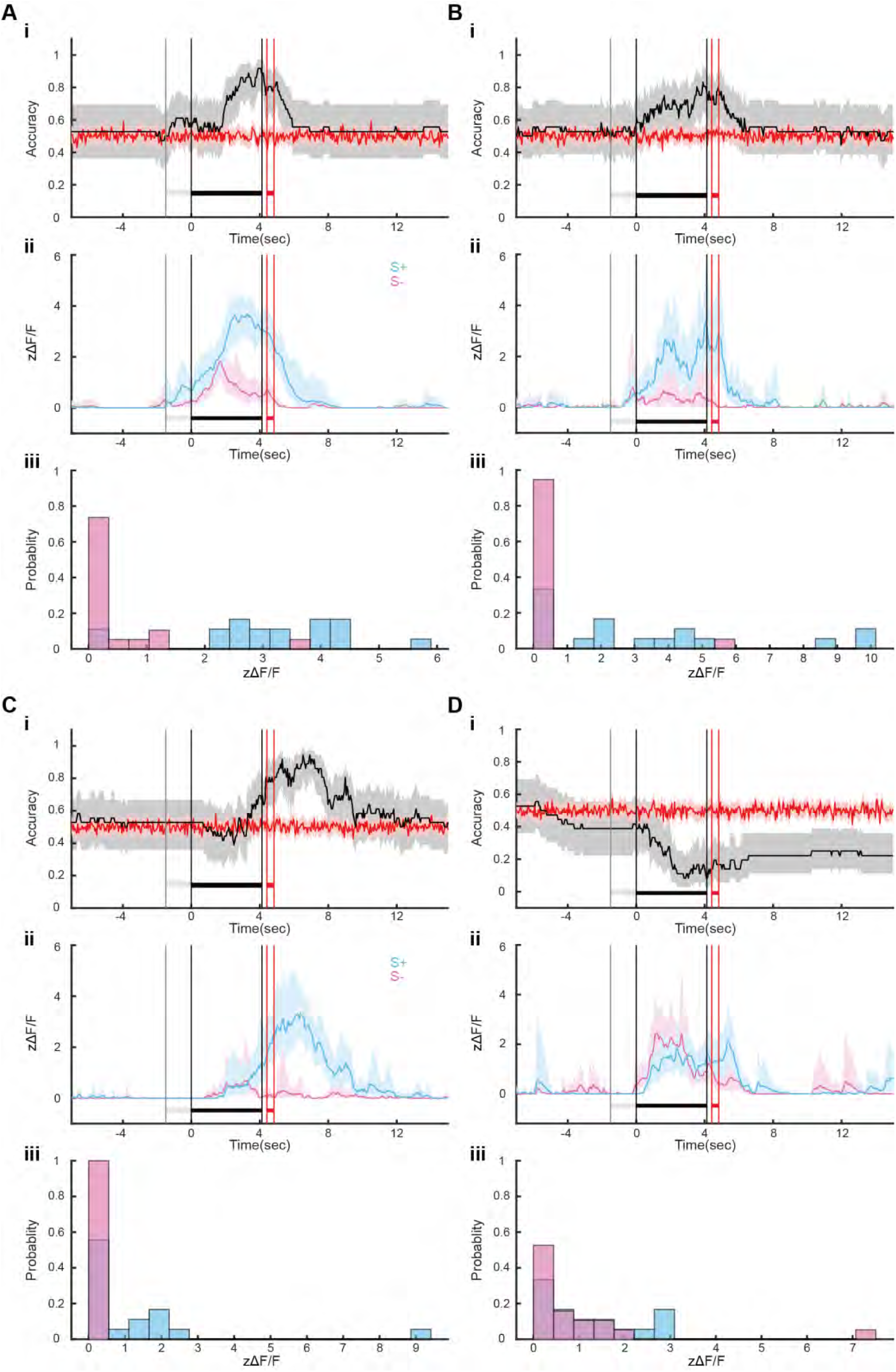
Examples of single ROI stimulus decoding. (A to D) show four examples of single ROI stimulus decoding. i. Decoding accuracy (black) vs. shuffled (red). ii. zΔF/F time courses for S+ (cyan) and S- (magenta). iii. Histogram of mean zΔF/F computed between 2 and 4.1 sec. This session had 62 ROIs with 18 S+ trials and 19 S- trials.

**Figure S11.**
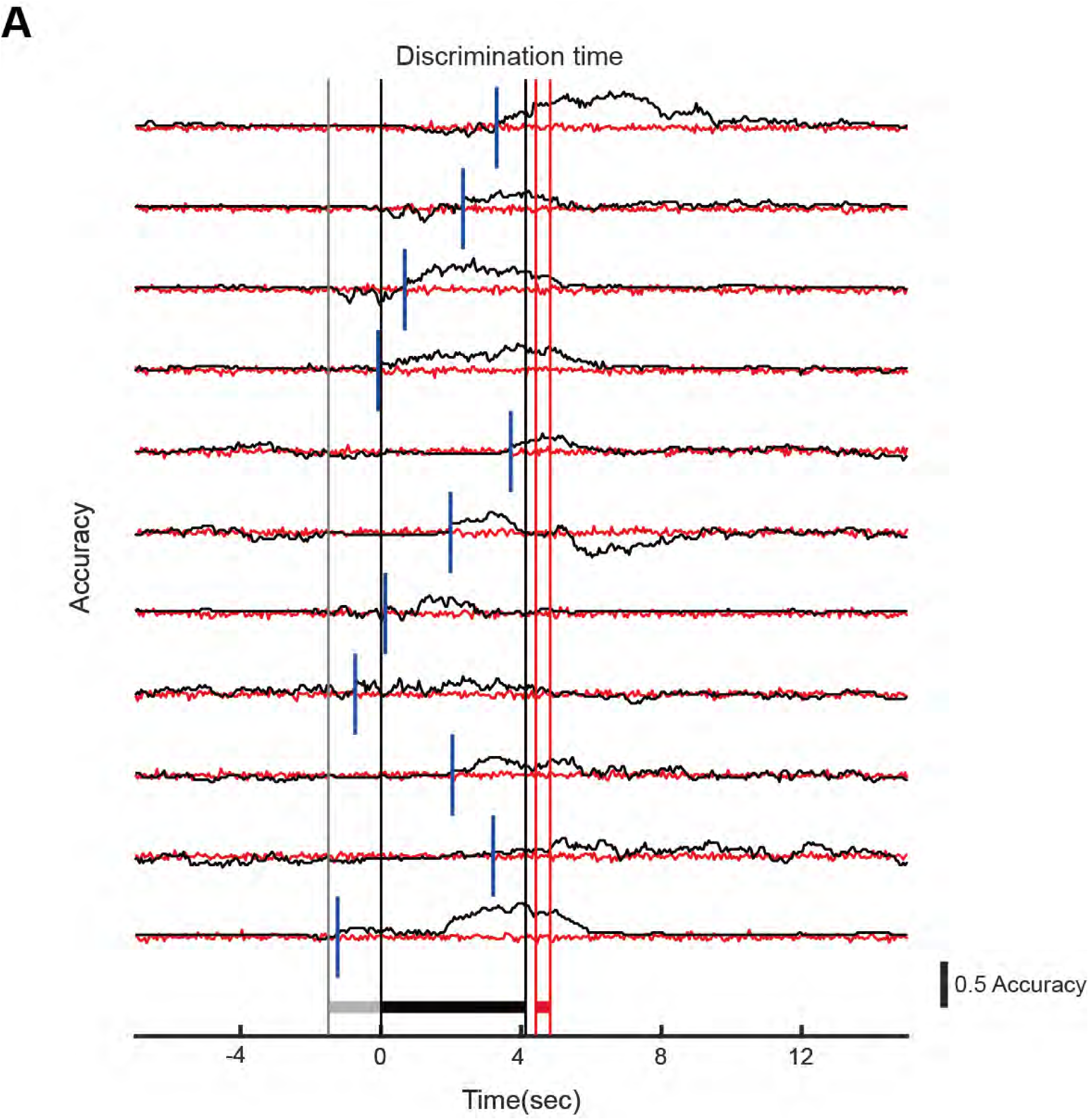
Examples of stimulus decoding accuracy time courses for one session. The figure shows stimulus decoding accuracy time courses for all ROIs that yielded accuracies > 0.65 for the proficient session of the examples shown in Figure S10.

**Figure S12.**
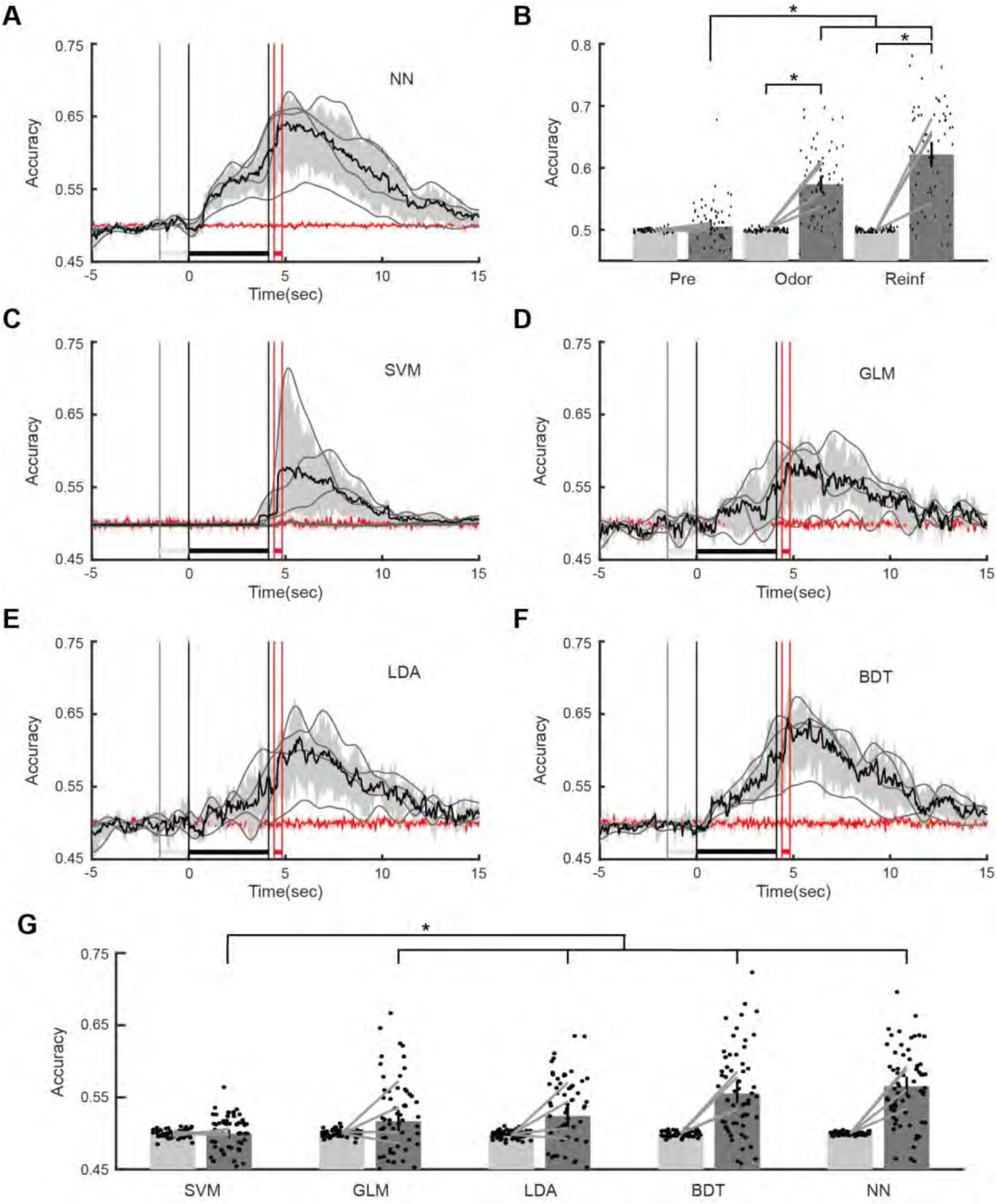
Lick decoding accuracy. **A.** Time course per trial for the accuracy for NN decoding of lick fraction from zΔF/F responses for all trials and all ROIs per session for all proficient sessions (4 mice, 66 proficient sessions). The bounded black line represents the mean accuracy bounded by the 95% CI. The grey lines are per mouse accuracies. The red line is the stimulus decoding accuracy calculated after shuffling the stimulus labels (S+ vs S-). **B.** Bar graph showing the mean accuracy for different accuracy periods (Pre -1 to 0, Odor 3.1 to 4.1 and Reinf 4.5 to 5.5). Light gray bars are the shuffled stimulus accuracies. Points are per session accuracies and bars are 95% CIs. **C-F.** Time course per trial for the accuracy for decoding of lick fraction from zΔF/F responses for all trials and all ROIs per session for all proficient sessions (4 mice, 66 proficient sessions). The algorithms used for the different panels are SVM (C), GLM (D), LDA (E) and BDT (F). The bounded black line represents the mean accuracy bounded by the 95% CI. The grey lines are per mouse accuracies. The red line is the stimulus decoding accuracy calculated after shuffling the stimulus labels (S+ vs S-). **G.** The bar graph shows the mean lick fraction decoding accuracy obtained in the 3.1 to 4.1 sec window for different decoding algorithms (66 sessions, 4 mice). The different algorithms are: GLM: generalized linear model, SVM: support vector machine, BDT: binary decision tree, NN: neural network and LDA: linear discriminant analysis. A GLM analysis yields a statistically significant difference between all algorithms and SVM (n=630, 617 d.f., F-statistic 27.5, p-value <0.001, Table S12). *p<0.05 for a pFDR-corrected t-test or ranksum tests, GLM statistics for B and G are in Table S12.

**Video 1.**
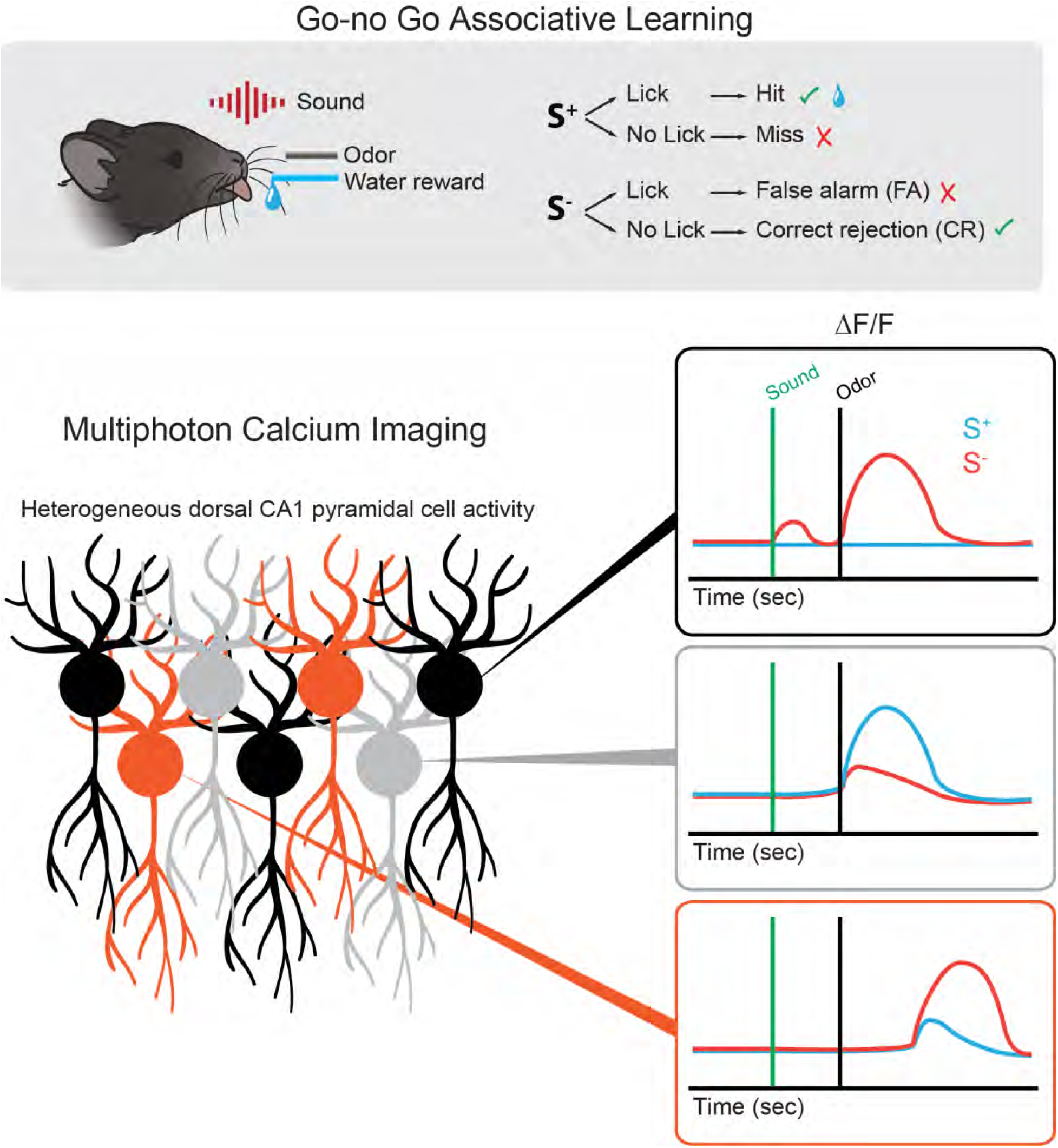
Movie of fluorescence changes for CA1 pyramidal cells shown in Figure 1C

## In Brief

Ma et al. show that stimulus divergence of pyramidal cell neural activity in dorsal CA1 takes place at discrete times during the mouse’s response in a go-no go associative learning odorant discrimination task. The sequential neural dynamics of these “decision- making time cells” during odorant discrimination in dorsal CA1 would provide a cognitive time map for decision making in associative learning.

## Highlights

- Spiking of pyramidal cells encodes for stimulus identity and lick behavior in dorsal CA1 for a mouse engaged in a go-no go odorant associative learning task
- The time of onset of stimulus divergence in neural activity of individual pyramidal cells in dorsal CA1 is time-tiled
- Learning increases the accuracy of decoding stimulus identity in this task
- The majority of pyramidal cells encode for odorant valence as opposed to odorant identity

