## Supplementary material for "Decision-Making Time Cells in Hippocampal Dorsal CA1": Table S2 (from another version of this preprint: version 2, 2023-10-11)

Table S2 glm statistics for Figures 2C and 2D

glm statistics for mean zdF/F bar graph in Figure 2C

mdl =

Generalized linear regression model:
    mean_dFF ~ p_correct+mice+time_window+sp_vs_sm+p_correct*time_window*sp_vs_sm
    Distribution = Normal

p_correct is learning (1) vs. proficient (3) stages
time_window is baseline (1, -4.5 to -3.5), pre (2, -1 to 0), odor (3, 3.1 to 4.1) and reinforcement (4, 4.5 5.5)
sp_vs_sm is S- (0) and S+ (1)
mice are the mouse numbers 1-4

Estimated Coefficients:

| | Estimate | SE | tStat | pValue |
|---|---|---|---|---|
| (Intercept) | 0.20075 | 0.0088004 | 22.812 | 3.053e-87 |
| p_correct_3 | 0.0017895 | 0.010356 | 0.1728 | 0.86286 |
| time_window_2 | 0.12052 | 0.011848 | 10.172 | 8.2579e-23 |
| time_window_3 | 0.048157 | 0.011848 | 4.0645 | 5.3388e-05 |
| time_window_4 | 0.027305 | 0.011848 | 2.3046 | 0.021471 |
| sp_vs_sm_1 | -0.0087942 | 0.011848 | -0.74226 | 0.45817 |
| mice_2 | -0.0033362 | 0.0044465 | -0.75029 | 0.45333 |
| mice_3 | -0.011421 | 0.0054235 | -2.1058 | 0.035568 |
| mice_4 | 0.018515 | 0.0047376 | 3.908 | 0.00010178 |
| p_correct_3:time_window_2 | -0.015015 | 0.014629 | -1.0264 | 0.30506 |
| p_correct_3:time_window_3 | -0.0084742 | 0.014629 | -0.57926 | 0.56259 |
| p_correct_3:time_window_4 | -0.0011829 | 0.014629 | -0.080859 | 0.93558 |
| p_correct_3:sp_vs_sm_1 | 0.014269 | 0.014629 | 0.97539 | 0.32969 |
| time_window_2:sp_vs_sm_1 | -0.0053483 | 0.016756 | -0.31919 | 0.74967 |
| time_window_3:sp_vs_sm_1 | 0.011913 | 0.016756 | 0.71101 | 0.47731 |
| time_window_4:sp_vs_sm_1 | 0.029637 | 0.016756 | 1.7688 | 0.077349 |
| p_correct_3:time_window_2:sp_vs_sm_1 | 0.016012 | 0.020689 | 0.77392 | 0.43923 |
| p_correct_3:time_window_3:sp_vs_sm_1 | 0.018498 | 0.020689 | 0.89411 | 0.37156 |
| p_correct_3:time_window_4:sp_vs_sm_1 | 0.00053314 | 0.020689 | 0.025769 | 0.97945 |

744 observations, 725 error degrees of freedom
Estimated Dispersion: 0.00225
F-statistic vs. constant model: 34.6, p-value = 3.79e-85

Ranksum or t-test p values for mean dFF

pFDR = 3.875000e-02

p value t-test for Pre S+ learning vs Baseline S+ proficient =  3.244708e-19
p value t-test for Pre S+ learning vs Baseline S- proficient =  4.532024e-19
p value ranksum for Baseline S+ proficient vs Pre S+ proficient =  7.142261e-18
p value ranksum for Baseline S+ proficient vs Pre S- proficient =  1.520838e-17
p value t-test for Pre S- learning vs Baseline S- proficient =  2.472835e-17
p value t-test for Pre S- learning vs Baseline S+ proficient =  3.636530e-17
p value t-test for Baseline S- proficient vs Pre S- proficient =  1.577037e-16
p value t-test for Pre S+ learning vs Reinf S- proficient =  5.905494e-16
p value ranksum for Pre S+ proficient vs Reinf S- proficient =  8.323538e-16
p value t-test for Baseline S- proficient vs Pre S+ proficient =  1.000287e-14
p value ranksum for Baseline S+ proficient vs Odor S+ proficient =  2.551906e-14
p value t-test for Pre S- learning vs Reinf S- proficient =  3.451704e-14
p value t-test for Pre S- proficient vs Reinf S- proficient =  4.956258e-14
p value ranksum for Baseline S+ learning vs Pre S- proficient =  1.035955e-13
p value ranksum for Baseline S+ learning vs Pre S+ proficient =  1.322561e-13
p value ranksum for Pre S+ proficient vs Odor S- proficient =  7.028861e-13
p value ranksum for Baseline S+ proficient vs Reinf S+ proficient =  8.473449e-13
p value ranksum for Baseline S- learning vs Pre S+ proficient =  2.424230e-12
p value ranksum for Baseline S+ learning vs Odor S+ proficient =  2.882264e-12

```
p value ranksum for Baseline S- learning vs Pre S- proficient =  3.053031e-12
p value t-test for Pre S- proficient vs Odor S- proficient =  3.191235e-12
p value t-test for Baseline S- proficient vs Odor S+ proficient =  3.903648e-12
p value t-test for Pre S+ learning vs Odor S- proficient =  2.511419e-11
p value ranksum for Baseline S+ learning vs Pre S- learning =  4.089275e-11
p value ranksum for Baseline S+ learning vs Reinf S+ proficient =  4.247975e-11
p value t-test for Baseline S- proficient vs Reinf S+ proficient =  6.208020e-11
p value t-test for Pre S- learning vs Odor S- proficient =  7.976828e-11
p value t-test for Pre S- proficient vs Reinf S+ proficient =  9.973710e-11
p value ranksum for Baseline S+ learning vs Pre S+ learning =  2.029819e-10
p value ranksum for Baseline S- learning vs Odor S+ proficient =  6.301050e-10
p value ranksum for Baseline S- learning vs Pre S+ learning =  8.746076e-10
p value t-test for Baseline S- proficient vs Odor S- proficient =  9.496892e-10
p value ranksum for Baseline S- learning vs Pre S- learning =  1.224217e-09
p value ranksum for Baseline S+ proficient vs Odor S- proficient =  1.520906e-09
p value t-test for Odor S+ learning vs Baseline S- proficient =  2.300095e-09
p value t-test for Odor S+ proficient vs Reinf S- proficient =  3.043137e-09
p value t-test for Odor S+ learning vs Baseline S+ proficient =  4.628484e-09
p value ranksum for Baseline S+ learning vs Odor S- proficient =  6.529109e-09
p value ranksum for Reinf S- learning vs Pre S+ proficient =  1.540078e-08
p value ranksum for Baseline S+ learning vs Odor S+ learning =  2.596052e-08
p value ranksum for Baseline S- learning vs Reinf S+ proficient =  3.099365e-08
p value t-test for Reinf S+ learning vs Baseline S- proficient =  3.516513e-08
p value t-test for Baseline S- proficient vs Reinf S- proficient =  7.869508e-08
p value t-test for Pre S- learning vs Reinf S- learning =  7.965428e-08
p value t-test for Reinf S+ learning vs Baseline S+ proficient =  9.236531e-08
p value ranksum for Baseline S+ learning vs Reinf S+ learning =  1.355021e-07
p value ranksum for Reinf S- learning vs Pre S- proficient =  1.786085e-07
p value ranksum for Pre S+ proficient vs Reinf S+ proficient =  1.967541e-07
p value ranksum for Baseline S+ learning vs Reinf S- proficient =  4.048395e-07
p value ranksum for Odor S- learning vs Pre S+ proficient =  9.739229e-07
p value ranksum for Reinf S+ proficient vs Reinf S- proficient =  1.008201e-06
p value t-test for Odor S+ proficient vs Odor S- proficient =  1.228321e-06
p value t-test for Reinf S+ learning vs Pre S+ proficient =  2.553190e-06
p value t-test for Pre S+ learning vs Reinf S- learning =  3.944320e-06
p value t-test for Reinf S+ learning vs Pre S- proficient =  4.091071e-06
p value t-test for Odor S+ learning vs Pre S+ proficient =  4.982811e-06
p value t-test for Pre S- learning vs Odor S- learning =  5.175747e-06
p value ranksum for Baseline S+ proficient vs Reinf S- proficient =  5.576530e-06
p value ranksum for Baseline S- learning vs Odor S+ learning =  7.045520e-06
p value ranksum for Odor S- learning vs Pre S- proficient =  8.832255e-06
p value t-test for Odor S+ learning vs Pre S- proficient =  8.852150e-06
p value ranksum for Baseline S+ learning vs Odor S- learning =  9.055054e-06
p value t-test for Pre S- learning vs Reinf S+ proficient =  9.751293e-06
p value t-test for Pre S- proficient vs Odor S+ proficient =  1.193370e-05
p value ranksum for Baseline S- learning vs Odor S- proficient =  1.330735e-05
p value ranksum for Odor S- learning vs Baseline S- proficient =  3.059629e-05
p value ranksum for Baseline S- learning vs Reinf S+ learning =  3.425965e-05
p value t-test for Pre S+ learning vs Reinf S+ learning =  3.578687e-05
p value ranksum for Odor S- learning vs Baseline S+ proficient =  3.781190e-05
p value t-test for Pre S+ learning vs Reinf S+ proficient =  6.359675e-05
p value ranksum for Reinf S- learning vs Odor S+ proficient =  7.794699e-05
p value t-test for Pre S+ learning vs Odor S+ learning =  8.279111e-05
p value t-test for Odor S+ learning vs Reinf S- proficient =  9.017046e-05
p value ranksum for Pre S+ proficient vs Odor S+ proficient =  1.153991e-04
p value t-test for Pre S- learning vs Reinf S+ learning =  2.024255e-04
p value t-test for Pre S+ learning vs Odor S- learning =  2.253608e-04
p value t-test for Pre S- learning vs Odor S+ learning =  3.795512e-04
p value ranksum for Baseline S- learning vs Odor S- learning =  4.448745e-04
p value t-test for Pre S- learning vs Odor S+ proficient =  7.482486e-04
p value t-test for Odor S- proficient vs Reinf S- proficient =  8.203481e-04
p value t-test for Reinf S+ learning vs Reinf S- proficient =  9.233557e-04
p value t-test for Odor S- proficient vs Reinf S+ proficient =  9.721085e-04
p value ranksum for Baseline S- learning vs Reinf S- proficient =  1.166142e-03
p value ranksum for Baseline S+ learning vs Reinf S- learning =  1.297798e-03
p value ranksum for Reinf S- learning vs Reinf S+ proficient =  1.925663e-03
p value t-test for Odor S+ proficient vs Reinf S+ proficient =  3.997192e-03
p value t-test for Pre S+ learning vs Odor S+ proficient =  6.265701e-03
p value t-test for Reinf S+ learning vs Odor S+ proficient =  1.083575e-02
p value ranksum for Odor S- learning vs Odor S+ proficient =  1.348648e-02
p value ranksum for Reinf S- learning vs Baseline S- proficient =  1.379451e-02
p value t-test for Odor S+ learning vs Odor S+ proficient =  2.122361e-02
```

p value ranksum for Baseline S- learning vs Reinf S- learning = 2.363928e-02
p value ranksum for Baseline S+ learning vs Baseline S+ proficient = 2.430321e-02

p values below are > pFDR

p value ranksum for Reinf S- learning vs Baseline S+ proficient = 4.620569e-02
p value ranksum for Reinf S- learning vs Odor S- proficient = 5.897562e-02
p value t-test for Odor S+ learning vs Odor S- proficient = 8.495736e-02
p value t-test for Odor S+ learning vs Reinf S- learning = 8.617064e-02
p value ranksum for Odor S- learning vs Reinf S- learning = 1.085212e-01
p value ranksum for Odor S- learning vs Reinf S- proficient = 1.157253e-01
p value t-test for Reinf S+ learning vs Reinf S- learning = 1.166726e-01
p value ranksum for Odor S- learning vs Reinf S+ proficient = 1.195186e-01
p value t-test for Reinf S+ learning vs Reinf S+ proficient = 1.907769e-01
p value t-test for Reinf S+ learning vs Odor S- proficient = 2.139943e-01
p value ranksum for Baseline S+ learning vs Baseline S- proficient = 2.204924e-01
p value ranksum for Pre S+ proficient vs Pre S- proficient = 2.450964e-01
p value t-test for Pre S+ learning vs Pre S- learning = 2.452675e-01
p value t-test for Pre S- learning vs Pre S- proficient = 2.658255e-01
p value t-test for Odor S+ learning vs Reinf S+ proficient = 3.108015e-01
p value t-test for Pre S+ learning vs Pre S+ proficient = 3.472551e-01
p value ranksum for Baseline S- learning vs Baseline S+ proficient = 3.629192e-01
p value ranksum for Baseline S+ learning vs Baseline S- learning = 4.724486e-01
p value ranksum for Reinf S- learning vs Reinf S- proficient = 4.791718e-01
p value ranksum for Baseline S+ proficient vs Baseline S- proficient = 5.321712e-01
p value t-test for Odor S+ learning vs Reinf S+ learning = 5.451055e-01
p value ranksum for Odor S- learning vs Reinf S+ learning = 6.239980e-01
p value t-test for Odor S+ learning vs Odor S- learning = 8.065586e-01
p value ranksum for Baseline S- learning vs Baseline S- proficient = 8.874536e-01
p value t-test for Pre S+ learning vs Pre S- proficient = 8.910594e-01
p value t-test for Pre S- learning vs Pre S+ proficient = 9.866997e-01
p value ranksum for Odor S- learning vs Odor S- proficient = 9.903212e-01

glm statistics for delta euclidean distance bar graph in Figure 2E

mdl =

Generalized linear regression model:
euclidean_distance ~
p_correct+time_window+mice+between_vs_within+p_correct*time_window*between_vs_within
Distribution = Normal

p_correct is learning (1) vs. proficient (3) stages
time_window is baseline (1, -4.5 to -3.5), pre (2, -1 to 0), odor (3, 3.1 to 4.1) and reinforcement (4, 4.5 5.5)
between_vs_within is between S+ and S- (1) vs. wothin S+ or S- (0)
mice are the mouse numbers 1-4

Estimated Coefficients:

| | Estimate | SE | tStat | pValue |
|---|---|---|---|---|
| (Intercept) | -0.20384 | 0.27381 | -0.74444 | 0.45685 |
| p_correct_3 | 0.18264 | 0.32222 | 0.56683 | 0.57101 |
| time_window_2 | 3.5086 | 0.36863 | 9.5178 | 2.5606e-20 |
| time_window_3 | 1.9318 | 0.36863 | 5.2403 | 2.1038e-07 |
| time_window_4 | 1.5507 | 0.36863 | 4.2066 | 2.917e-05 |
| between_vs_within_1 | 0.0043261 | 0.36863 | 0.011735 | 0.99064 |
| mice_2 | 0.43566 | 0.13835 | 3.149 | 0.0017055 |
| mice_3 | -0.86197 | 0.16875 | -5.1081 | 4.1641e-07 |
| mice_4 | 0.19865 | 0.1474 | 1.3477 | 0.17818 |
| p_correct_3:time_window_2 | 0.12604 | 0.45517 | 0.2769 | 0.78193 |
| p_correct_3:time_window_3 | 0.55967 | 0.45517 | 1.2296 | 0.21925 |
| p_correct_3:time_window_4 | 0.17404 | 0.45517 | 0.38235 | 0.70231 |
| p_correct_3:between_vs_within_1 | -0.0035289 | 0.45517 | -0.007753 | 0.99382 |
| time_window_2:between_vs_within_1 | -0.050111 | 0.52133 | -0.096122 | 0.92345 |
| time_window_3:between_vs_within_1 | -0.39513 | 0.52133 | -0.75793 | 0.44874 |
| time_window_4:between_vs_within_1 | -0.49467 | 0.52133 | -0.94888 | 0.343 |
| p_correct_3:time_window_2:between_vs_within_1 | 0.069145 | 0.6437 | 0.10742 | 0.91449 |

```
    p_correct_3:time_window_3:between_vs_within_1      -0.40267     0.6437     -0.62555       0.53181
    p_correct_3:time_window_4:between_vs_within_1      -0.32371     0.6437     -0.50288        0.6152


744 observations, 725 error degrees of freedom
Estimated Dispersion: 2.17
F-statistic vs. constant model: 36.6, p-value = 3.49e-89



Ranksum or t-test p values for euclidean distance


pFDR = 3.791667e-02

p value t-test for Pre within learning vs Baseline within proficient =  2.075095e-25
p value t-test for Pre within learning vs Baseline between proficient =  2.669672e-25
p value t-test for Pre between learning vs Baseline within proficient =  2.874446e-24
p value t-test for Pre between learning vs Baseline between proficient =  3.576862e-24
p value t-test for Baseline between proficient vs Pre within proficient =  1.473805e-19
p value t-test for Baseline within proficient vs Pre within proficient =  1.545371e-19
p value t-test for Baseline between proficient vs Pre between proficient =  2.098321e-19
p value t-test for Baseline within proficient vs Pre between proficient =  2.217221e-19
p value t-test for Baseline between learning vs Pre within proficient =  4.529538e-17
p value t-test for Baseline within learning vs Pre within proficient =  4.662274e-17
p value t-test for Baseline between learning vs Pre between proficient =  6.362717e-17
p value t-test for Baseline within learning vs Pre between proficient =  6.550536e-17
p value t-test for Pre within proficient vs Reinf within proficient =  1.493866e-16
p value t-test for Pre between proficient vs Reinf within proficient =  1.883191e-16
p value t-test for Reinf within proficient vs Reinf between proficient =  5.050262e-16
p value t-test for Baseline within proficient vs Odor between proficient =  8.971982e-16
p value t-test for Baseline between proficient vs Odor between proficient =  9.532307e-16
p value t-test for Odor within proficient vs Odor between proficient =  1.825790e-15
p value t-test for Baseline between learning vs Odor between proficient =  8.921742e-15
p value t-test for Baseline within learning vs Odor between proficient =  9.210953e-15
p value t-test for Baseline between learning vs Pre within learning =  3.998822e-14
p value t-test for Baseline within learning vs Pre within learning =  5.257405e-14
p value t-test for Baseline between learning vs Pre between learning =  9.006212e-14
p value t-test for Baseline within learning vs Pre between learning =  1.210481e-13
p value t-test for Pre within proficient vs Odor within proficient =  1.166483e-12
p value t-test for Pre between proficient vs Odor within proficient =  2.310089e-12
p value t-test for Pre within learning vs Reinf within proficient =  3.330069e-12
p value t-test for Odor between proficient vs Reinf within proficient =  3.876411e-12
p value t-test for Baseline within proficient vs Reinf between proficient =  5.072155e-12
p value t-test for Pre between learning vs Reinf within proficient =  5.669923e-12
p value t-test for Baseline between proficient vs Reinf between proficient =  7.056166e-12
p value t-test for Pre within proficient vs Reinf between proficient =  9.601531e-12
p value t-test for Pre between proficient vs Reinf between proficient =  1.302084e-11
p value t-test for Baseline within proficient vs Odor within proficient =  1.391287e-10
p value t-test for Baseline between proficient vs Odor within proficient =  1.514706e-10
p value t-test for Baseline between learning vs Reinf between proficient =  1.923330e-10
p value t-test for Baseline within learning vs Reinf between proficient =  2.002927e-10
p value t-test for Baseline between learning vs Odor within proficient =  3.402093e-09
p value t-test for Baseline within learning vs Odor within proficient =  3.547501e-09
p value ranksum for Reinf within learning vs Pre within proficient =  2.527825e-07
p value t-test for Odor within learning vs Baseline within proficient =  2.899293e-07
p value t-test for Odor within learning vs Baseline between proficient =  3.101248e-07
p value ranksum for Reinf within learning vs Pre between proficient =  3.271333e-07
p value t-test for Odor within learning vs Pre within proficient =  5.899519e-07
p value t-test for Odor within learning vs Pre between proficient =  7.401686e-07
p value ranksum for Odor between learning vs Baseline between proficient =  1.147543e-06
p value ranksum for Odor between learning vs Baseline within proficient =  1.195396e-06
p value t-test for Pre within learning vs Reinf within learning =  1.239267e-06
p value t-test for Pre within learning vs Reinf between proficient =  1.332079e-06
p value t-test for Pre between learning vs Reinf between proficient =  1.418524e-06
p value t-test for Pre within proficient vs Odor between proficient =  1.498358e-06
p value t-test for Pre between learning vs Reinf within learning =  2.040676e-06
p value t-test for Reinf between learning vs Pre within proficient =  2.241282e-06
p value t-test for Pre between proficient vs Odor between proficient =  2.527087e-06
p value t-test for Pre within learning vs Odor within proficient =  2.730668e-06
p value t-test for Reinf between learning vs Pre between proficient =  2.751478e-06
p value t-test for Pre between learning vs Odor within proficient =  2.753772e-06
```

```
p value t-test for Reinf between learning vs Baseline within proficient =  5.279049e-06
p value t-test for Reinf between learning vs Baseline between proficient =  5.488456e-06
p value t-test for Odor within proficient vs Reinf within proficient =  5.769590e-06
p value t-test for Baseline within proficient vs Reinf within proficient =  7.116701e-06
p value t-test for Pre within learning vs Odor within learning =  7.455720e-06
p value t-test for Pre between learning vs Odor within learning =  8.588666e-06
p value t-test for Baseline between proficient vs Reinf within proficient =  8.737603e-06
p value t-test for Odor between proficient vs Reinf between proficient =  1.690230e-05
p value ranksum for Odor between learning vs Pre between proficient =  2.383124e-05
p value ranksum for Odor between learning vs Pre within proficient =  2.560288e-05
p value t-test for Baseline between learning vs Odor between learning =  5.811247e-05
p value t-test for Baseline within learning vs Odor between learning =  6.737320e-05
p value t-test for Baseline between learning vs Reinf within proficient =  6.834456e-05
p value t-test for Baseline between learning vs Odor within learning =  6.942655e-05
p value t-test for Baseline within learning vs Reinf within proficient =  7.151884e-05
p value t-test for Pre within learning vs Reinf between learning =  7.460743e-05
p value t-test for Baseline within learning vs Odor within learning =  8.124653e-05
p value t-test for Pre between learning vs Reinf between learning =  8.853852e-05
p value ranksum for Reinf within learning vs Odor between proficient =  1.089454e-04
p value t-test for Baseline between learning vs Reinf between learning =  5.075998e-04
p value t-test for Baseline within learning vs Reinf between learning =  5.464373e-04
p value t-test for Pre between learning vs Odor between learning =  7.055563e-04
p value t-test for Pre within learning vs Odor between learning =  8.729948e-04
p value t-test for Odor within learning vs Odor between proficient =  2.508565e-03
p value t-test for Reinf between learning vs Odor between proficient =  5.535229e-03
p value t-test for Odor within learning vs Odor between learning =  6.755985e-03
p value t-test for Baseline between learning vs Reinf within learning =  8.278292e-03
p value t-test for Baseline within learning vs Reinf within learning =  8.704675e-03
p value t-test for Pre between learning vs Odor between proficient =  1.075800e-02
p value t-test for Pre within learning vs Odor between proficient =  1.284909e-02
p value ranksum for Reinf within learning vs Reinf between proficient =  1.880628e-02
p value ranksum for Odor between learning vs Odor between proficient =  2.587785e-02
p value ranksum for Reinf within learning vs Odor within proficient =  3.444544e-02
p value t-test for Odor within learning vs Reinf within learning =  3.557688e-02

p values below are > pFDR

p value ranksum for Reinf within learning vs Baseline within proficient =  4.892466e-02
p value ranksum for Reinf within learning vs Baseline between proficient =  4.986032e-02
p value ranksum for Odor between learning vs Reinf within proficient =  6.230580e-02
p value ranksum for Odor between learning vs Reinf within learning =  1.208643e-01
p value t-test for Odor within learning vs Reinf within proficient =  1.215296e-01
p value t-test for Reinf between learning vs Reinf within proficient =  1.502738e-01
p value t-test for Pre within learning vs Pre between learning =  2.747752e-01
p value ranksum for Reinf within learning vs Reinf between learning =  2.796488e-01
p value t-test for Odor within learning vs Reinf between proficient =  3.392288e-01
p value ranksum for Odor between learning vs Reinf between learning =  3.646608e-01
p value t-test for Pre within proficient vs Pre between proficient =  3.825032e-01
p value t-test for Baseline between learning vs Baseline within proficient =  3.884727e-01
p value t-test for Baseline between learning vs Baseline between proficient =  3.957086e-01
p value t-test for Reinf between learning vs Reinf between proficient =  3.968655e-01
p value t-test for Baseline within learning vs Baseline within proficient =  4.025409e-01
p value t-test for Odor within learning vs Odor within proficient =  4.096653e-01
p value t-test for Baseline within learning vs Baseline between proficient =  4.098388e-01
p value t-test for Pre within learning vs Pre within proficient =  4.242702e-01
p value t-test for Pre within learning vs Pre between proficient =  4.537002e-01
p value t-test for Reinf between learning vs Odor within proficient =  4.630607e-01
p value t-test for Pre between learning vs Pre within proficient =  5.004703e-01
p value t-test for Pre between learning vs Pre between proficient =  5.319733e-01
p value ranksum for Reinf within learning vs Reinf within proficient =  6.535450e-01
p value t-test for Baseline within learning vs Baseline between learning =  7.095733e-01
p value ranksum for Odor between learning vs Reinf between proficient =  7.555274e-01
p value ranksum for Odor between learning vs Odor within proficient =  7.678526e-01
p value t-test for Odor within proficient vs Reinf between proficient =  8.564304e-01
p value t-test for Baseline within proficient vs Baseline between proficient =  9.467667e-01
p value t-test for Odor within learning vs Reinf between learning =  9.608590e-01
```
