## Supplementary material for "Decision-Making Time Cells in Hippocampal Dorsal CA1": Table S3 (from another version of this preprint: version 2, 2023-10-11)

Table S3 glm statistics for dimensionality bar graph in Figure 2G

mdl =

Generalized linear regression model:
    dimensionality ~ 1 + mice + percent_correct + window + percent_correct*window
    Distribution = Normal

percent_correct is learning (1) vs. proficient (4)
window is pre (1, -1 to 0 sec) vs. odor (2, 3.1 to 4.1 sec)

Estimated Coefficients:

| | Estimate | SE | tStat | pValue |
|---|---|---|---|---|
| (Intercept) | 3.3649 | 0.62743 | 5.3631 | 1.0923e-06 |
| percent_correct_4 | 1.8838 | 0.6523 | 2.8879 | 0.0052181 |
| window_2 | 0.034444 | 0.78677 | 0.04378 | 0.96521 |
| mice_2 | 1.1964 | 0.56257 | 2.1267 | 0.037129 |
| mice_3 | -0.17506 | 0.484 | -0.36169 | 0.71872 |
| mice_4 | 2.7495 | 0.58288 | 4.717 | 1.258e-05 |
| percent_correct_4:window_2 | 0.46223 | 0.88868 | 0.52012 | 0.60469 |

74 observations, 67 error degrees of freedom
Estimated Dispersion: 2.48
F-statistic vs. constant model: 11.5, p-value = 8.39e-09

Ranksum or t-test p values for decoding lick fraction

pFDR = 4.166667e-02

p value t-test for Odor learning vs Odor proficient =  4.738985e-04
p value t-test for Pre learning vs Odor proficient =  4.774453e-04
p value t-test for Pre learning vs Pre proficient =  1.037346e-02
p value t-test for Odor learning vs Pre proficient =  1.077647e-02
p value t-test for Pre proficient vs Odor proficient =  3.423626e-02

p values below are > pFDR

p value t-test for Pre learning vs Odor learning =  9.040236e-01
