## Supplementary material for "Decision-Making Time Cells in Hippocampal Dorsal CA1": Table S4 (from another version of this preprint: version 2, 2023-10-11)

Table S4 glm statistics for Figures 3B and 3D

glm for bar graph of sum of dPCA stimulus components Figure 3B

mdl =

Generalized linear regression model:
dPCA_component ~ p_correct+time_window+sp_vs_sm+p_correct*time_window*sp_vs_sm
Distribution = Normal

p_correct is learning (1) vs. proficient (3) stages
time_window is baseline (1, -4.5 to -3.5), pre (2, -1 to 0), odor (3, 3.1 to 4.1) and reinforcement (4, 4.5 5.5)
sp_vs_sm is S- (0) and S+ (1)

Estimated Coefficients:

| | Estimate | SE | tStat | pValue |
|---|---|---|---|---|
| (Intercept) | 0.13442 | 2.0283 | 0.066273 | 0.94757 |
| p_correct_3 | -0.2706 | 2.8684 | -0.094335 | 0.92543 |
| time_window_2 | 3.0676 | 2.8684 | 1.0694 | 0.29288 |
| time_window_3 | 1.1035 | 2.8684 | 0.38471 | 0.703 |
| time_window_4 | 1.0895 | 2.8684 | 0.37981 | 0.70659 |
| sp_vs_sm_1 | -0.1902 | 2.8684 | -0.066307 | 0.94755 |
| p_correct_3:time_window_2 | -2.6992 | 4.0566 | -0.66538 | 0.51058 |
| p_correct_3:time_window_3 | -13.112 | 4.0566 | -3.2322 | 0.0028453 |
| p_correct_3:time_window_4 | -13.945 | 4.0566 | -3.4377 | 0.0016473 |
| p_correct_3:sp_vs_sm_1 | 0.45505 | 4.0566 | 0.11218 | 0.91138 |
| time_window_2:sp_vs_sm_1 | -4.8824 | 4.0566 | -1.2036 | 0.23758 |
| time_window_3:sp_vs_sm_1 | -2.1255 | 4.0566 | -0.52395 | 0.60392 |
| time_window_4:sp_vs_sm_1 | -2.1512 | 4.0566 | -0.53031 | 0.59956 |
| p_correct_3:time_window_2:sp_vs_sm_1 | 4.1492 | 5.7369 | 0.72326 | 0.47478 |
| p_correct_3:time_window_3:sp_vs_sm_1 | 26.172 | 5.7369 | 4.562 | 7.0625e-05 |
| p_correct_3:time_window_4:sp_vs_sm_1 | 27.896 | 5.7369 | 4.8626 | 2.9612e-05 |

48 observations, 32 error degrees of freedom
Estimated Dispersion: 12.3
F-statistic vs. constant model: 10.6, p-value = 1.77e-08

Ranksum for dPCA stimulus component

pFDR = 4.166667e-04

p values below are > pFDR

p value t-test for Pre S- learning vs Reinf S- proficient = 8.277020e-03
p value t-test for Odor S- learning vs Reinf S- proficient = 2.470342e-02
p value t-test for Pre S+ learning vs Reinf S+ proficient = 2.796285e-02
p value t-test for Pre S- learning vs Odor S- proficient = 2.990469e-02
p value t-test for Odor S+ learning vs Reinf S- learning = 3.278732e-02
p value t-test for Odor S+ learning vs Reinf S+ proficient = 3.485994e-02
p value t-test for Baseline S- learning vs Pre S+ learning = 3.658351e-02
p value t-test for Pre S+ proficient vs Reinf S+ proficient = 4.223515e-02
p value t-test for Pre S- proficient vs Reinf S- proficient = 4.226303e-02
p value t-test for Baseline S+ learning vs Reinf S- proficient = 4.686538e-02
p value t-test for Baseline S- learning vs Reinf S+ proficient = 4.767606e-02
p value t-test for Baseline S+ proficient vs Reinf S- proficient = 4.796368e-02
p value t-test for Baseline S- proficient vs Reinf S+ proficient = 4.803626e-02
p value t-test for Reinf S+ proficient vs Reinf S- proficient = 5.135212e-02
p value t-test for Reinf S- learning vs Reinf S- proficient = 5.201095e-02
p value t-test for Reinf S+ learning vs Reinf S+ proficient = 5.439491e-02
p value t-test for Baseline S- proficient vs Reinf S- proficient = 5.490043e-02
p value t-test for Baseline S+ proficient vs Reinf S+ proficient = 5.496036e-02
p value t-test for Baseline S- learning vs Reinf S- proficient = 5.598358e-02
p value t-test for Baseline S+ learning vs Reinf S+ proficient = 5.656941e-02

```
p value t-test for Reinf S+ learning vs Reinf S- proficient =  5.996977e-02
p value t-test for Pre S+ learning vs Baseline S- proficient =  6.283282e-02
p value t-test for Reinf S- learning vs Reinf S+ proficient =  6.284600e-02
p value t-test for Odor S- learning vs Odor S- proficient =  6.422479e-02
p value t-test for Pre S- proficient vs Reinf S+ proficient =  6.562389e-02
p value t-test for Pre S+ proficient vs Reinf S- proficient =  6.573054e-02
p value t-test for Pre S+ learning vs Odor S+ proficient =  6.748894e-02
p value t-test for Odor S- proficient vs Reinf S+ proficient =  7.730704e-02
p value t-test for Odor S+ proficient vs Reinf S- proficient =  7.734069e-02
p value t-test for Odor S+ learning vs Odor S+ proficient =  7.922417e-02
p value t-test for Odor S+ learning vs Reinf S- proficient =  8.164329e-02
p value t-test for Pre S+ learning vs Pre S+ proficient =  8.534612e-02
p value t-test for Pre S+ learning vs Reinf S- proficient =  9.060815e-02
p value t-test for Pre S+ learning vs Baseline S+ proficient =  9.255825e-02
p value t-test for Pre S+ proficient vs Odor S+ proficient =  9.388076e-02
p value t-test for Pre S- proficient vs Odor S- proficient =  9.395870e-02
p value t-test for Odor S- learning vs Reinf S+ proficient =  9.522872e-02
p value t-test for Reinf S- learning vs Odor S- proficient =  1.019079e-01
p value t-test for Baseline S+ learning vs Odor S- proficient =  1.025508e-01
p value t-test for Baseline S+ proficient vs Odor S- proficient =  1.032578e-01
p value t-test for Baseline S- proficient vs Odor S+ proficient =  1.032778e-01
p value t-test for Baseline S- learning vs Odor S+ proficient =  1.039723e-01
p value t-test for Reinf S+ learning vs Odor S+ proficient =  1.055159e-01
p value t-test for Odor S+ proficient vs Odor S- proficient =  1.085229e-01
p value t-test for Baseline S- learning vs Odor S- proficient =  1.138950e-01
p value t-test for Baseline S+ proficient vs Odor S+ proficient =  1.140136e-01
p value t-test for Baseline S- proficient vs Odor S- proficient =  1.140254e-01
p value t-test for Odor S- learning vs Reinf S+ learning =  1.146380e-01
p value t-test for Baseline S+ learning vs Odor S+ proficient =  1.150765e-01
p value t-test for Reinf S+ learning vs Odor S- proficient =  1.239897e-01
p value t-test for Pre S- proficient vs Odor S+ proficient =  1.278811e-01
p value t-test for Pre S+ proficient vs Odor S- proficient =  1.281051e-01
p value t-test for Reinf S- learning vs Odor S+ proficient =  1.284734e-01
p value t-test for Pre S+ learning vs Reinf S- learning =  1.323091e-01
p value t-test for Baseline S+ learning vs Pre S+ learning =  1.524022e-01
p value t-test for Odor S+ learning vs Odor S- proficient =  1.528723e-01
p value t-test for Pre S+ learning vs Odor S- learning =  1.544816e-01
p value t-test for Pre S+ learning vs Pre S- learning =  1.674651e-01
p value t-test for Pre S- learning vs Pre S- proficient =  1.682193e-01
p value t-test for Pre S+ learning vs Odor S- proficient =  1.708338e-01
p value t-test for Odor S- learning vs Odor S+ proficient =  1.711887e-01
p value t-test for Baseline S+ learning vs Pre S- learning =  1.790281e-01
p value t-test for Pre S- learning vs Reinf S+ proficient =  1.794895e-01
p value t-test for Pre S- learning vs Odor S+ learning =  1.925334e-01
p value t-test for Pre S- learning vs Reinf S+ learning =  1.975586e-01
p value t-test for Pre S+ learning vs Pre S- proficient =  2.073197e-01
p value t-test for Pre S- learning vs Baseline S+ proficient =  2.180518e-01
p value t-test for Pre S- learning vs Baseline S- proficient =  2.274715e-01
p value t-test for Odor S+ learning vs Pre S- proficient =  2.520909e-01
p value t-test for Reinf S- learning vs Pre S- proficient =  2.599109e-01
p value t-test for Baseline S- learning vs Reinf S+ learning =  2.742341e-01
p value t-test for Baseline S+ learning vs Reinf S- learning =  2.752335e-01
p value t-test for Pre S- learning vs Odor S+ proficient =  2.810825e-01
p value t-test for Baseline S- learning vs Pre S- learning =  2.830411e-01
p value t-test for Pre S- learning vs Pre S+ proficient =  2.882631e-01
p value t-test for Odor S- learning vs Baseline S- proficient =  2.935075e-01
p value t-test for Pre S- learning vs Odor S- learning =  2.955543e-01
p value t-test for Baseline S+ learning vs Odor S- learning =  3.055629e-01
p value t-test for Odor S+ learning vs Baseline S+ proficient =  3.117889e-01
p value t-test for Odor S+ learning vs Odor S- learning =  3.248674e-01
p value t-test for Reinf S+ learning vs Pre S+ proficient =  3.273390e-01
p value t-test for Odor S- learning vs Pre S+ proficient =  3.329405e-01
p value t-test for Odor S- learning vs Baseline S+ proficient =  3.462261e-01
p value t-test for Baseline S+ learning vs Odor S+ learning =  3.497810e-01
p value t-test for Reinf S- learning vs Baseline S- proficient =  3.515113e-01
p value t-test for Baseline S+ proficient vs Baseline S- proficient =  3.677087e-01
p value t-test for Baseline S- learning vs Baseline S- proficient =  3.684866e-01
p value t-test for Reinf S- learning vs Baseline S+ proficient =  3.840396e-01
p value t-test for Reinf S+ learning vs Reinf S- learning =  3.857055e-01
p value t-test for Baseline S- learning vs Odor S- learning =  3.859438e-01
p value t-test for Reinf S+ learning vs Baseline S+ proficient =  3.885780e-01
p value t-test for Baseline S- learning vs Odor S+ learning =  3.943657e-01
```

p value t-test for Baseline S- learning vs Pre S+ proficient =  4.033984e-01
p value t-test for Pre S- learning vs Reinf S- learning =  4.100197e-01
p value t-test for Baseline S+ learning vs Baseline S+ proficient =  4.251405e-01
p value t-test for Odor S+ learning vs Baseline S- proficient =  4.271653e-01
p value t-test for Reinf S+ learning vs Baseline S- proficient =  4.341727e-01
p value t-test for Reinf S- learning vs Pre S+ proficient =  4.741148e-01
p value t-test for Odor S- learning vs Pre S- proficient =  4.794550e-01
p value t-test for Pre S+ learning vs Odor S+ learning =  4.839357e-01
p value t-test for Baseline S+ learning vs Reinf S+ learning =  4.993440e-01
p value t-test for Baseline S- learning vs Reinf S- learning =  5.029876e-01
p value t-test for Reinf S+ learning vs Pre S- proficient =  5.038457e-01
p value t-test for Baseline S+ learning vs Pre S- proficient =  5.450702e-01
p value t-test for Odor S+ proficient vs Reinf S+ proficient =  5.804979e-01
p value t-test for Odor S- proficient vs Reinf S- proficient =  5.816885e-01
p value t-test for Pre S+ learning vs Reinf S+ learning =  5.843766e-01
p value t-test for Odor S+ learning vs Pre S+ proficient =  6.100572e-01
p value t-test for Baseline S- proficient vs Pre S- proficient =  6.913351e-01
p value t-test for Baseline S+ proficient vs Pre S+ proficient =  6.953289e-01
p value t-test for Pre S+ proficient vs Pre S- proficient =  7.679537e-01
p value t-test for Baseline S+ learning vs Baseline S- learning =  7.944761e-01
p value t-test for Baseline S+ learning vs Baseline S- proficient =  8.629380e-01
p value t-test for Baseline S+ learning vs Pre S+ proficient =  8.725037e-01
p value t-test for Baseline S+ proficient vs Pre S- proficient =  8.755038e-01
p value t-test for Baseline S- proficient vs Pre S+ proficient =  8.804764e-01
p value t-test for Baseline S- learning vs Pre S- proficient =  9.333658e-01
p value t-test for Odor S+ learning vs Reinf S+ learning =  9.856221e-01
p value t-test for Baseline S- learning vs Baseline S+ proficient =  9.910825e-01
p value t-test for Odor S- learning vs Reinf S- learning =  9.944002e-01

glm for bar graph of sum of dPCA stimulus-independent components Figure 3D

mdl =

Generalized linear regression model:
dPCA_component ~ ~ p_correct+time_window+sp_vs_sm+p_correct*time_window*sp_vs_sm
Distribution = Normal

p_correct is learning (1) vs. proficient (3) stages
time_window is baseline (1, -4.5 to -3.5), pre (2, -1 to 0), odor (3, 3.1 to 4.1) and reinforcement (4, 4.5 5.5)
sp_vs_sm is S- (0) and S+ (1)

Estimated Coefficients:

| | Estimate | SE | tStat | pValue |
|---|---|---|---|---|
| (Intercept) | 1.0525 | 4.8316 | 0.21784 | 0.82894 |
| p_correct_3 | -4.2375 | 6.8329 | -0.62016 | 0.53955 |
| time_window_2 | -3.869 | 6.8329 | -0.56623 | 0.57519 |
| time_window_3 | -1.9425 | 6.8329 | -0.28428 | 0.77802 |
| time_window_4 | -0.96877 | 6.8329 | -0.14178 | 0.88814 |
| sp_vs_sm_1 | 0.096838 | 6.8329 | 0.014172 | 0.98878 |
| p_correct_3:time_window_2 | 17.011 | 9.6632 | 1.7604 | 0.087888 |
| p_correct_3:time_window_3 | 6.8814 | 9.6632 | 0.71213 | 0.48155 |
| p_correct_3:time_window_4 | 8.4407 | 9.6632 | 0.87349 | 0.3889 |
| p_correct_3:sp_vs_sm_1 | -0.10047 | 9.6632 | -0.010398 | 0.99177 |
| time_window_2:sp_vs_sm_1 | -0.63048 | 9.6632 | -0.065246 | 0.94838 |
| time_window_3:sp_vs_sm_1 | -0.17396 | 9.6632 | -0.018002 | 0.98575 |
| time_window_4:sp_vs_sm_1 | -0.25118 | 9.6632 | -0.025993 | 0.97942 |
| p_correct_3:time_window_2:sp_vs_sm_1 | 0.64365 | 13.666 | 0.047099 | 0.96273 |
| p_correct_3:time_window_3:sp_vs_sm_1 | 0.16551 | 13.666 | 0.012111 | 0.99041 |
| p_correct_3:time_window_4:sp_vs_sm_1 | 0.28434 | 13.666 | 0.020807 | 0.98353 |

48 observations, 32 error degrees of freedom
Estimated Dispersion: 70
F-statistic vs. constant model: 0.744, p-value = 0.724

Ranksum for dPCA stimulus-independent component

pFDR = 4.166667e-04

p values below are > pFDR

p value t-test for Reinf S+ learning vs Reinf S- proficient = 4.424750e-02
p value t-test for Reinf S+ learning vs Reinf S+ proficient = 4.531953e-02
p value t-test for Reinf S- learning vs Reinf S- proficient = 5.071399e-02
p value t-test for Reinf S- learning vs Reinf S+ proficient = 5.160321e-02
p value t-test for Reinf S+ learning vs Pre S+ proficient = 5.524242e-02
p value t-test for Reinf S+ learning vs Pre S- proficient = 5.541182e-02
p value t-test for Reinf S- learning vs Pre S+ proficient = 5.882487e-02
p value t-test for Reinf S- learning vs Pre S- proficient = 5.902377e-02
p value t-test for Pre S+ proficient vs Reinf S+ proficient = 6.449245e-02
p value t-test for Pre S- proficient vs Reinf S+ proficient = 6.487050e-02
p value t-test for Pre S+ proficient vs Reinf S- proficient = 6.510696e-02
p value t-test for Pre S- proficient vs Reinf S- proficient = 6.547999e-02
p value t-test for Baseline S+ proficient vs Baseline S- proficient = 9.290133e-02
p value t-test for Pre S+ proficient vs Odor S- proficient = 1.007085e-01
p value t-test for Pre S+ proficient vs Odor S+ proficient = 1.011444e-01
p value t-test for Pre S- proficient vs Odor S- proficient = 1.013195e-01
p value t-test for Pre S- proficient vs Odor S+ proficient = 1.017552e-01
p value t-test for Baseline S+ learning vs Baseline S+ proficient = 1.061799e-01
p value t-test for Baseline S+ learning vs Baseline S- proficient = 1.064289e-01
p value t-test for Baseline S- learning vs Baseline S+ proficient = 1.093714e-01
p value t-test for Baseline S- learning vs Baseline S- proficient = 1.096397e-01
p value t-test for Odor S+ learning vs Pre S+ proficient = 1.343127e-01
p value t-test for Odor S+ learning vs Pre S- proficient = 1.346305e-01
p value t-test for Odor S- learning vs Pre S+ proficient = 1.365752e-01
p value t-test for Odor S- learning vs Pre S- proficient = 1.369282e-01
p value t-test for Odor S+ learning vs Reinf S+ proficient = 2.166762e-01
p value t-test for Odor S+ learning vs Reinf S- proficient = 2.168938e-01
p value t-test for Odor S- learning vs Reinf S+ proficient = 2.249486e-01
p value t-test for Odor S- learning vs Reinf S- proficient = 2.254921e-01
p value t-test for Pre S+ learning vs Pre S+ proficient = 2.691859e-01
p value t-test for Pre S+ learning vs Pre S- proficient = 2.695059e-01
p value t-test for Pre S- learning vs Pre S+ proficient = 2.884003e-01
p value t-test for Pre S- learning vs Pre S- proficient = 2.887644e-01
p value t-test for Baseline S+ proficient vs Pre S+ proficient = 2.954530e-01
p value t-test for Baseline S- proficient vs Pre S+ proficient = 2.955742e-01
p value t-test for Baseline S+ proficient vs Pre S- proficient = 2.956559e-01
p value t-test for Baseline S- proficient vs Pre S- proficient = 2.957773e-01
p value t-test for Pre S+ proficient vs Pre S- proficient = 3.023690e-01
p value t-test for Reinf S+ proficient vs Reinf S- proficient = 3.190485e-01
p value t-test for Baseline S- learning vs Pre S+ proficient = 3.782923e-01
p value t-test for Baseline S- learning vs Pre S- proficient = 3.786109e-01
p value t-test for Baseline S+ learning vs Pre S+ proficient = 3.797443e-01
p value t-test for Baseline S+ learning vs Pre S- proficient = 3.800723e-01
p value t-test for Pre S+ learning vs Reinf S+ proficient = 4.032052e-01
p value t-test for Pre S+ learning vs Reinf S- proficient = 4.038042e-01
p value t-test for Baseline S+ learning vs Pre S+ learning = 4.281865e-01
p value t-test for Pre S- learning vs Reinf S+ proficient = 4.377006e-01
p value t-test for Pre S- learning vs Reinf S- proficient = 4.385220e-01
p value t-test for Baseline S- learning vs Pre S+ learning = 4.480038e-01
p value t-test for Baseline S+ proficient vs Reinf S+ proficient = 4.529985e-01
p value t-test for Baseline S- proficient vs Reinf S+ proficient = 4.532082e-01
p value t-test for Baseline S+ proficient vs Reinf S- proficient = 4.534595e-01
p value t-test for Baseline S- proficient vs Reinf S- proficient = 4.536700e-01
p value t-test for Reinf S+ learning vs Reinf S- learning = 4.720805e-01
p value t-test for Odor S- proficient vs Reinf S+ proficient = 4.905724e-01
p value t-test for Odor S+ proficient vs Reinf S+ proficient = 4.906077e-01
p value t-test for Odor S- proficient vs Reinf S- proficient = 4.975020e-01
p value t-test for Odor S+ proficient vs Reinf S- proficient = 4.975053e-01
p value t-test for Odor S+ proficient vs Odor S- proficient = 5.061564e-01
p value t-test for Baseline S+ learning vs Pre S- learning = 5.375315e-01
p value t-test for Baseline S- learning vs Pre S- learning = 5.557084e-01
p value t-test for Baseline S+ learning vs Baseline S- learning = 5.725650e-01
p value t-test for Pre S+ learning vs Pre S- learning = 5.976317e-01

```
p value t-test for Odor S- learning vs Odor S- proficient =  6.279841e-01
p value t-test for Odor S- learning vs Odor S+ proficient =  6.303643e-01
p value t-test for Pre S+ learning vs Odor S- proficient =  6.313030e-01
p value t-test for Pre S+ learning vs Odor S+ proficient =  6.324952e-01
p value t-test for Odor S+ learning vs Odor S- proficient =  6.348847e-01
p value t-test for Odor S+ learning vs Odor S+ proficient =  6.371803e-01
p value t-test for Pre S+ learning vs Odor S- learning =  6.388320e-01
p value t-test for Pre S+ learning vs Reinf S- learning =  6.397961e-01
p value t-test for Pre S+ learning vs Odor S+ learning =  6.404443e-01
p value t-test for Odor S+ learning vs Reinf S- learning =  6.477704e-01
p value t-test for Pre S+ learning vs Reinf S+ learning =  6.562208e-01
p value t-test for Pre S- learning vs Odor S- proficient =  6.606493e-01
p value t-test for Pre S- learning vs Odor S+ proficient =  6.618345e-01
p value t-test for Reinf S+ learning vs Odor S- proficient =  6.715185e-01
p value t-test for Reinf S+ learning vs Odor S+ proficient =  6.746395e-01
p value t-test for Baseline S- learning vs Reinf S+ proficient =  6.770588e-01
p value t-test for Baseline S- learning vs Reinf S- proficient =  6.787235e-01
p value t-test for Odor S- learning vs Reinf S- learning =  6.818863e-01
p value t-test for Baseline S+ learning vs Reinf S+ proficient =  6.825998e-01
p value t-test for Baseline S+ learning vs Reinf S- proficient =  6.843125e-01
p value t-test for Reinf S- learning vs Odor S- proficient =  6.889899e-01
p value t-test for Reinf S- learning vs Odor S+ proficient =  6.921833e-01
p value t-test for Baseline S+ proficient vs Odor S- proficient =  6.944193e-01
p value t-test for Baseline S- proficient vs Odor S- proficient =  6.946321e-01
p value t-test for Baseline S+ proficient vs Odor S+ proficient =  6.954728e-01
p value t-test for Baseline S- proficient vs Odor S+ proficient =  6.956855e-01
p value t-test for Pre S- learning vs Reinf S- learning =  6.957886e-01
p value t-test for Reinf S- learning vs Baseline S+ proficient =  6.986732e-01
p value t-test for Reinf S- learning vs Baseline S- proficient =  6.989933e-01
p value t-test for Odor S+ learning vs Reinf S+ learning =  7.055675e-01
p value t-test for Reinf S+ learning vs Baseline S+ proficient =  7.092198e-01
p value t-test for Reinf S+ learning vs Baseline S- proficient =  7.095465e-01
p value t-test for Pre S- learning vs Odor S- learning =  7.096486e-01
p value t-test for Pre S- learning vs Reinf S+ learning =  7.133680e-01
p value t-test for Baseline S+ learning vs Odor S+ learning =  7.180575e-01
p value t-test for Pre S- learning vs Odor S+ learning =  7.183076e-01
p value t-test for Baseline S- learning vs Odor S+ learning =  7.356989e-01
p value t-test for Odor S- learning vs Reinf S+ learning =  7.415176e-01
p value t-test for Baseline S+ learning vs Odor S- learning =  7.466690e-01
p value t-test for Odor S+ learning vs Baseline S+ proficient =  7.550698e-01
p value t-test for Odor S+ learning vs Baseline S- proficient =  7.554518e-01
p value t-test for Odor S- learning vs Baseline S+ proficient =  7.608311e-01
p value t-test for Odor S- learning vs Baseline S- proficient =  7.611912e-01
p value t-test for Baseline S- learning vs Odor S- learning =  7.628686e-01
p value t-test for Baseline S+ learning vs Reinf S+ learning =  8.544882e-01
p value t-test for Baseline S- learning vs Reinf S+ learning =  8.676567e-01
p value t-test for Baseline S+ learning vs Reinf S- learning =  8.743450e-01
p value t-test for Odor S+ learning vs Odor S- learning =  8.765546e-01
p value t-test for Baseline S- learning vs Reinf S- learning =  8.872132e-01
p value t-test for Baseline S- learning vs Odor S- proficient =  9.486147e-01
p value t-test for Baseline S- learning vs Odor S+ proficient =  9.495755e-01
p value t-test for Pre S- learning vs Baseline S+ proficient =  9.509528e-01
p value t-test for Pre S- learning vs Baseline S- proficient =  9.514241e-01
p value t-test for Baseline S+ learning vs Odor S- proficient =  9.552822e-01
p value t-test for Baseline S+ learning vs Odor S+ proficient =  9.562424e-01
p value t-test for Pre S+ learning vs Baseline S- proficient =  9.740548e-01
p value t-test for Pre S+ learning vs Baseline S+ proficient =  9.746296e-01
```
