## Supplementary material for "Decision-Making Time Cells in Hippocampal Dorsal CA1": Table S5 (from another version of this preprint: version 2, 2023-10-11)

Table S5 glm statistics for Figure 4F

glm for zdFF for cluster 1 peak 1

mdl =

Generalized linear regression model:
    zdFF ~ 1 + p_correct + spm + p_correct*spm
    p_correct is learning (1) vs. proficient (3)
    spm is S+ (1) vs. S- (0)
    Distribution = Normal

Estimated Coefficients:

| | Estimate | SE | tStat | pValue |
|---|---|---|---|---|
| (Intercept) | -0.057777 | 0.079758 | -0.7244 | 0.46914 |
| p_correct_3 | 0.079369 | 0.086633 | 0.91616 | 0.36 |
| spm_1 | 0.33075 | 0.11279 | 2.9323 | 0.003509 |
| p_correct_3:spm_1 | 0.032607 | 0.12252 | 0.26614 | 0.79023 |

538 observations, 534 error degrees of freedom
Estimated Dispersion: 0.261
F-statistic vs. constant model: 22.9, p-value = 5.74e-14

Ranksum or t-test p values for zdFF for cluster 1 peak 1

pFDR = 3.333333e-02

p value ranksum for Proficient S- vs Proficient S+ =  3.707091e-07
p value ranksum for Learning S- vs Proficient S+ =  2.072073e-04
p value ranksum for Learning S- vs Learning S+ =  1.472530e-02
p value ranksum for Learning S+ vs Proficient S- =  3.079188e-02

p values below are > pFDR

p value ranksum for Learning S- vs Proficient S- =  2.224759e-01
p value ranksum for Learning S+ vs Proficient S+ =  3.557845e-01

glm for zdFF for cluster 1 peak 2

mdl =

Generalized linear regression model:
    zdFF ~ 1 + p_correct*spm
    p_correct is learning (1) vs. proficient (3)
    spm is S+ (1) vs. S- (0)
    Distribution = Normal

Estimated Coefficients:

| | Estimate | SE | tStat | pValue |
|---|---|---|---|---|
| (Intercept) | 0.26447 | 0.057839 | 4.5725 | 5.9963e-06 |
| p_correct_3 | 0.13386 | 0.062824 | 2.1306 | 0.033575 |
| spm_1 | -0.15948 | 0.081796 | -1.9497 | 0.051737 |
| p_correct_3:spm_1 | -0.24161 | 0.088847 | -2.7195 | 0.0067519 |

538 observations, 534 error degrees of freedom
Estimated Dispersion: 0.137
F-statistic vs. constant model: 45.9, p-value = 2.19e-26

```
Ranksum or t-test p values for zdFF for cluster 1 peak 2


pFDR = 4.166667e-02

p value ranksum for Proficient S- vs Proficient S+ =  2.864640e-32
p value t-test for Learning S+ vs Proficient S- =  6.920049e-05
p value ranksum for Learning S- vs Proficient S+ =  3.533274e-04
p value t-test for Learning S+ vs Proficient S+ =  9.336759e-03
p value ranksum for Learning S- vs Proficient S- =  2.563431e-02

p values below are > pFDR

p value ranksum for Learning S- vs Learning S+ =  1.453871e-01



glm for zdFF for cluster 2

mdl = 


Generalized linear regression model:
    zdFF ~ 1 + p_correct*spm
    p_correct is learning (1) vs. proficient (3)
    spm is S+ (1) vs. S- (0)
    Distribution = Normal

Estimated Coefficients:
                         Estimate        SE        tStat       pValue  
                         _________    ________    _______    __________

    (Intercept)           -0.18148    0.042287    -4.2916    1.8958e-05
    p_correct_3          -0.055676    0.045074    -1.2352       0.21696
    spm_1                   0.2074    0.059803      3.468    0.00054027
    p_correct_3:spm_1      0.25711    0.063744     4.0335    5.7922e-05


1402 observations, 1398 error degrees of freedom
Estimated Dispersion: 0.15
F-statistic vs. constant model: 153, p-value = 5.67e-86



Ranksum or t-test p values for zdFF for cluster 2


pFDR = 5.000000e-02

p value ranksum for Proficient S- vs Proficient S+ =  5.212662e-88
p value ranksum for Learning S- vs Proficient S+ =  3.000704e-17
p value ranksum for Learning S+ vs Proficient S- =  1.688143e-09
p value ranksum for Learning S- vs Learning S+ =  4.479394e-04
p value ranksum for Learning S+ vs Proficient S+ =  5.748235e-04
p value ranksum for Learning S- vs Proficient S- =  3.887809e-02



glm for zdFF for cluster 3

mdl = 


Generalized linear regression model:
    zdFF ~ 1 + p_correct*spm
    p_correct is learning (1) vs. proficient (3)
    spm is S+ (1) vs. S- (0)
    Distribution = Normal

Estimated Coefficients:
                         Estimate        SE        tStat       pValue  
```

| | | | | |
|---|---|---|---|---|
| (Intercept) | -0.11434 | 0.045782 | -2.4975 | 0.012766 |
| p_correct_3 | 0.14743 | 0.053555 | 2.7528 | 0.0060821 |
| spm_1 | 0.261 | 0.064745 | 4.0312 | 6.2407e-05 |
| p_correct_3:spm_1 | -0.080936 | 0.075739 | -1.0686 | 0.28565 |

624 observations, 620 error degrees of freedom
Estimated Dispersion: 0.176
F-statistic vs. constant model: 15.1, p-value = 1.76e-09

Ranksum or t-test p values for zdFF for cluster 3

pFDR = 3.333333e-02

p value ranksum for Learning S- vs Proficient S+ = 9.277858e-10
p value ranksum for Learning S- vs Proficient S- = 7.319314e-07
p value ranksum for Learning S- vs Learning S+ = 3.879231e-04
p value ranksum for Proficient S- vs Proficient S+ = 4.070510e-04

p values below are > pFDR

p value ranksum for Learning S+ vs Proficient S+ = 8.094633e-02
p value ranksum for Learning S+ vs Proficient S- = 4.806209e-01
