## Supplementary material for "Decision-Making Time Cells in Hippocampal Dorsal CA1": Table S7 (from another version of this preprint: version 2, 2023-10-11)

Table S7 glm statistics for decoding accuracy bar graphs in Figure 6

glm for decoding accuracy, glm fit with data from 0.5 to 5.5 sec (odorant + reinforcement, Figure 6Aiii)

mdl =

Generalized linear regression model:
    accuracy ~ 1 + mice + percent_correct + window + percent_correct*window
    Distribution = Normal

percent_correct is learning (1) vs. proficient (3) stages
time_window is pre (2, -1 to 0), odor (3, 3.1 to 4.1) and reinforcement (4, 4.5 5.5)
mice are the mouse numbers 1-4

Estimated Coefficients:

| | Estimate | SE | tStat | pValue |
|---|---|---|---|---|
| (Intercept) | 0.5415 | 0.026471 | 20.456 | 8.1993e-57 |
| percent_correct_3 | 0.03749 | 0.028579 | 1.3118 | 0.1907 |
| window_2 | 0.082692 | 0.034319 | 2.4095 | 0.016644 |
| window_3 | 0.10454 | 0.034319 | 3.0461 | 0.0025478 |
| mice_2 | -0.065614 | 0.019249 | -3.4087 | 0.00075236 |
| mice_3 | -0.060187 | 0.018633 | -3.2302 | 0.0013904 |
| mice_4 | -0.051928 | 0.020344 | -2.5525 | 0.011246 |
| percent_correct_3:window_2 | 0.16609 | 0.039557 | 4.1988 | 3.6466e-05 |
| percent_correct_3:window_3 | 0.13507 | 0.039557 | 3.4145 | 0.0007373 |

279 observations, 270 error degrees of freedom
Estimated Dispersion: 0.0135
F-statistic vs. constant model: 37.7, p-value = 6.59e-40

Ranksum or t-test p values for decoding accuracy

pFDR = 2.727273e-02

p value t-test for proficient Odor vs Shuffled proficient Reinf =  3.184059e-31
p value t-test for Shuffled proficient Odor vs proficient Odor =  3.284259e-31
p value t-test for Shuffled proficient Pre vs proficient Odor =  5.666732e-31
p value t-test for Shuffled proficient Reinf vs proficient Reinf =  9.691511e-19
p value t-test for Shuffled proficient Odor vs proficient Reinf =  1.015447e-18
p value t-test for Shuffled proficient Pre vs proficient Reinf =  1.426848e-18
p value t-test for proficient Pre vs Shuffled proficient Reinf =  3.439739e-15
p value t-test for proficient Pre vs Shuffled proficient Odor =  3.680389e-15
p value t-test for Shuffled proficient Pre vs proficient Pre =  2.427954e-14
p value t-test for learning Reinf vs Shuffled proficient Reinf =  2.821720e-13
p value t-test for learning Reinf vs Shuffled proficient Odor =  2.885468e-13
p value t-test for learning Reinf vs Shuffled proficient Pre =  1.310711e-12
p value t-test for Shuffled learning Pre vs proficient Odor =  4.166642e-12
p value t-test for Shuffled learning Odor vs proficient Odor =  4.183051e-12
p value t-test for learning Odor vs Shuffled proficient Reinf =  3.436215e-10
p value t-test for learning Odor vs Shuffled proficient Odor =  3.629284e-10
p value t-test for learning Odor vs Shuffled proficient Pre =  1.042849e-09
p value t-test for Shuffled learning Pre vs proficient Reinf =  2.478975e-07
p value t-test for Shuffled learning Odor vs proficient Reinf =  2.546328e-07
p value ranksum for Shuffled learning Reinf vs proficient Odor =  3.511406e-06
p value t-test for Shuffled learning Odor vs proficient Pre =  4.480063e-06
p value t-test for Shuffled learning Pre vs proficient Pre =  4.613952e-06
p value t-test for Shuffled learning Odor vs learning Reinf =  2.828412e-05
p value t-test for Shuffled learning Pre vs learning Reinf =  2.974964e-05
p value t-test for proficient Pre vs proficient Odor =  7.160082e-05
p value ranksum for Shuffled learning Reinf vs proficient Reinf =  1.496217e-04
p value t-test for learning Odor vs Shuffled learning Reinf =  2.483597e-04
p value t-test for Shuffled learning Pre vs learning Odor =  3.799712e-04
p value t-test for Shuffled learning Odor vs learning Odor =  3.800311e-04
p value t-test for learning Pre vs proficient Odor =  5.359495e-04

```
p value t-test for learning Reinf vs proficient Odor =  2.228401e-03
p value t-test for learning Odor vs proficient Odor =  2.466029e-03
p value ranksum for Shuffled learning Reinf vs proficient Pre =  3.229474e-03
p value t-test for proficient Pre vs proficient Reinf =  5.206104e-03
p value t-test for learning Pre vs proficient Reinf =  7.534619e-03
p value ranksum for Shuffled learning Reinf vs learning Reinf =  1.430000e-02

p values below are > pFDR

p value t-test for learning Odor vs proficient Reinf =  4.152076e-02
p value t-test for learning Reinf vs proficient Reinf =  4.202457e-02
p value ranksum for Shuffled learning Reinf vs Shuffled proficient Pre =  7.136014e-02
p value t-test for learning Pre vs Shuffled proficient Pre =  1.035406e-01
p value t-test for learning Pre vs proficient Pre =  1.089238e-01
p value t-test for learning Pre vs Shuffled proficient Odor =  1.168125e-01
p value t-test for learning Pre vs Shuffled proficient Reinf =  1.209863e-01
p value ranksum for Shuffled learning Reinf vs Shuffled proficient Odor =  1.661555e-01
p value t-test for Shuffled learning Odor vs Shuffled learning Reinf =  2.072005e-01
p value t-test for learning Pre vs learning Odor =  2.277739e-01
p value t-test for Shuffled proficient Pre vs Shuffled proficient Reinf =  2.511623e-01
p value t-test for proficient Odor vs proficient Reinf =  2.540390e-01
p value t-test for Shuffled proficient Pre vs Shuffled proficient Odor =  2.929092e-01
p value t-test for learning Pre vs learning Reinf =  2.929630e-01
p value t-test for Shuffled learning Odor vs Shuffled proficient Reinf =  3.146229e-01
p value ranksum for Shuffled learning Reinf vs Shuffled proficient Reinf =  3.384446e-01
p value t-test for learning Pre vs Shuffled learning Odor =  3.547425e-01
p value t-test for Shuffled learning Pre vs learning Pre =  3.651285e-01
p value t-test for learning Pre vs Shuffled learning Reinf =  3.936840e-01
p value t-test for Shuffled learning Odor vs Shuffled proficient Odor =  3.942558e-01
p value t-test for Shuffled learning Pre vs Shuffled learning Reinf =  4.693581e-01
p value t-test for Shuffled learning Pre vs Shuffled proficient Reinf =  5.224103e-01
p value t-test for Shuffled learning Pre vs Shuffled proficient Odor =  6.265247e-01
p value t-test for learning Reinf vs proficient Pre =  6.335521e-01
p value t-test for learning Odor vs proficient Pre =  7.338449e-01
p value t-test for Shuffled proficient Odor vs Shuffled proficient Reinf =  8.073723e-01
p value t-test for Shuffled learning Pre vs Shuffled learning Odor =  8.414302e-01
p value t-test for Shuffled learning Pre vs Shuffled proficient Pre =  8.651150e-01
p value t-test for Shuffled learning Odor vs Shuffled proficient Pre =  9.054246e-01
p value t-test for learning Odor vs learning Reinf =  9.583331e-01



glm for decoding accuracy, glm fit with data from 4.5 to 6.5 sec (reinforcement, Figure 6Biii)

mdl = 


Generalized linear regression model:
    accuracy ~ 1 + mice + percent_correct + window + percent_correct*window
    Distribution = Normal

percent_correct is learning (1) vs. proficient (3) stages
time_window is pre (2, -1 to 0), odor (3, 3.1 to 4.1) and reinforcement (4, 4.5 5.5)
mice are the mouse numbers 1-4

Estimated Coefficients:
                                 Estimate       SE        tStat       pValue  
                                 _________    ________    _______    __________

    (Intercept)                    0.50367    0.023801     21.162    8.4837e-59
    percent_correct_3             0.048511    0.025443     1.9066      0.057657
    window_2                      0.066082    0.030624     2.1578      0.031843
    window_3                       0.18282    0.030624     5.9699    7.6259e-09
    mice_2                        -0.07207    0.016988    -4.2425    3.0618e-05
    mice_3                       -0.055606    0.016778    -3.3142     0.0010476
    mice_4                       -0.059461    0.017652    -3.3684    0.00086882
    percent_correct_3:window_2     0.13912    0.035169     3.9558    9.8052e-05
    percent_correct_3:window_3    0.067644    0.035169     1.9234      0.055507


273 observations, 264 error degrees of freedom
```

Estimated Dispersion: 0.0103
F-statistic vs. constant model: 45, p-value = 4.07e-45

Ranksum or t-test p values for decoding accuracy

pFDR = 2.196970e-02

p value t-test for Shuffled proficient Pre vs proficient Odor =  3.448809e-25
p value t-test for proficient Odor vs Shuffled proficient Reinf =  4.001252e-25
p value t-test for Shuffled proficient Odor vs proficient Odor =  4.342513e-25
p value t-test for Shuffled proficient Pre vs proficient Reinf =  5.672642e-22
p value t-test for Shuffled proficient Reinf vs proficient Reinf =  6.356555e-22
p value t-test for Shuffled proficient Odor vs proficient Reinf =  6.615001e-22
p value t-test for learning Reinf vs Shuffled proficient Pre =  4.430811e-21
p value t-test for learning Reinf vs Shuffled proficient Reinf =  7.069157e-21
p value t-test for learning Reinf vs Shuffled proficient Odor =  1.175668e-20
p value t-test for Shuffled learning Reinf vs proficient Odor =  8.383979e-10
p value t-test for Shuffled learning Odor vs proficient Odor =  1.220174e-09
p value t-test for Shuffled learning Pre vs proficient Odor =  1.472667e-09
p value t-test for Shuffled learning Reinf vs proficient Reinf =  1.447083e-08
p value t-test for Shuffled learning Odor vs proficient Reinf =  2.020951e-08
p value t-test for Shuffled learning Pre vs proficient Reinf =  2.411259e-08
p value t-test for Shuffled learning Reinf vs learning Reinf =  4.179579e-08
p value t-test for Shuffled learning Odor vs learning Reinf =  7.245372e-08
p value t-test for Shuffled learning Pre vs learning Reinf =  8.585902e-08
p value t-test for proficient Pre vs proficient Odor =  9.505129e-04
p value t-test for learning Pre vs proficient Odor =  1.047919e-03
p value t-test for learning Odor vs proficient Odor =  1.080114e-03
p value t-test for proficient Pre vs proficient Reinf =  1.418961e-03
p value t-test for learning Pre vs proficient Reinf =  1.967300e-03
p value t-test for learning Odor vs proficient Reinf =  2.816883e-03
p value t-test for learning Reinf vs proficient Pre =  5.411871e-03
p value t-test for learning Pre vs Shuffled proficient Reinf =  1.207075e-02
p value t-test for learning Pre vs Shuffled proficient Pre =  1.228522e-02
p value t-test for learning Pre vs Shuffled proficient Odor =  1.289198e-02
p value t-test for Shuffled learning Pre vs Shuffled learning Reinf =  2.189770e-02

p values below are > pFDR

p value t-test for Shuffled learning Pre vs Shuffled proficient Pre =  4.018108e-02
p value t-test for learning Odor vs learning Reinf =  4.583095e-02
p value t-test for Shuffled learning Pre vs Shuffled proficient Reinf =  7.053178e-02
p value t-test for learning Pre vs learning Reinf =  7.066225e-02
p value t-test for Shuffled learning Pre vs Shuffled proficient Odor =  7.361261e-02
p value t-test for learning Odor vs Shuffled proficient Pre =  1.027715e-01
p value t-test for learning Reinf vs proficient Odor =  1.045923e-01
p value t-test for learning Odor vs Shuffled proficient Reinf =  1.159864e-01
p value t-test for learning Odor vs Shuffled proficient Odor =  1.163471e-01
p value t-test for learning Reinf vs proficient Reinf =  1.296024e-01
p value t-test for Shuffled learning Pre vs learning Pre =  1.356082e-01
p value t-test for Shuffled learning Reinf vs Shuffled proficient Reinf =  1.403753e-01
p value t-test for Shuffled learning Reinf vs Shuffled proficient Pre =  1.502559e-01
p value t-test for learning Pre vs Shuffled learning Odor =  1.552525e-01
p value t-test for Shuffled learning Odor vs Shuffled learning Reinf =  1.818408e-01
p value t-test for learning Pre vs Shuffled learning Reinf =  1.900061e-01
p value t-test for Shuffled learning Reinf vs Shuffled proficient Odor =  2.196256e-01
p value t-test for learning Odor vs Shuffled learning Reinf =  2.388562e-01
p value t-test for learning Pre vs proficient Pre =  3.247909e-01
p value t-test for Shuffled proficient Pre vs proficient Pre =  3.385000e-01
p value t-test for proficient Pre vs Shuffled proficient Odor =  3.415419e-01
p value t-test for proficient Pre vs Shuffled proficient Reinf =  3.538219e-01
p value t-test for learning Pre vs learning Odor =  3.552929e-01
p value t-test for Shuffled learning Pre vs Shuffled learning Odor =  3.557897e-01
p value t-test for Shuffled learning Odor vs learning Odor =  3.991325e-01
p value t-test for Shuffled learning Reinf vs proficient Pre =  4.806767e-01
p value t-test for Shuffled learning Odor vs Shuffled proficient Pre =  5.111295e-01
p value t-test for Shuffled learning Pre vs learning Odor =  5.143327e-01
p value t-test for Shuffled learning Odor vs Shuffled proficient Odor =  5.495886e-01
p value t-test for Shuffled learning Odor vs Shuffled proficient Reinf =  6.126583e-01

p value t-test for Shuffled learning Odor vs proficient Pre =  6.456169e-01
p value t-test for Shuffled learning Pre vs proficient Pre =  7.556245e-01
p value t-test for proficient Odor vs proficient Reinf =  7.658725e-01
p value t-test for Shuffled proficient Odor vs Shuffled proficient Reinf =  8.492510e-01
p value t-test for Shuffled proficient Pre vs Shuffled proficient Reinf =  8.575122e-01
p value t-test for learning Odor vs proficient Pre =  9.436577e-01
p value t-test for Shuffled proficient Pre vs Shuffled proficient Odor =  9.656179e-01

glm for decoding accuracy, glm fit with data from -5 to -3 sec (before stimulus presentation, Fig. 6Ciii)

mdl =

Generalized linear regression model:
    accuracy ~ 1 + mice + percent_correct + window + percent_correct*window
    Distribution = Normal

percent_correct is learning (1) vs. proficient (3) stages
time_window is pre (2, -1 to 0), odor (3, 3.1 to 4.1) and reinforcement (4, 4.5 5.5)
mice are the mouse numbers 1-4

Estimated Coefficients:

| | Estimate | SE | tStat | pValue |
|---|---|---|---|---|
| (Intercept) | 0.50452 | 0.025776 | 19.573 | 1.0285e-53 |
| percent_correct_3 | -0.0007055 | 0.027549 | -0.025608 | 0.97959 |
| window_2 | -0.0013861 | 0.032973 | -0.042039 | 0.9665 |
| window_3 | -0.00037037 | 0.032973 | -0.011232 | 0.99105 |
| mice_2 | -0.013711 | 0.018533 | -0.73984 | 0.46004 |
| mice_3 | -0.0093454 | 0.018204 | -0.51338 | 0.60811 |
| mice_4 | -0.041229 | 0.019319 | -2.1341 | 0.033737 |
| percent_correct_3:window_2 | 0.011006 | 0.038006 | 0.2896 | 0.77235 |
| percent_correct_3:window_3 | 0.0051901 | 0.038006 | 0.13656 | 0.89148 |

279 observations, 270 error degrees of freedom
Estimated Dispersion: 0.0125
F-statistic vs. constant model: 0.643, p-value = 0.741

Ranksum or t-test p values for decoding accuracy

pFDR = 1.590909e-02

p value t-test for learning Pre vs Shuffled proficient Odor =  2.907989e-07
p value t-test for learning Pre vs Shuffled proficient Reinf =  4.087509e-07
p value t-test for learning Pre vs Shuffled proficient Pre =  4.337883e-07
p value t-test for Shuffled proficient Odor vs proficient Reinf =  1.319734e-06
p value t-test for Shuffled proficient Reinf vs proficient Reinf =  1.915186e-06
p value t-test for Shuffled proficient Pre vs proficient Reinf =  1.925618e-06
p value t-test for Shuffled learning Reinf vs learning Reinf =  5.905390e-05
p value t-test for Shuffled learning Odor vs learning Reinf =  6.790477e-05
p value ranksum for learning Reinf vs Shuffled proficient Pre =  7.541678e-05
p value ranksum for learning Reinf vs Shuffled proficient Odor =  7.541678e-05
p value ranksum for learning Reinf vs Shuffled proficient Reinf =  8.173554e-05
p value ranksum for Shuffled learning Pre vs learning Reinf =  3.432792e-04
p value ranksum for learning Reinf vs proficient Pre =  2.046973e-03
p value t-test for learning Pre vs Shuffled learning Reinf =  2.849841e-03
p value t-test for learning Pre vs Shuffled learning Odor =  3.398594e-03
p value t-test for learning Odor vs Shuffled proficient Odor =  3.620475e-03
p value t-test for learning Odor vs Shuffled proficient Pre =  4.031736e-03
p value t-test for learning Odor vs Shuffled proficient Reinf =  4.093421e-03
p value t-test for Shuffled learning Reinf vs proficient Reinf =  4.319889e-03
p value t-test for Shuffled learning Odor vs proficient Reinf =  5.444356e-03
p value ranksum for Shuffled learning Pre vs learning Pre =  1.430000e-02

p values below are > pFDR

```
p value t-test for learning Pre vs proficient Pre =  4.938983e-02
p value t-test for proficient Pre vs proficient Reinf =  5.490624e-02
p value t-test for Shuffled proficient Odor vs proficient Odor =  9.008523e-02
p value t-test for Shuffled proficient Pre vs proficient Odor =  9.479688e-02
p value t-test for learning Odor vs Shuffled learning Reinf =  9.484538e-02
p value t-test for proficient Odor vs Shuffled proficient Reinf =  9.714226e-02
p value t-test for Shuffled learning Odor vs learning Odor =  1.032793e-01
p value ranksum for learning Reinf vs proficient Reinf =  1.246499e-01
p value ranksum for learning Reinf vs proficient Odor =  2.128852e-01
p value t-test for proficient Pre vs Shuffled proficient Reinf =  2.159505e-01
p value t-test for learning Odor vs proficient Pre =  2.284394e-01
p value ranksum for Shuffled learning Pre vs proficient Reinf =  2.305443e-01
p value t-test for Shuffled proficient Pre vs proficient Pre =  2.365299e-01
p value t-test for proficient Pre vs Shuffled proficient Odor =  2.459458e-01
p value ranksum for Shuffled learning Pre vs proficient Odor =  2.587068e-01
p value t-test for Shuffled learning Reinf vs proficient Odor =  3.241135e-01
p value t-test for Shuffled learning Odor vs proficient Odor =  3.463557e-01
p value ranksum for Shuffled learning Pre vs proficient Pre =  4.141105e-01
p value ranksum for Shuffled learning Pre vs learning Odor =  4.350062e-01
p value t-test for proficient Pre vs proficient Odor =  4.605008e-01
p value t-test for Shuffled learning Odor vs proficient Pre =  4.663918e-01
p value t-test for learning Odor vs learning Reinf =  5.018345e-01
p value t-test for learning Pre vs learning Reinf =  5.182626e-01
p value t-test for Shuffled learning Reinf vs Shuffled proficient Reinf =  5.411468e-01
p value t-test for Shuffled learning Reinf vs proficient Pre =  5.454472e-01
p value t-test for learning Pre vs proficient Odor =  5.629988e-01
p value t-test for Shuffled learning Reinf vs Shuffled proficient Pre =  6.304418e-01
p value t-test for proficient Odor vs proficient Reinf =  6.335666e-01
p value t-test for learning Pre vs proficient Reinf =  6.392362e-01
p value t-test for Shuffled learning Odor vs Shuffled learning Reinf =  6.509631e-01
p value t-test for Shuffled learning Reinf vs Shuffled proficient Odor =  7.043819e-01
p value t-test for Shuffled proficient Odor vs Shuffled proficient Reinf =  7.117857e-01
p value t-test for learning Odor vs proficient Odor =  7.511388e-01
p value ranksum for Shuffled learning Pre vs Shuffled learning Odor =  7.920649e-01
p value ranksum for Shuffled learning Pre vs Shuffled proficient Reinf =  8.134548e-01
p value t-test for Shuffled learning Odor vs Shuffled proficient Odor =  8.207703e-01
p value t-test for learning Pre vs learning Odor =  8.282017e-01
p value t-test for Shuffled proficient Pre vs Shuffled proficient Odor =  8.322217e-01
p value ranksum for Shuffled learning Pre vs Shuffled learning Reinf =  8.604876e-01
p value t-test for Shuffled proficient Pre vs Shuffled proficient Reinf =  8.869032e-01
p value t-test for learning Odor vs proficient Reinf =  9.261596e-01
p value t-test for Shuffled learning Odor vs Shuffled proficient Pre =  9.416772e-01
p value ranksum for Shuffled learning Pre vs Shuffled proficient Odor =  9.467533e-01
p value t-test for Shuffled learning Odor vs Shuffled proficient Reinf =  9.753617e-01
p value ranksum for Shuffled learning Pre vs Shuffled proficient Pre =  9.893431e-01


glm for decoding accuracy, glm fit with data from 2 to 4 sec (narrow window at end of odorant period)

mdl =


Generalized linear regression model:
    accuracy ~ 1 + mice + percent_correct + window + percent_correct*window
    Distribution = Normal

percent_correct is learning (1) vs. proficient (3) stages
time_window is pre (2, -1 to 0), odor (3, 3.1 to 4.1) and reinforcement (4, 4.5 5.5)
mice are the mouse numbers 1-4

Estimated Coefficients:
                                  Estimate       SE        tStat       pValue  
                                 _________    ________    _______    __________

    (Intercept)                    0.53337    0.024795     21.511    5.4861e-60
    percent_correct_3             0.036253    0.026507     1.3677       0.17257
    window_2                        0.1099    0.031904     3.4448    0.00066464
    window_3                       0.07623    0.031904     2.3894       0.01758
    mice_2                       -0.063292    0.017698    -3.5763    0.00041437
    mice_3                       -0.068924    0.017479    -3.9433    0.00010304
    mice_4                       -0.047125     0.01839    -2.5625      0.010946
```

```
    percent_correct_3:window_2      0.10519    0.036639     2.8709     0.0044246
    percent_correct_3:window_3     0.099434    0.036639     2.7139     0.0070876


273 observations, 264 error degrees of freedom
Estimated Dispersion: 0.0112
F-statistic vs. constant model: 30.4, p-value = 1.62e-33



Ranksum or t-test p values for decoding accuracy


pFDR = 2.424242e-02

p value t-test for Shuffled proficient Odor vs proficient Reinf =  1.655313e-18
p value t-test for Shuffled proficient Odor vs proficient Odor =  1.708193e-18
p value t-test for Shuffled proficient Reinf vs proficient Reinf =  2.144025e-18
p value t-test for proficient Odor vs Shuffled proficient Reinf =  2.177340e-18
p value t-test for proficient Pre vs Shuffled proficient Odor =  3.511454e-09
p value t-test for proficient Pre vs Shuffled proficient Reinf =  6.050248e-09
p value t-test for learning Pre vs Shuffled proficient Odor =  9.711150e-08
p value t-test for learning Pre vs Shuffled proficient Pre =  1.050545e-07
p value t-test for learning Pre vs Shuffled proficient Reinf =  1.260368e-07
p value t-test for Shuffled learning Reinf vs proficient Reinf =  3.271633e-07
p value t-test for Shuffled learning Reinf vs proficient Odor =  3.395932e-07
p value t-test for Shuffled learning Pre vs proficient Reinf =  3.862306e-07
p value t-test for Shuffled learning Pre vs proficient Odor =  3.933518e-07
p value ranksum for Shuffled proficient Pre vs proficient Odor =  4.956415e-06
p value t-test for learning Odor vs Shuffled proficient Odor =  1.742121e-05
p value t-test for learning Odor vs Shuffled proficient Pre =  1.888734e-05
p value t-test for learning Odor vs Shuffled proficient Reinf =  2.378122e-05
p value ranksum for Shuffled learning Odor vs proficient Odor =  7.943243e-05
p value ranksum for Shuffled proficient Pre vs proficient Reinf =  2.496630e-04
p value t-test for Shuffled learning Reinf vs proficient Pre =  4.455716e-04
p value t-test for Shuffled learning Pre vs proficient Pre =  7.496707e-04
p value ranksum for Shuffled learning Odor vs proficient Reinf =  1.614463e-03
p value t-test for proficient Pre vs proficient Odor =  2.011126e-03
p value t-test for proficient Pre vs proficient Reinf =  2.106915e-03
p value t-test for learning Pre vs Shuffled learning Odor =  2.317308e-03
p value t-test for learning Pre vs Shuffled learning Reinf =  2.586015e-03
p value t-test for Shuffled learning Pre vs learning Pre =  3.078009e-03
p value ranksum for Shuffled proficient Pre vs proficient Pre =  5.805191e-03
p value t-test for learning Odor vs Shuffled learning Reinf =  1.395805e-02
p value ranksum for Shuffled learning Odor vs learning Odor =  1.486989e-02
p value ranksum for Shuffled learning Odor vs proficient Pre =  1.655278e-02
p value t-test for Shuffled learning Pre vs learning Odor =  1.745350e-02

p values below are > pFDR

p value t-test for learning Odor vs proficient Odor =  3.293729e-02
p value t-test for learning Odor vs proficient Reinf =  4.040403e-02
p value ranksum for learning Reinf vs proficient Odor =  1.024564e-01
p value ranksum for learning Reinf vs proficient Reinf =  1.024564e-01
p value t-test for learning Pre vs proficient Odor =  1.063706e-01
p value t-test for learning Pre vs proficient Reinf =  1.407642e-01
p value t-test for Shuffled learning Reinf vs learning Reinf =  1.665393e-01
p value t-test for Shuffled learning Pre vs learning Reinf =  1.777782e-01
p value ranksum for learning Reinf vs proficient Pre =  1.797027e-01
p value t-test for learning Pre vs proficient Pre =  2.788060e-01
p value t-test for proficient Odor vs proficient Reinf =  3.047191e-01
p value t-test for Shuffled learning Pre vs Shuffled learning Odor =  3.151132e-01
p value t-test for Shuffled proficient Odor vs Shuffled proficient Reinf =  3.172228e-01
p value t-test for Shuffled learning Reinf vs Shuffled proficient Reinf =  4.060007e-01
p value ranksum for Shuffled learning Odor vs Shuffled proficient Reinf =  4.782475e-01
p value ranksum for Shuffled proficient Pre vs Shuffled proficient Odor =  4.796446e-01
p value t-test for Shuffled learning Reinf vs Shuffled proficient Pre =  4.832350e-01
p value t-test for Shuffled learning Pre vs Shuffled proficient Odor =  5.378240e-01
p value t-test for learning Pre vs learning Odor =  5.486662e-01
p value t-test for Shuffled learning Pre vs Shuffled learning Reinf =  5.880344e-01
p value t-test for learning Pre vs learning Reinf =  7.219234e-01
p value ranksum for Shuffled proficient Pre vs Shuffled proficient Reinf =  7.269550e-01
```

```
p value t-test for learning Odor vs proficient Pre =  7.878599e-01
p value ranksum for learning Reinf vs Shuffled proficient Pre =  8.071421e-01
p value ranksum for learning Reinf vs Shuffled proficient Reinf =  8.071421e-01
p value ranksum for Shuffled learning Odor vs Shuffled proficient Pre =  8.711342e-01
p value t-test for Shuffled learning Pre vs Shuffled proficient Pre =  9.018900e-01
p value t-test for learning Odor vs learning Reinf =  9.204761e-01
p value ranksum for learning Reinf vs Shuffled proficient Odor =  9.299711e-01
p value ranksum for Shuffled learning Odor vs learning Reinf =  9.330693e-01
p value t-test for Shuffled learning Reinf vs Shuffled proficient Odor =  9.488335e-01
p value t-test for Shuffled learning Pre vs Shuffled proficient Reinf =  9.615502e-01
p value ranksum for Shuffled learning Odor vs Shuffled learning Reinf =  9.719131e-01
p value ranksum for Shuffled learning Odor vs Shuffled proficient Odor =  9.963020e-01
```
