## Supplementary material for "Decision-Making Time Cells in Hippocampal Dorsal CA1": Table S8 (from another version of this preprint: version 2, 2023-10-11)

Table S8 glm statistics for Figures 7C and F

glm prediction for S+ and S- trials, Figure 7C

mdl =

Generalized linear regression model:
prediction ~ 1 + mice + sp_vs_sm + time_interval + sp_vs_sm*time_interval
Distribution = Normal

time_interval is baseline (1, -2.5 to -1.5) and odor (2, 2 to 4.1)
sp_vs_sm is S- (0) and S+ (1)
mice is the mouse number (1-4)

Estimated Coefficients:

| | Estimate | SE | tStat | pValue |
|---|---|---|---|---|
| (Intercept) | 0.30064 | 0.018616 | 16.149 | 7.706e-39 |
| sp_vs_sm_1 | 0.031375 | 0.021958 | 1.4288 | 0.15452 |
| time_interval_1 | -0.10358 | 0.021958 | -4.717 | 4.3276e-06 |
| mice_2 | 0.01811 | 0.019311 | 0.93779 | 0.34941 |
| mice_3 | -0.087965 | 0.021024 | -4.1841 | 4.1856e-05 |
| mice_4 | -0.070229 | 0.025285 | -2.7775 | 0.0059658 |
| sp_vs_sm_1:time_interval_1 | 0.64098 | 0.031054 | 20.641 | 1.0184e-52 |

220 observations, 213 error degrees of freedom
Estimated Dispersion: 0.0133
F-statistic vs. constant model: 194, p-value = 1.69e-83

Ranksum or t-test p values for prediction for S+ and S- trials

pFDR = 5.000000e-02

p value t-test for S- odor vs S+ odor = 1.754638e-38
p value t-test for S+ base vs S+ odor = 6.162267e-28
p value t-test for S- base vs S+ odor = 8.259852e-28
p value t-test for S+ base vs S- odor = 1.002091e-08
p value t-test for S- base vs S- odor = 4.979250e-06
p value t-test for S- base vs S+ base = 4.891135e-02

glm prediction for between trials vs. shuffled Figure 7D

mdl =

Generalized linear regression model:
prediction ~ 1 + shuffled + mice
Distribution = Normal

shuffled is between (0) and shuffled (1)
mice is the mouse number (1-4)

Estimated Coefficients:

| | Estimate | SE | tStat | pValue |
|---|---|---|---|---|
| (Intercept) | 0.22527 | 0.017388 | 12.955 | 1.6603e-23 |
| shuffled_1 | 0.29513 | 0.017958 | 16.434 | 8.5363e-31 |
| mice_2 | 0.0093926 | 0.022335 | 0.42053 | 0.67496 |
| mice_3 | -0.069813 | 0.024316 | -2.8711 | 0.0049481 |
| mice_4 | 0.018739 | 0.029244 | 0.6408 | 0.52305 |

110 observations, 105 error degrees of freedom
Estimated Dispersion: 0.00887
F-statistic vs. constant model: 70.7, p-value = 6.32e-29

Ranksum or t-test p values for prediction between vs. shuffled

pFDR = 5.000000e-02

p value t-test for Between vs Between shuffled =  2.583595e-20

glm prediction for spontaneous shifts vs. no change in Figure 7F

mdl =

Generalized linear regression model:
    prediction ~ 1 + mice + spont_vs_no_change*time_interval
    Distribution = Normal

time_interval is baseline (1, -2 to -0.5) and odor (2, 0 to 0.2)
sp_vs_sm is S- (0) and S+ (1)
mice is the mouse number (1-4)

Estimated Coefficients:

| | Estimate | SE | tStat | pValue |
|---|---|---|---|---|
| (Intercept) | 0.038782 | 0.015382 | 2.5213 | 0.012536 |
| spont_vs_no_change_1 | 0.15727 | 0.017672 | 8.8992 | 5.1201e-16 |
| time_interval_1 | 0.0050877 | 0.018618 | 0.27327 | 0.78495 |
| mice_2 | 0.034426 | 0.0162 | 2.125 | 0.034913 |
| mice_3 | -0.011105 | 0.016133 | -0.6883 | 0.49212 |
| mice_4 | -0.010901 | 0.019015 | -0.57327 | 0.56715 |
| spont_vs_no_change_1:time_interval_1 | 0.77325 | 0.024824 | 31.149 | 1.6589e-75 |

192 observations, 185 error degrees of freedom
Estimated Dispersion: 0.00728
F-statistic vs. constant model: 703, p-value = 1.94e-124

Ranksum or t-test p values for prediction for between trials

pFDR = 5.000000e-02

p value t-test for Spontaneous base vs Spontaneous onset =  1.620129e-61
p value ranksum for No change base vs Spontaneous onset =  4.060890e-17
p value ranksum for No change onset vs Spontaneous onset =  1.972160e-16
p value ranksum for No change base vs Spontaneous base =  8.066333e-15
p value t-test for Spontaneous base vs No change onset =  1.989703e-09
p value ranksum for No change base vs No change onset =  2.466061e-03

glm for correlation Figure 7G

mdl =

Generalized linear regression model:
    rho ~ 1 + mice + time_window + sp_vs_sm +time_window*sp_vs_sm
    Distribution = Normal

time_window is baseline (1, -2.5 to -1.5) and odor (2, 2 to 4.1)
sp_vs_sm is S- (0) and S+ (1)
mice is the mouse number (1-4)

Estimated Coefficients:

| | Estimate | SE | tStat | pValue |
|---|---|---|---|---|

| | | | | |
|---|---|---|---|---|
| (Intercept) | 0.038744 | 0.0034184 | 11.334 | 1.5963e-23 |
| time_window_1 | -0.010752 | 0.0040497 | -2.6551 | 0.0085389 |
| sp_vs_sm_1 | 0.0031345 | 0.0040497 | 0.77401 | 0.4398 |
| mice_2 | -0.015862 | 0.003529 | -4.4946 | 1.1537e-05 |
| mice_3 | -0.018398 | 0.0039495 | -4.6583 | 5.6683e-06 |
| mice_4 | -0.0017083 | 0.0046206 | -0.36972 | 0.71197 |
| time_window_1:sp_vs_sm_1 | 0.027952 | 0.0057272 | 4.8806 | 2.0964e-06 |

216 observations, 209 error degrees of freedom
Estimated Dispersion: 0.000443
F-statistic vs. constant model: 15.7, p-value = 7.09e-15

Ranksum or t-test p values for correlation

pFDR = 4.166667e-02

p value ranksum for S- odor vs S+ odor =  2.422162e-10
p value t-test for S+ baseline vs S- odor =  8.500401e-05
p value t-test for S- baseline vs S+ odor =  2.195161e-04
p value t-test for S+ baseline vs S+ odor =  7.790802e-04
p value t-test for S- baseline vs S- odor =  1.983722e-03

p values below are > pFDR

p value t-test for S- baseline vs S+ baseline =  1.626560e-01
