## Supplementary material for "Decision-Making Time Cells in Hippocampal Dorsal CA1": Table S9 (from another version of this preprint: version 2, 2023-10-11)

Table S9 glm statistics for Figures 8A, B and C

glm for decoding accuracy (Figures 8A and B)

mdl =

Generalized linear regression model:
accuracy ~ 1 + number_of_ROIs+time_window +number_of_ROIs*time_window
Distribution = Normal

number_of_ROIs: 1: 1 ROI, 2: 2 ROIs, 3: 5 ROIs, 4: 15 ROIs, 5: all ROIs
time_window: 0: Pre, 1: Odor

Estimated Coefficients:

| | Estimate | SE | tStat | pValue |
|---|---|---|---|---|
| (Intercept) | 0.34943 | 0.0022253 | 157.03 | 0 |
| number_of_ROIs_2 | 0.065771 | 0.0044325 | 14.838 | 1.2905e-49 |
| number_of_ROIs_3 | 0.18946 | 0.0044122 | 42.939 | 0 |
| number_of_ROIs_4 | 0.31902 | 0.0043686 | 73.025 | 0 |
| number_of_ROIs_5 | 0.44718 | 0.023643 | 18.914 | 2.7889e-79 |
| time_window_1 | 0.042685 | 0.003147 | 13.564 | 8.9703e-42 |
| number_of_ROIs_2:time_window_1 | -0.031088 | 0.0062685 | -4.9593 | 7.1166e-07 |
| number_of_ROIs_3:time_window_1 | -0.10962 | 0.0062398 | -17.568 | 1.0251e-68 |
| number_of_ROIs_4:time_window_1 | -0.20359 | 0.0061782 | -32.953 | 1.4349e-233 |
| number_of_ROIs_5:time_window_1 | -0.29452 | 0.033436 | -8.8085 | 1.3415e-18 |

27354 observations, 27344 error degrees of freedom
Estimated Dispersion: 0.0332
F-statistic vs. constant model: 798, p-value = 0

glm for divergence times (Figure 8C)

mdl =

Generalized linear regression model:
accuracy ~ 1 + number_of_ROIs
Distribution = Normal

number_of_ROIs: 1: 1 ROI, 2: 2 ROIs, 3: 5 ROIs, 4: 15 ROIs, 5: all ROIs

Estimated Coefficients:

| | Estimate | SE | tStat | pValue |
|---|---|---|---|---|
| (Intercept) | 0.37077 | 0.0016108 | 230.18 | 0 |
| number_of_ROIs_2 | 0.050228 | 0.0032085 | 15.655 | 5.3643e-55 |
| number_of_ROIs_3 | 0.13465 | 0.0031938 | 42.158 | 0 |
| number_of_ROIs_4 | 0.21722 | 0.0031623 | 68.692 | 0 |
| number_of_ROIs_5 | 0.29992 | 0.017114 | 17.525 | 2.1746e-68 |

27354 observations, 27349 error degrees of freedom
Estimated Dispersion: 0.0348
F-statistic vs. constant model: 1.4e+03, p-value = 0
