## Supplementary material for "Decision-Making Time Cells in Hippocampal Dorsal CA1": Table S10 (from another version of this preprint: version 2, 2023-10-11)

Table S10 glm statistics for Figures 9B, 9D and 9F

glm for lick fraction, Figure 9B

mdl =

Generalized linear regression model:
    lick_f ~ 1 + mice + window + trial_type + window*trial_type
    Distribution = Normal

lick_f is lick fraction
window is RA1 (1, 0 to 2 sec) and RA2 (2, 2 to 4 sec)
trial type is fa (0), CR (1), miss (2) and hit (3)
mice is the mouse number (1-4)

Estimated Coefficients:

| | Estimate | SE | tStat | pValue |
|---|---|---|---|---|
| (Intercept) | 0.07034 | 0.011801 | 5.9607 | 5.1481e-09 |
| window_2 | -0.019551 | 0.014663 | -1.3334 | 0.18309 |
| trial_type_1 | -0.081351 | 0.01377 | -5.9077 | 6.9432e-09 |
| trial_type_2 | -0.087379 | 0.016863 | -5.1817 | 3.3534e-07 |
| trial_type_3 | -0.0030771 | 0.01377 | -0.22346 | 0.82328 |
| mice_2 | 0.060318 | 0.0089129 | 6.7675 | 4.1855e-11 |
| mice_3 | 0.036513 | 0.010003 | 3.6502 | 0.00029343 |
| mice_4 | 0.057122 | 0.01105 | 5.1696 | 3.5632e-07 |
| window_2:trial_type_1 | -0.0023612 | 0.019436 | -0.12148 | 0.90336 |
| window_2:trial_type_2 | 0.086898 | 0.02367 | 3.6712 | 0.00027103 |
| window_2:trial_type_3 | 0.1114 | 0.019436 | 5.7314 | 1.8486e-08 |

452 observations, 441 error degrees of freedom
Estimated Dispersion: 0.0057
F-statistic vs. constant model: 36.7, p-value = 4.52e-52

Ranksum or t-test p values for decoding lick fraction

pFDR = 3.750000e-02

p value ranksum for CR RA2 vs Hit RA2 =  3.704094e-21
p value ranksum for FA RA2 vs CR RA2 =  9.038216e-20
p value t-test for Hit RA1 vs CR RA2 =  1.429718e-19
p value ranksum for FA RA1 vs CR RA2 =  4.069942e-19
p value ranksum for CR RA1 vs Hit RA2 =  3.237006e-16
p value ranksum for CR RA1 vs CR RA2 =  9.243886e-14
p value t-test for Hit RA1 vs Hit RA2 =  2.536200e-13
p value ranksum for Miss RA1 vs Hit RA2 =  7.040433e-13
p value ranksum for CR RA1 vs Hit RA1 =  7.743892e-12
p value ranksum for FA RA1 vs CR RA1 =  1.106009e-11
p value ranksum for Miss RA1 vs Hit RA1 =  3.457106e-11
p value ranksum for FA RA1 vs Miss RA1 =  8.026909e-11
p value ranksum for Miss RA1 vs FA RA2 =  6.863695e-10
p value ranksum for CR RA1 vs FA RA2 =  5.093459e-08
p value ranksum for FA RA2 vs Hit RA2 =  7.654723e-07
p value ranksum for CR RA2 vs Miss RA2 =  9.783260e-07
p value ranksum for Miss RA2 vs Hit RA2 =  5.378109e-06
p value ranksum for FA RA1 vs Hit RA2 =  7.794340e-05
p value ranksum for CR RA1 vs Miss RA1 =  1.158677e-04
p value ranksum for Miss RA1 vs Miss RA2 =  5.959864e-04
p value ranksum for FA RA1 vs Miss RA2 =  2.951126e-02

p values below are > pFDR

p value ranksum for FA RA1 vs FA RA2 =  7.222818e-02
p value ranksum for FA RA2 vs Miss RA2 =  8.167497e-02
p value t-test for Hit RA1 vs Miss RA2 =  1.571962e-01

p value ranksum for CR RA1 vs Miss RA2 =  3.065255e-01
p value t-test for Hit RA1 vs FA RA2 =  3.588729e-01
p value ranksum for FA RA1 vs Hit RA1 =  4.824948e-01
p value ranksum for Miss RA1 vs CR RA2 =  6.122093e-01

glm for prediction for lick decoding Figure 9D

mdl =

Generalized linear regression model:
    prediction ~ 1 + mice + window + trial_type + window*trial_type
    Distribution = Normal

window is RA1 (1, 0 to 2 sec) and RA2 (2, 2 to 4 sec)
trial type is fa (0), CR (1), miss (2) and hit (3)
mice is the mouse number (1-4)

Estimated Coefficients:

| | Estimate | SE | tStat | pValue |
|---|---|---|---|---|
| (Intercept) | 0.11114 | 0.012522 | 8.8756 | 1.4106e-17 |
| window_2 | -0.014374 | 0.014524 | -0.98965 | 0.32285 |
| trial_type_1 | -0.013913 | 0.014524 | -0.95792 | 0.33859 |
| trial_type_2 | -0.042086 | 0.015114 | -2.7846 | 0.0055723 |
| trial_type_3 | 0.039482 | 0.014524 | 2.7184 | 0.0067989 |
| mice_2 | -0.06102 | 0.010082 | -6.0524 | 2.8904e-09 |
| mice_3 | 0.0081111 | 0.010373 | 0.78192 | 0.43465 |
| mice_4 | 0.028315 | 0.013646 | 2.0749 | 0.038533 |
| window_2:trial_type_1 | -0.013238 | 0.02054 | -0.64452 | 0.51955 |
| window_2:trial_type_2 | 0.033767 | 0.021274 | 1.5873 | 0.11312 |
| window_2:trial_type_3 | 0.074756 | 0.02054 | 3.6396 | 0.00030291 |

488 observations, 477 error degrees of freedom
Estimated Dispersion: 0.00664
F-statistic vs. constant model: 24.3, p-value = 4.52e-37

Ranksum or t-test p values for prediction of lick fraction

pFDR = 3.571429e-02

p value t-test for CR RA2 vs Hit RA2 =  3.474436e-16
p value t-test for Hit RA1 vs CR RA2 =  1.551539e-14
p value ranksum for Miss RA1 vs Hit RA2 =  1.717194e-12
p value ranksum for Miss RA1 vs Hit RA1 =  4.176381e-10
p value ranksum for Miss RA2 vs Hit RA2 =  1.454843e-09
p value ranksum for CR RA1 vs Hit RA2 =  5.056222e-09
p value t-test for Hit RA1 vs Hit RA2 =  1.251441e-08
p value ranksum for FA RA2 vs Hit RA2 =  1.730618e-08
p value ranksum for FA RA1 vs Hit RA2 =  4.985877e-07
p value t-test for Hit RA1 vs FA RA2 =  1.531499e-06
p value ranksum for CR RA1 vs Miss RA1 =  2.570391e-06
p value ranksum for CR RA1 vs Hit RA1 =  4.917732e-05
p value t-test for Hit RA1 vs Miss RA2 =  6.014429e-05
p value ranksum for Miss RA1 vs CR RA2 =  2.536787e-04
p value ranksum for FA RA1 vs Hit RA1 =  6.400447e-04
p value ranksum for CR RA1 vs Miss RA2 =  7.517950e-04
p value ranksum for FA RA1 vs Miss RA1 =  1.458223e-03
p value ranksum for Miss RA1 vs FA RA2 =  6.102703e-03
p value ranksum for CR RA1 vs CR RA2 =  1.071574e-02
p value ranksum for FA RA1 vs Miss RA2 =  3.153581e-02

p values below are > pFDR

p value ranksum for FA RA2 vs Miss RA2 =  8.183068e-02
p value ranksum for CR RA1 vs FA RA2 =  1.757823e-01
p value ranksum for FA RA1 vs CR RA1 =  2.782358e-01
p value t-test for CR RA2 vs Miss RA2 =  3.584036e-01
p value ranksum for Miss RA1 vs Miss RA2 =  4.169846e-01
p value ranksum for FA RA1 vs FA RA2 =  7.155994e-01
p value ranksum for FA RA1 vs CR RA2 =  8.757761e-01
p value ranksum for FA RA2 vs CR RA2 =  8.872538e-01

glm for prediction for stimulus decoding Figure 9F

mdl =

Generalized linear regression model:
    prediction ~ 1 + mice + window + trial_type + window*trial_type
    Distribution = Normal

window is RA1 (1, 0 to 2 sec) and RA2 (2, 2 to 4 sec)
trial type is fa (0), CR (1), miss (2) and hit (3)
mice is the mouse number (1-4)

Estimated Coefficients:

| | Estimate | SE | tStat | pValue |
|---|---|---|---|---|
| (Intercept) | 0.35796 | 0.030663 | 11.674 | 1.2669e-27 |
| window_2 | 0.011815 | 0.0381 | 0.31011 | 0.75662 |
| trial_type_1 | -0.071999 | 0.035781 | -2.0122 | 0.044806 |
| trial_type_2 | 0.28348 | 0.043817 | 6.4695 | 2.618e-10 |
| trial_type_3 | 0.36959 | 0.035781 | 10.329 | 1.5224e-22 |
| mice_2 | -0.034848 | 0.023159 | -1.5047 | 0.13311 |
| mice_3 | 0.039824 | 0.025992 | 1.5321 | 0.12621 |
| mice_4 | -0.013511 | 0.028711 | -0.47059 | 0.63817 |
| window_2:trial_type_1 | -0.070644 | 0.050504 | -1.3988 | 0.16258 |
| window_2:trial_type_2 | 0.0091297 | 0.061505 | 0.14844 | 0.88206 |
| window_2:trial_type_3 | 0.060556 | 0.050504 | 1.199 | 0.23116 |

452 observations, 441 error degrees of freedom
Estimated Dispersion: 0.0385
F-statistic vs. constant model: 57.3, p-value = 2.16e-73

Ranksum or t-test p values for prediction of stimulus

pFDR = 4.107143e-02

p value t-test for CR RA1 vs Hit RA2 =  7.236846e-29
p value t-test for CR RA1 vs Hit RA1 =  3.175462e-27
p value t-test for Hit RA1 vs CR RA2 =  7.646617e-27
p value ranksum for CR RA2 vs Hit RA2 =  8.432984e-24
p value t-test for Hit RA1 vs FA RA2 =  1.509209e-17
p value t-test for Miss RA1 vs CR RA2 =  8.177990e-17
p value t-test for CR RA1 vs Miss RA1 =  2.846288e-15
p value t-test for CR RA1 vs Miss RA2 =  4.386624e-15
p value ranksum for FA RA2 vs Hit RA2 =  3.147908e-14
p value ranksum for FA RA1 vs Hit RA2 =  7.745631e-14
p value ranksum for FA RA1 vs Hit RA1 =  1.537192e-12
p value ranksum for CR RA2 vs Miss RA2 =  1.479269e-09
p value t-test for Hit RA1 vs Hit RA2 =  9.402201e-07
p value t-test for Miss RA1 vs FA RA2 =  1.993430e-05
p value ranksum for FA RA1 vs Miss RA2 =  3.773171e-05
p value ranksum for FA RA1 vs Miss RA1 =  4.080473e-05
p value ranksum for FA RA2 vs Miss RA2 =  5.108977e-05
p value t-test for CR RA1 vs CR RA2 =  6.651804e-05

```
p value t-test for Miss RA1 vs Hit RA2 =  1.226457e-04
p value ranksum for FA RA2 vs CR RA2 =  1.070743e-03
p value ranksum for FA RA1 vs CR RA2 =  8.595952e-03
p value t-test for CR RA1 vs FA RA2 =  1.331064e-02
p value t-test for Miss RA1 vs Hit RA1 =  2.668990e-02

p values below are > pFDR

p value ranksum for Miss RA2 vs Hit RA2 =  1.025200e-01
p value t-test for Hit RA1 vs Miss RA2 =  1.118235e-01
p value ranksum for FA RA1 vs CR RA1 =  4.825188e-01
p value t-test for Miss RA1 vs Miss RA2 =  6.804136e-01
p value ranksum for FA RA1 vs FA RA2 =  6.859160e-01
```
